# Phylobone: A comprehensive database of bone extracellular matrix proteins in human and model organisms

**DOI:** 10.1101/2023.03.28.534628

**Authors:** Margalida Fontcuberta-Rigo, Miho Nakamura, Pere Puigbò

## Abstract

The bone extracellular matrix (ECM) contains minerals deposited on highly crosslinked collagen fibrils and hundreds of non-collagenous proteins. Some of these proteins are key to the regulation of bone formation and regeneration *via* signaling pathways, and play important regulatory and structural roles. However, the complete list of bone extracellular matrix proteins, their roles, and the extent of individual and cross-species variations have not been fully captured in both humans and model organisms. Here, we introduce the most comprehensive resource of bone extracellular matrix (ECM) proteins that can be used in research fields such as bone regeneration, osteoporosis, and mechanobiology. The Phylobone database (available at https://phylobone.com) includes 255 proteins potentially expressed in the bone extracellular matrix (ECM) of humans and 30 species of vertebrates. A bioinformatics pipeline was used to identify the evolutionary relationships of bone ECM proteins. The analysis facilitated the identification of potential model organisms to study the molecular mechanisms of bone regeneration. A network analysis showed high connectivity of bone ECM proteins. A total of 214 functional protein domains were identified, including collagen and the domains involved in bone formation and resorption. Information from public drug repositories was used to identify potential repurposing of existing drugs. The Phylobone database provides a platform to study bone regeneration and osteoporosis in light of (biological) evolution, and will substantially contribute to the identification of molecular mechanisms and drug targets.

## INTRODUCTION

The bone extracellular matrix (ECM) consists of both organic compounds, which make up approximately 40% of the matrix, and inorganic compounds, which account for the remaining 60% ^1^. The organic fraction of the ECM is mostly composed of collagen (90%) and hundreds of non-collagenous proteins ^2^ that play structural and regulatory roles ^3,4^. Liquid chromatography tandem mass spectrometry (LC-MS-MS) proteomics has eased the elucidation of regulatory [.1] mechanisms, therapeutic strategies, and biomarkers for bone regeneration and osteoporosis research ^5^. Several studies of bone proteomics have reported using secreted proteins from cells, including mesenchymal stem cells, osteoblasts, and osteoclasts ^6^. However, since the bone ECM contains minerals deposited on highly crosslinked collagen fibrils, it is highly challenging to solubilize for proteomics analyses ^7^. Usually, a combination of decalcification and chemical treatments is necessary for the extraction of proteins and LC-MS-MS-based proteomics ^8^. Thus, the availability of bioinformatics resources is necessary to facilitate comparative studies of bone extracellular matrix proteomes, both within human populations and across different model organisms. Although several non-collagenous proteins of the bone ECM proteome have been identified, the list and roles of most proteins ^1^, as well as individual and cross-species variations, have not been fully explicated. Therefore, the identification of non-collagenous proteins in humans and model organisms will significantly advance the field of bone regeneration and osteoporosis.

In this article, we introduce the most comprehensive [.1] database of putative bone ECM proteins in 39 species, including *Homo sapiens* and the most common animal models (e.g., *Danio rerio, Mus musculus, and Xenopus laevis*), useful for the study of osteoporosis ^9^. Due to deer antlers being a speculated model of bone regeneration and osteoporosis ^10–14^, the database includes proteins from six species of the Cerevidae family. The Phylobone database, which includes 255 (28 collagenous and 227 non-collagenous) proteins, presents information on protein sequences, functional characterizations, and potential drugs that interact with bone ECM proteins. The database provides a robust tool for the study of bone weakness and regeneration, and will be a suitable resource for the identification of novel target proteins and therapeutic peptides for the treatment and prevention of osteoporosis. Thus, Phylobone will support emerging therapies targeting novel disease mechanisms to provide a powerful strategy for osteoporosis management in the future ^15^.

## MATERIALS AND METHODS

### Preliminary list of bone ECM proteins

A list of 255 seed proteins previously identified in the literature as proteins present in the ECM was gathered from the UniProt database ^16^. These seed proteins were previously identified in the proteome of the bone ECM of *D. rerio* (n = 243), *H. sapiens* (n = 255), and *Cervus nippon* (n = 57)^17,18^ (supplementary table S1). UniProt codes of *D. rerio* (Zebrafish) proteins that were obsolete (or fractions of protein sequences) were updated with current UniProt codes. Additional putative bone ECM proteins from *H. sapiens* were identified from the literature ^11,18–21^ and gathered from the UniProt database ^16^. Each protein group was initially identified based on the human bone ECM proteome, with a unique code ranging from PB0001 to PB0255.

### Selection of species of interest

We selected 31 representative species of vertebrates to cover a wide range of phylogenetic groups (supplementary Table S2). The list includes a wide coverage of vertebrates, including most common model organisms and other vertebrates that are potentially important as model organisms of bone regeneration and osteoporosis (e.g., six members of the Cervidae family, including *Cervus hanglu yarkandensis, Odocoileus virginianus texanus, Cervus canadensis, Cervus elaphus, Muntiacus muntjak,* and *Muntiacus reevesiveus).* Moreover, we included in the search some species of invertebrates (n = 8) and other taxonomic groups that are phylogenetically distant to vertebrates (e.g., *Arabidopsis,* Archaea, Bacteria, Choanoflagellata, Cnidaria, Ctenophora, Fungi, and Porifera).

### Gathering ECM proteins from public databases

Orthologous proteins were gathered from public repositories of the National Center for Biotechnology Information (NCBI). Phylobone draws its results from NCBI’s Eukaryotic Genome Annotation pipeline ^22^ and NCBI Gene dataset ^23^ to identify orthologous [.1] sequences of some vertebrates (e.g., *M. musculus, Rattus norvegicus, Gallus gallus,* and *Bos taurus).* Additional protein sequences of vertebrates were gathered from protein BLAST searches (blastp) ^24^ with a threshold of 10^-6^ on the NCBI’s non-redundant protein sequences (nr). Synthetic construct sequences (taxid: 32630) and annotated partial proteins were not included in the Phylobone database.

### Phylogenetic analysis and reconstruction of phyletic patterns

Phylogenetic analyses were performed with NGPhylogeny web server ^25^. We utilized the *One Click Workflow* option to obtain multiple sequence alignments (MAFFT algorithm ^26^), clean alignments (BMGE algorithm ^27^) and phylogenetic trees (PhyML algorithm ^28^) of the 255 protein. Phyletic patterns of each protein group were built with custom-made perl scripts and information from NCBI’s Taxonomy ^29^. We selected various species of different taxonomic groups, including 5 primates, 1 rabbit, 2 rodents, 2 carnivores, 10 even-toed ungulates (including 5 members of the family Cervidae), 2 reptiles, 3 birds, 2 frogs, 4 bony fishes, and 8 invertebrates (Supplementary Table 3).

### Functional analysis

Clusters of Orthologous Groups (COG) functional categories and Kyoto Encyclopedia of Genes and Genomes (KEGG) annotations of all the selected proteins in the database are retrieved from eggnog ^30,31^. The dataset of 255 protein groups was mapped onto eggNOG-mapper^30,31^ to identify the COG functional categories involved in bone ECM proteins. COGs are classified into 26 functional groups ^32^. Gene Ontology (GO) enrichment data for human and zebrafish proteins were collected from GO ^33^ with the Panther resource. Benjamini–Hochberg false discovery rate (FDR) correction was used for the searches. Pfam ^34^ was used to retrieve data about the domains of each human protein in the dataset. Pfam has structural and functional information for each domain as well as links to other databases, such as InterPro ^35^. In addition, information on the functional domains of all protein sequences in the dataset was annotated with CD-Search (with default parameters: E-value cut-off of 0.01; composition-corrected scoring: applied; low-complexity regions: not filtered; maximum aligns: 500) ^36^.

### Protein-protein interaction network

Protein–protein interactions of each human protein in the database were retrieved from European Bioinformatics Institute’s Intact [.1] webserver ^37^ and visualized with Cytoscape ^38^. Betweenness centrality was used to evaluate the centrality of each protein in the network. This parameter gives a value to a node calculated according to Equation 1:

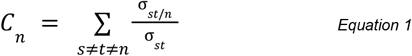

where *C_n_* is the betweenness centrality value of the node n, *σ_st_* is the number of shortest paths from *s* to *t* and σ_*st/n*_ is the number of shortest paths from *s* to *t* that *n* lies in between.

### Targets by existing drugs

To facilitate the research on repurposing drugs for osteoporosis treatment, we used information from the Drug Bank ^39^ and KEGG ^40^ databases to identify bone ECM proteins that are targets of currently available drugs on the market.

## RESULTS AND DISCUSSION

### Phyletic patterns

The phyletic distribution of proteins in the selected group of species shows a high conservation of the number of bone ECM proteins within the vertebrate species (Figure 1 and Supplementary Table 3). Seed proteins to build the Phylobone database were obtained from humans and zebrafish, which were present in 255 (100%) and 251 (98%) protein groups. Most vertebrates were present in ~90% of the protein groups. This finding underscores the significance of the presence of these proteins in the bone ECM, as they exhibit a high degree of conservation across multiple species. Such conservation may have important implications for the function and regulation of bone tissue across vertebrates. Moreover, certain proteins, such as amelogenin (PB0250), are only present in mammals, amphibians, and reptiles because of potential evolutionary adaptations in tetrapods ^41^. Note that there are exceptional gaps in certain members of the Cervidae family (e.g., *C. elaphus, M. reeversi*, and *M. muntjak* have fewer known proteins than the rest of mammals) due to incomplete annotation of the genomes. As expected, only around one-third of the proteins found in invertebrates have homologs in vertebrate species, and in many cases, the levels of similarity were found to be quite low. None of the protein sequences have homologs in lower taxonomic groups, such as bacteria.

**Figure 1.**
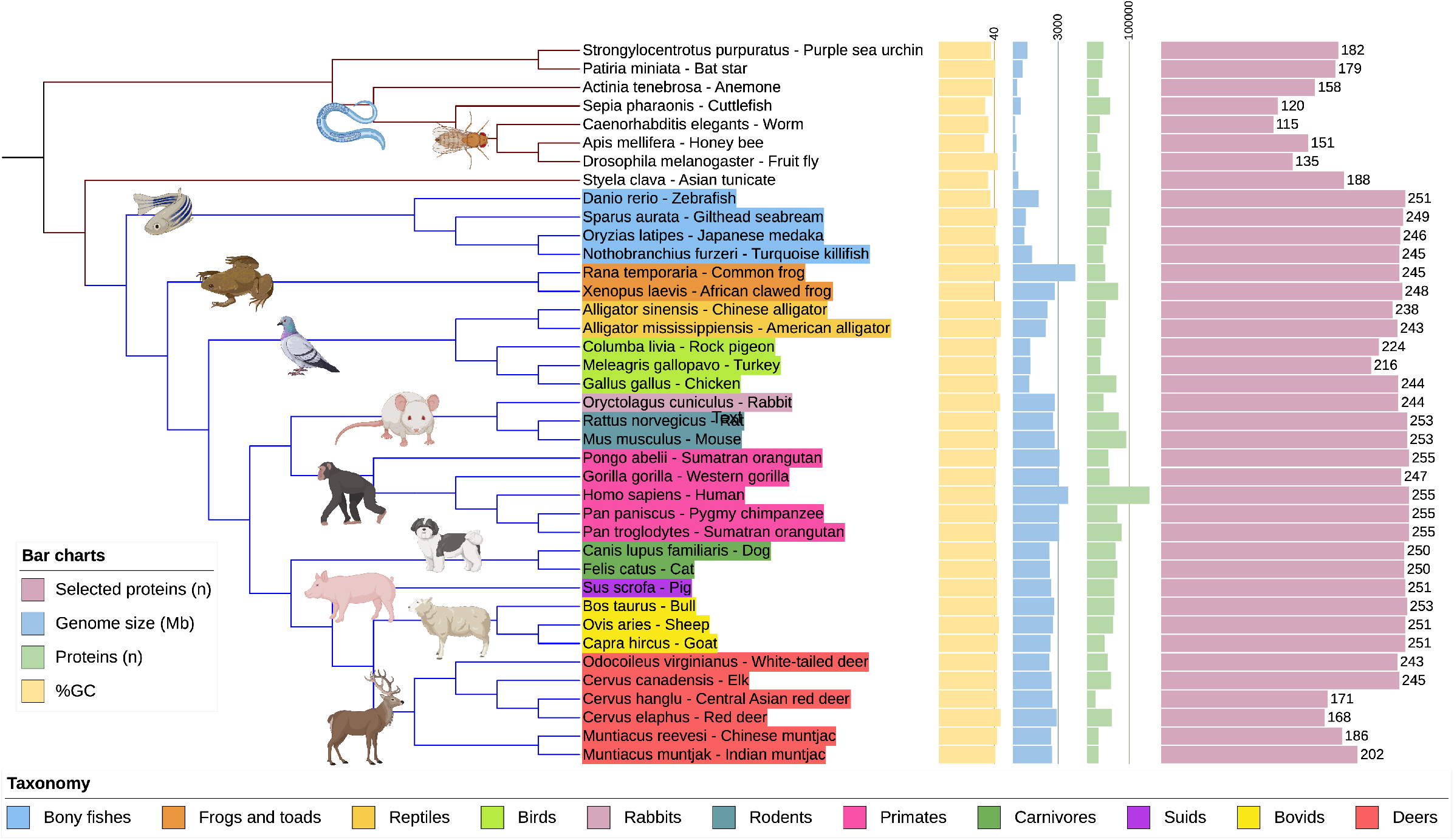
General parameters of the selected species in the Phylobone database. Tree visualization was made with iTOL v5 ^77^ and shows the following datasets: G+C percentage (yellow), genome size (blue), number of proteins (green), number of putative bone extracellular matrix proteins (purple), and taxonomic information.

### Functional analysis

#### Protein functional domains

As expected, the most abundant protein functional domain in the Phylobone dataset is collagen, which is present in 12% of human proteins, covering 90% of the organic fraction of the bone ECM ^2^. Collagen is essential for homeostasis maintenance, and it serves as a scaffold to many other macromolecules and hydroxyapatite, enabling cell attachment and bone resistance to mechanical forces ^1,42,43^. Furthermore, 214 non-collagenous functional domains are present in human sequences of the dataset (Figure 2 and Supplementary Table S4). They are mostly common in bone formation, resorption, cell attachment, or as intermediaries in a variety of metabolic pathways. Leucine-rich repeats (LRRs) are the second most common functional domains due to the large abundance of leucine-rich proteoglycans (SLRPs) in bone ECM ^1^. LRRs play an important role in bone formation and the maintenance of bone homeostasis. Moreover, laminins, von Willebrand factor (VWF), epidermal growth factor (EGF), and trypsin domains are also frequent. Laminins, VWF, and EGF have significant roles in bone regeneration and stability—laminins are involved in cell adhesion, proliferation, and differentiation ^44^; VWF enhances the inhibition of osteoclastogenesis ^45^; and EGF can stimulate bone resorption ^46^. Trypsin is an important domain in the ECM because of its ability to cleave proteins. Several matrix-degrading enzymes, such as cathepsin K and matrix metalloproteinases, also contribute to the efficient degradation of bone ^47^.

**Figure 2.**
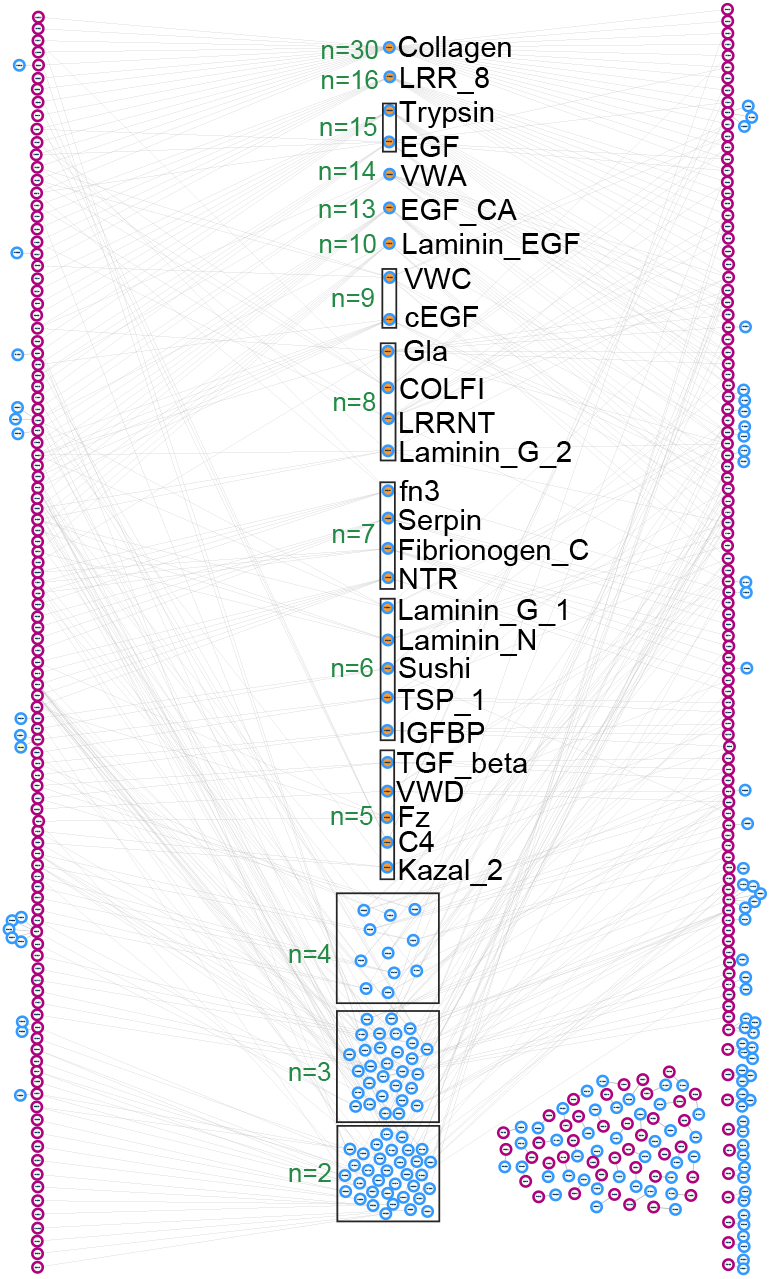
Distribution of protein functional domains in the Phylobone database. Relation between the human proteins of the dataset (purple) and their domains (blue). Dots filled in orange are the most common domains in the database (notice that only domains present in at least 5 proteins are shown). The number of proteins that each domain has is annotated in green (n).

#### Analysis of gene ontology

We compared GO enrichment analyses of biological process, molecular function, and cellular component categories between humans and zebrafish (Figure 3 and supplementary figure 1-6)). GO annotations are more abundant in humans, but there is a certain functional overlap with zebrafish that can be utilized to evaluate the use of zebrafish as a model organism of osteoporosis ^48^. A common category in biological processes is bone mineralization, which is essential for bone regeneration ^49^. In terms of molecular functions, several binding activities are abundant in both humans and zebrafish. As expected, collagen is frequent in the GO cellular component categories of both species, as it is the main component of bone ECM ^1^.

**Figure 3.**
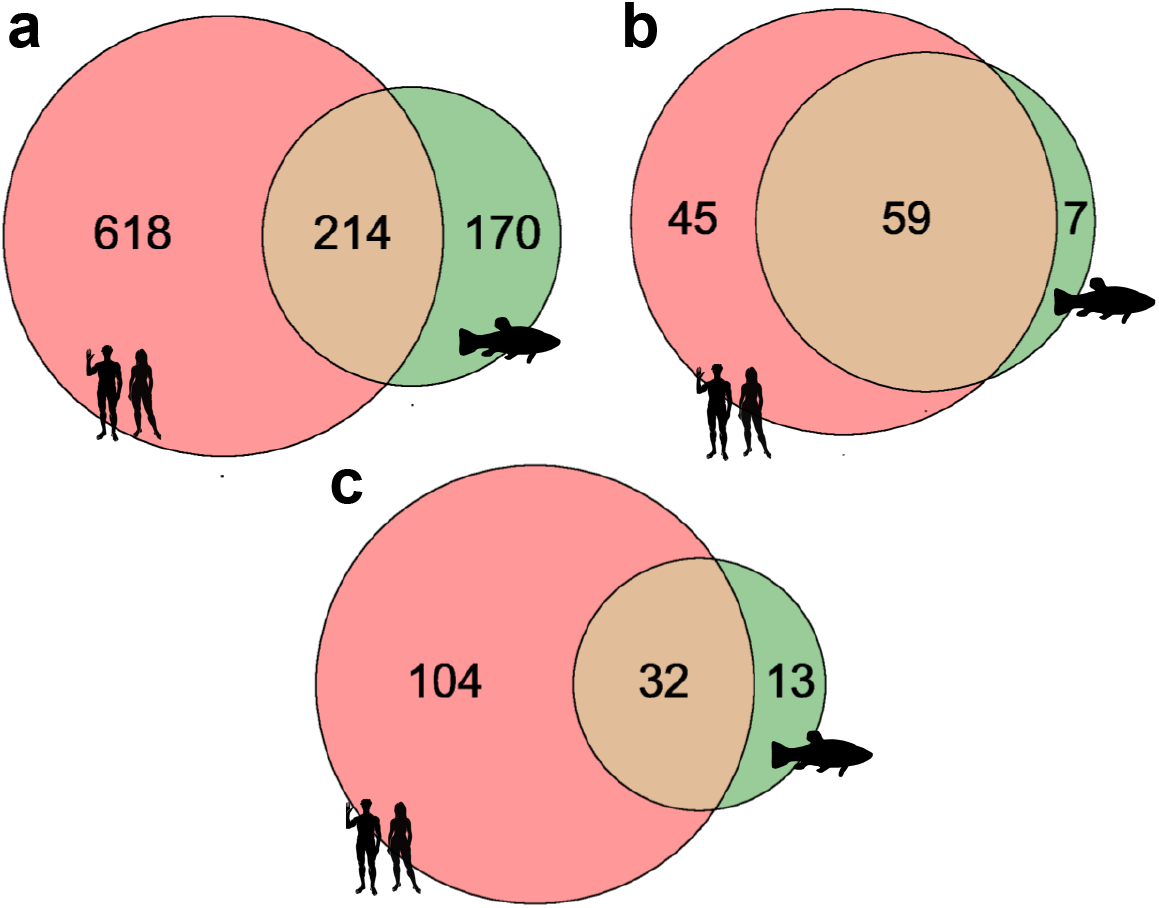
Venn diagram of the number of GO categories in the bone extracellular matrix proteins of *Homo sapiens* and *Danio rerio.* **a.** Biological process categories. **b.** Molecular function categories. **c.** Cellular component categories.

#### Clusters of orthologous groups

The Phylobone database is a compilation of ECM proteins; thus, several proteins are categorized in the COG database under the functions of *extracellular structures* (W) and *signal transduction* (T) (Figure 4a). Signal transduction is an indispensable function of the ECM for controlling homeostasis and transmission of molecular signals into the cell ^50^. Furthermore, proteins involved in bone signaling regulate bone formation and resorption ^49^ (Figure 5). The third and fourth most common COG functional categories are *function unknown* (S) (Phylobone may shed some light on the identification of the functions of these proteins when studied in suitable animal models), as well [.1] as *post-translational modification, protein turnover,* and *chaperones* (O). Proteins of the ECM are expected to be highly integrated to enable signaling functions and to control homeostasis.

**Figure 4.**
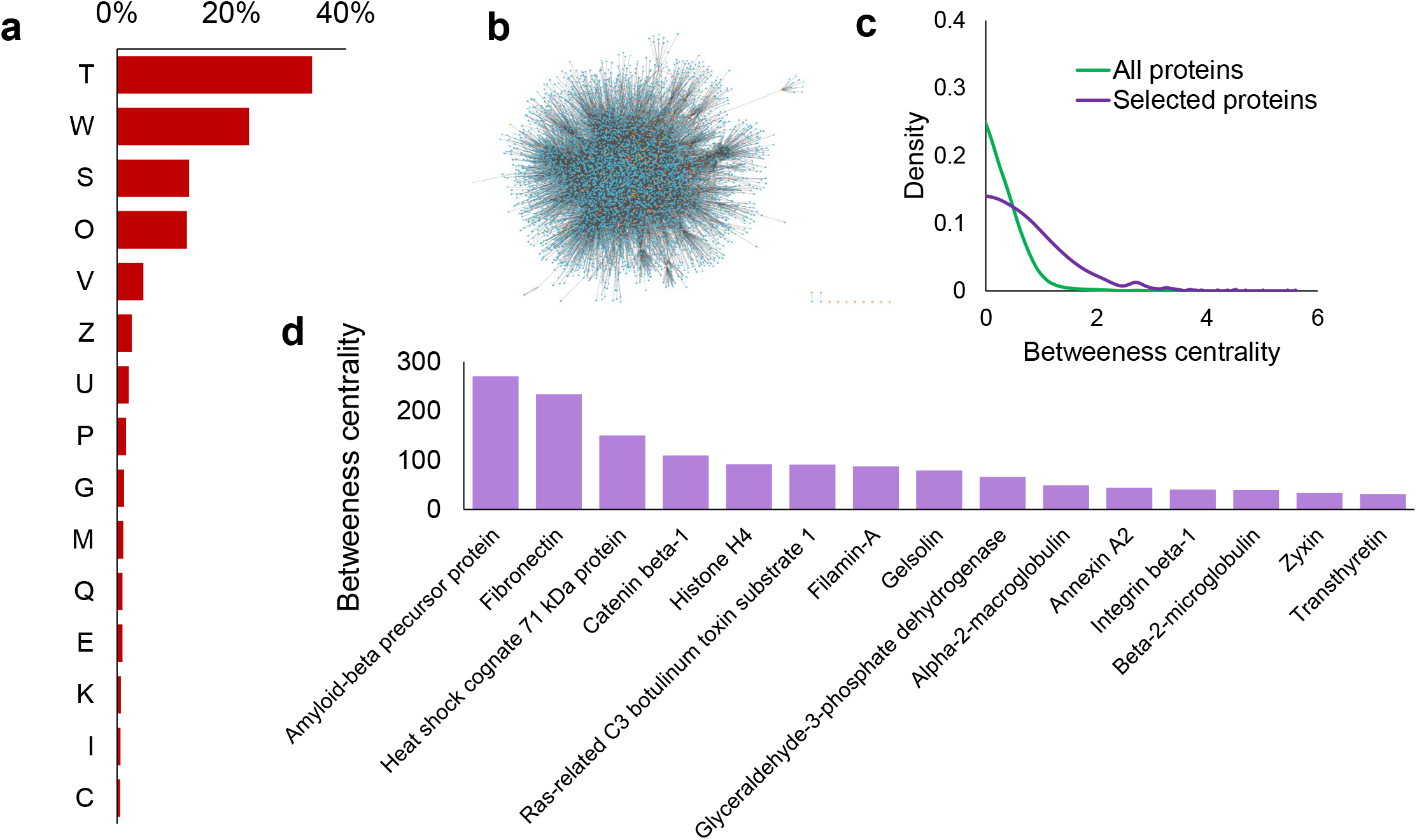
Protein-protein interaction network of human proteins in the bone extracellular matrix. **a.** Percentage of proteins that have each COG functional group. The sequences of all selected species are represented. Signal transduction mechanisms (T), extracellular structures (W), function unknown (S), post-translational modification, protein turnover, chaperones (O), defense mechanisms (V), cytoskeleton (Z), intracellular trafficking, secretion, and vesicular transport (U), inorganic ion transport and metabolism (P), carbohydrate transport and metabolism (G), cell wall/membrane/envelope biogenesis (M), secondary metabolite biosynthesis, transport, and catabolism (Q), amino acid transport and metabolism (E), transcription (K), lipid transport and metabolism (I), and energy proSignal transduction mechanisms (T). **b.** Tree of interactions between proteins. Orange dots are human proteins in the dataset, and blue dots are for other proteins with which they interact. **c.** Gaussian representation of proteins’ betweenness centrality on a logarithmic scale (ln). Green lines represent all 5871 proteins that interact with each other, and orange lines represent the 255 human proteins in the dataset. **d.** Human proteins that have a higher betweenness centrality value in our database.

**Figure 5.**
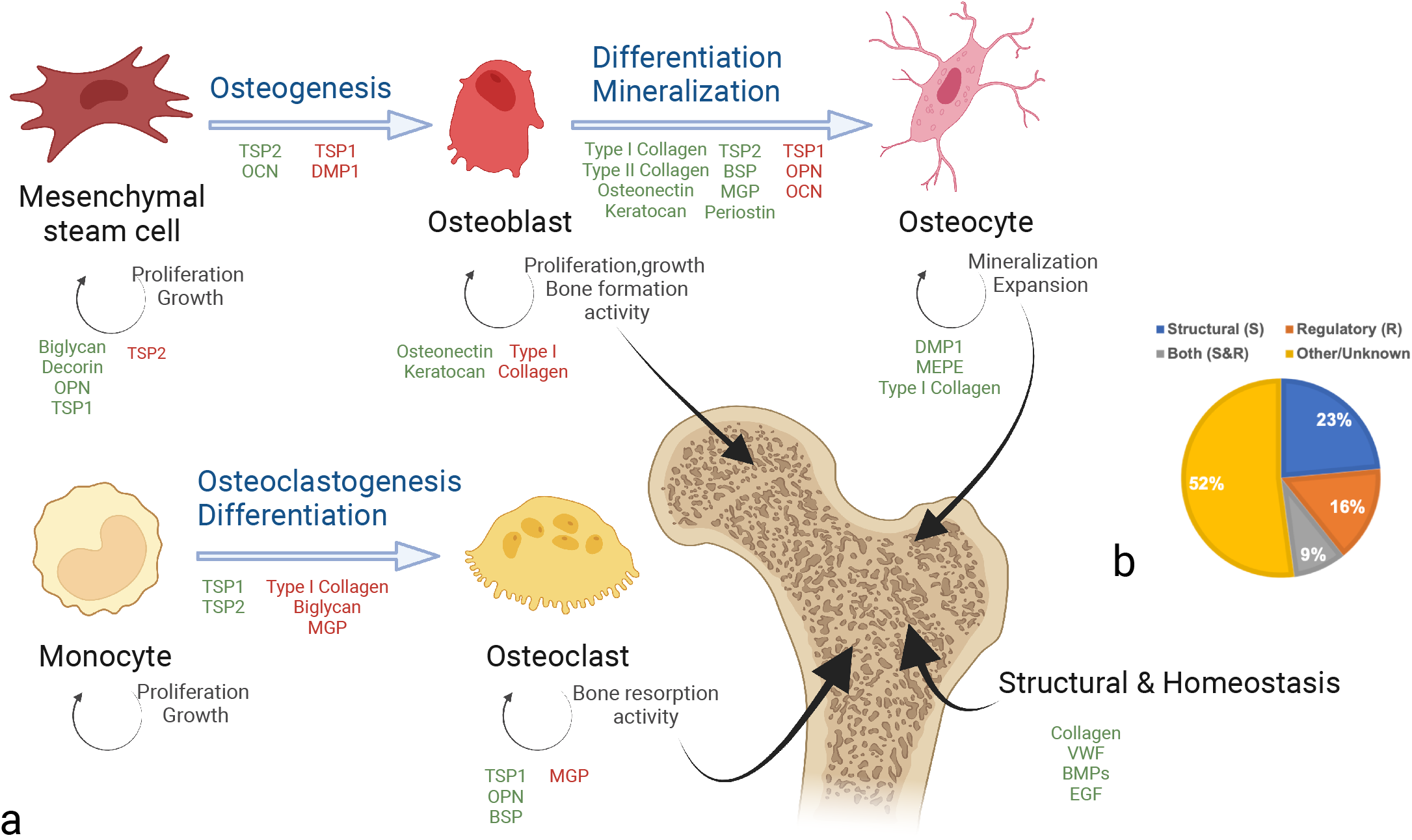
Schematic representation of the state of the art in bone formation and remodeling mediated by extracellular matrix (ECM) proteins. **a.** Simplified representation of processes involving osteocytes, osteoblasts, and osteoclast formation in the bone. ECM proteins can modulate them via activation or inhibition through signaling pathways. These mechanisms allow bone homeostasis to be controlled. **b.** Percentage of proteins with putative structural or regulatory roles in the bone ECM based on annotations in the UniProt database ^16^.

#### Protein interaction network of bone ECM proteins in human

We built a protein interaction network of 5781 human proteins, including putative bone ECM proteins and those that potentially interact (independently of body tissue) with them (Figure 4b). The parameter of betweenness centrality showed higher values in bone ECM proteins compared to those that interacted with them (Figure 4c). This result is in agreement with the central role of bone ECM proteins in maintaining tissue homeostasis and regulatory processe[.1] s. Interactions with cellular components also allow signaling (e.g., cell–matrix interactions permit osteoblast differentiation ^4^). The amyloid beta precursor protein, a transmembrane protein that can be cleaved by a secretase and released in the ECM ^51^, showed the highest value of betweenness centrality (Figure 4d). This protein regulates osteoclast function (affecting bone remodeling), is present in both bone and brain, and can link osteoporosis and neurodegenerative diseases ^52,53^. Fibronectin also had one of the largest values of betweenness centrality. This glycoprotein of the ECM extracellular matrix has an important role in osteogenesis because it promotes pre-osteoblast mineralization and differentiation ^54^. Furthermore, fibronectin acts as a scaffold for the cleavage of procollagen (interacting with both bone morphogenic protein 1 and procollagen) ^55^. This ability to bind to other ECM proteins gives fibronectin its high centrality value and an important role in bone formation and regeneration with a single mechanism.

### How to use the Phylobone database to study bone regeneration and osteoporosis

Osteoporosis is one of the most common bone problems in the middle age and elderly population worldwide ^56,57^. Approximately 9 million fractures per year (one every three seconds) are caused by osteoporosis, which contributes significantly to morbidity and mortality in developed countries ^15^. Given that life expectancy is increasing globally, osteoporosis has become an emerging topic, as it significantly affects the quality of life of individuals in most countries ^15^. Osteoporosis is a metabolic disease caused by an imbalance between bone anabolism and catabolism ^48^. In general, the recommendations to reduce the risk of osteoporosis are intake of adequate calcium, exposure to sunlight, intake of vitamin D, and engaging in weight-bearing exercises. It is known that a reduction in mechanical loading due to prolonged bed rest or long-term exposure to microgravity can lead to a reduction in bone mass ^58,59^. The mechanisms of weight-bearing exercises to prevent osteoporosis are not fully understood; yet, during weight-bearing exercise, collagen fibrils, which represent 90% of the organic components of the bone ECM, generate electricity that is stored in the inorganic components of the ECM ^60^. Since bone ECM has both structural and regulatory roles, non-collagenous organic components have a key role in bone regulation by mechanical loading ^3,4^.

Several non-collagenous proteins of the bone ECM proteome have been identified previously, including osteocalcin, osteonectin, and R-spondins; however, the list and roles of bone ECM proteins in humans (and model organisms) are not fully elucidated ^1^. Some ECM proteins have been reported to play an important role in promoting bone formation. Type I collagen, the most abundant protein in the bone ECM, plays a structural role in providing mechanical support, such as bone strength and fragility ^61^, and regulates integrin receptors for osteoblasts ^62^. Moreover, the bone ECM proteome comprises hundreds of non-collagenous proteins with structural, regulatory, and homeostatic roles ^63^ (Figure 5). Osteopontin is a non-collagenous protein in bone ECM related to mechanical stress with multifunctional functions ^64^. This protein is produced by osteoblasts as well as osteoclasts ^65^, and it inhibits the activities of osteoblasts while promoting osteoclast activities ^66^. There are other proteins responsible for maintaining bone homeostasis, such as bone morphogenetic proteins (BMPs)^67^.

### Phylobone database to study mechanobiology

Outcomes from the Phylobone project will also have an impact on the field of mechanobiology. Several publications have determined the role of some cell types, cellular membrane receptors, and bone ECM proteins in mechanical loading ^64,68,69^. For instance, osteocytes are known as mechanosensing cells in bone tissue, as they transduce mechanical signals to biochemical responses ^68^. PIEZO1 is a mechanosensitive ion channel component that became a topic of discussion because its discoverer won the Nobel Prize in 2021 ^70^. PIEZO1 regulates homeostasis *via* crosstalk of osteoblasts (bone-making cells) and osteoclasts (bone-eating cells)^69^. Further, mechanical properties, especially fluid shear stress in bone ECM, have significant effects on cells and their interactions ^71^. However, only a few non-collagenous proteins of the ECM (e.g., osteopontin) have been related to mechanical stress ^64^.

### Potential repurposing of existing drugs

The Phylobone database is a key resource for advancing research on the prevention of osteoporosis and the development of new treatments for bone regeneration. The most common treatments for osteoporosis are bisphosphonates, monoclonal receptor activator of nuclear factor-kB ligand (RANKL) antibodies, monoclonal sclerostin antibodies, and a parathyroid hormone peptide ^72^. The parathyroid hormone peptide increases osteoblast activity and inhibits osteoclast recruitment, whereas the targets of the other treatments inhibit osteoclast resorption ^73^. However, these treatments have limitations and problems with side effects and serious risks. For instance, bisphosphonates have been reported to have higher fracture rates after long-term use, characterized as more than six years and requiring a “bisphosphonate holiday”^74^. RANKL antibodies have been reported to have some side effects, such as skin eczema, flatulence, cellulitis, and osteonecrosis of the jaw ^75^. Thus, there is still a need for osteoporosis treatment. However, finding new drugs is expensive; thus, repurposing existing drugs is a main goal for the identification of novel treatments and preventive methods. Given that several proteins of the bone ECM proteome have been identified as playing a regulatory role in osteogenesis and bone degradation ^1^, we hypothesize that key proteins of the ECM involved in mechanical loading could be potential drug targets for the treatment and prevention of osteoporosis. The Phylobone database includes 36 proteins out of 255 proteins from the bone ECM that have existing drugs and are putative candidates for repurposing existing drugs (Table 1 and Supplementary Table S5), including 10 drugs that interact with more than one protein (Table 2).

**Table 1.** List of the 36 bone extracellular matrix (ECM) proteins that target at least one available drug on the market.

| PB Code | Uniprot Code | Uniprot Name | N. of Drugs | Protein Description |
| --- | --- | --- | --- | --- |
| PB0124 | P00742 | FA10_HUMAN | 17 | Coagulation factor X |
| PB0003 | P00734 | THRB_HUMAN | 13 | Prothrombin |
| PB0080 | P01031 | CO5_HUMAN | 12 | Complement C5 |
| PB0242 | P02766 | TTHY_HUMAN | 10 | Transthyretin |
| PB0107 | P14780 | MMP9_HUMAN | 8 | Matrix metalloproteinase-9 |
| PB0178 | O14793 | GDF8_HUMAN | 7 | Growth/differentiation factor 8 |
| PB0204 | P08253 | MMP2_HUMAN | 6 | 72 kDa type IV collagenase |
| PB0235 | P06756 | ITAV_HUMAN | 5 | Integrin alpha-V |
| PB0238 | P00747 | PLMN_HUMAN | 4 | Plasminogen |
| PB0067 | P00740 | FA9_HUMAN | 3 | Coagulation factor IX |
| PB0008 | P02743 | SAMP_HUMAN | 3 | Serum amyloid P-component |
| PB0132 | P05121 | PAI1_HUMAN | 3 | Plasminogen activator inhibitor 1 |
| PB0236 | P05556 | ITB1_HUMAN | 3 | Integrin beta-1 |
| PB0112 | P45452 | MMP13_HUMAN | 3 | Collagenase 3 |
| PB0160 | P61812 | TGFB2_HUMAN | 3 | Transforming growth factor beta-2 proprotein |
| PB0159 | P00751 | CFAB_HUMAN | 2 | Complement factor B |
| PB0161 | P21333 | FLNA_HUMAN | 2 | Filamin-A |
| PB0100 | P43235 | CATK_HUMAN | 2 | Cathepsin K |
| PB0017 | P69905 | HBA_HUMAN | 2 | Hemoglobin subunit alpha |
| PB0175 | P01009 | A1AT_HUMAN | 1 | Alpha-1-antitrypsin |
| PB0030 | P01024 | CO3_HUMAN | 1 | Complement C3 |
| PB0187 | P01042 | KNG1_HUMAN | 1 | Kininogen-1 |
| PB0212 | P02753 | RET4_HUMAN | 1 | Retinol-binding protein 4 |
| PB0077 | P04114 | APOB_HUMAN | 1 | Apolipoprotein B-100 |
| PB0092 | P05186 | PPBT_HUMAN | 1 | Alkaline phosphatase, tissue-nonspecific isozyme |
| PB0174 | P07477 | TRY1_HUMAN | 1 | Serine protease 1 |
| PB0234 | P08648 | ITA5_HUMAN | 1 | Integrin alpha-5 |
| PB0201 | P10600 | TGFB3_HUMAN | 1 | Transforming growth factor beta-3 proprotein |
| PB0200 | P10909 | CLUS_HUMAN | 1 | Clusterin |
| PB0196 | P29279 | CCN2_HUMAN | 1 | CCN family member 2 |
| PB0246 | P63000 | RAC1_HUMAN | 1 | Ras-related C3 botulinum toxin substrate 1 |
| PB0036 | Q92743 | HTRA1_HUMAN | 1 | Serine protease HTRA1 |
| PB0144 | Q9BX93 | PG12B_HUMAN | 1 | Group XIIb secretory phospholipase A2-like protein |
| PB0254 | Q9BXY4 | RSPO3_HUMAN | 1 | R-spondin-3 |
| PB0181 | Q9GZV9 | FGF23_HUMAN | 1 | Fibroblast growth factor 23 |
| PB0203 | Q9UNI1 | CELA1_HUMAN | 1 | Chymotrypsin-like elastase family member 1 |

**Table 2.** Drugs that interact with more than one protein of the bone extracellular matrix (ECM). There are 114 drugs that interact with at least 1 bone ECM protein (104 drugs interact with 1 protein, 9 drugs interact with 2 proteins, and 1 drug interacts with 3 proteins). The complete matrix of drug and protein interactions is available in Supplementary Table 5.

| Drug | Related proteins | Drug efficacy |
| --- | --- | --- |
| Batimastat | P08253, P14780 | Antineoplastic, Matrix metalloproteinase inhibitor |
| Cipemastat, Trocade | P14780, P45452 | Antirheumatic, Cartilage protectant, Matrix metalloproteinase inhibitor |
| Ilomastat | P08253, P14780 | Antineoplastic, Matrix metalloproteinase inhibitor |
| Marimastat | P08253, P14780 | Antineoplastic, Matrix metalloproteinase inhibitor |
| Rebimastat | P08253, P14780 | Antineoplastic, Matrix metalloproteinase inhibitor |
| Tanomastat | P08253, P14780 | Antineoplastic, Matrix metalloproteinase inhibitor |
| Volociximab | P05556, P08648 | Aging-related macular degeneration therapeutic agent, Antineoplastic, Angiogenesis inhibitor, Anti-alpha5beta1 integrin antibody |
| Fresolimumab | P10600, P61812 | Immunomodulator, Anti-transforming growth factor beta antibody |
| Emicizumab, Hemlibra | P00740, P00742 | Bleeding suppressant, Factor VIII replacement |
| Prinomastat | P08253, P14780, P45452 | Antineoplastic, Angiogenesis inhibitor, Matrix metalloproteinase inhibitor |

\* **P08253**: 72 kDa type IV collagenase; **P14780**: Matrix metalloproteinase-9; **P45452**: Collagenase 3; **P05556**: Integrin beta-1; **P08648** Integrin alpha-5; **P10600**: Transforming growth factor beta-3 proprotein; **P61812**: Transforming growth factor beta-2 proprotein; **P00740**: Coagulation factor IX; **P00742**: Coagulation factor X

## AVAILABILITY

The Phylobone database is freely accessible at https://phylobone.com. The current version of the dataset includes 8,615 putative bone ECM proteins from 39 species of vertebrates and invertebrates, and is categorized into 255 protein groups. Each protein in the database is annotated with basic information that includes its name, organism, a general protein description, a list of gene ontologies (GO) associated, protein–protein interactions (PPI), functional domains, metabolic pathways, and drugs. We have precomputed a phyletic profile of proteins and species based on the identification of orthologous sequences in vertebrates, including common model organisms. Protein sequences, multiple sequence alignments, and PhyML phylogenetic trees are available to visualize on the web browser and download. Seed sequences of humans and zebrafish, which have been used to build the database, are also available. Moreover, the database includes several links to external resources, including Uniprot ^16^, Protein Atlas ^76^, Intact ^37^, InterPro ^35^ and the BLAST alignment tool ^24^. All proteins have been mapped onto currently existing drugs and the database includes links to the DrugBank ^39^ and KEGG ^40^ databases.

## CONCLUSIONS AND FUTURE DEVELOPMENTS

The aim of the Phylobone research project is to provide a platform to study bone regeneration and osteoporosis in light of (biological) evolution. The current database includes the most comprehensive repository of bone ECM proteins in human and animal models. The phylobone database provides a bioinformatics resource that includes sequences, phylogenetics, and functionals. Thus, this resource will be helpful in addressing a major technical challenge in bone biology: the identification of non-collagenous proteins involved in bone formation, regeneration, and progression of osteoporosis disease. In the future, we anticipate an update of the Phylobone database with more experimental data on the regulatory role of bone ECM proteins in bone regeneration and osteoporosis.

## ACKNOWLEDGEMENT

We thank members of the Phylobone team and collaborators for their helpful discussions.

