## Supplementary material for "Phylobone: A comprehensive database of bone extracellular matrix proteins in human and model organisms" (from another version of this preprint: version 1, 2023-03-29)

**SUPPLEMENTARY FIGURES**

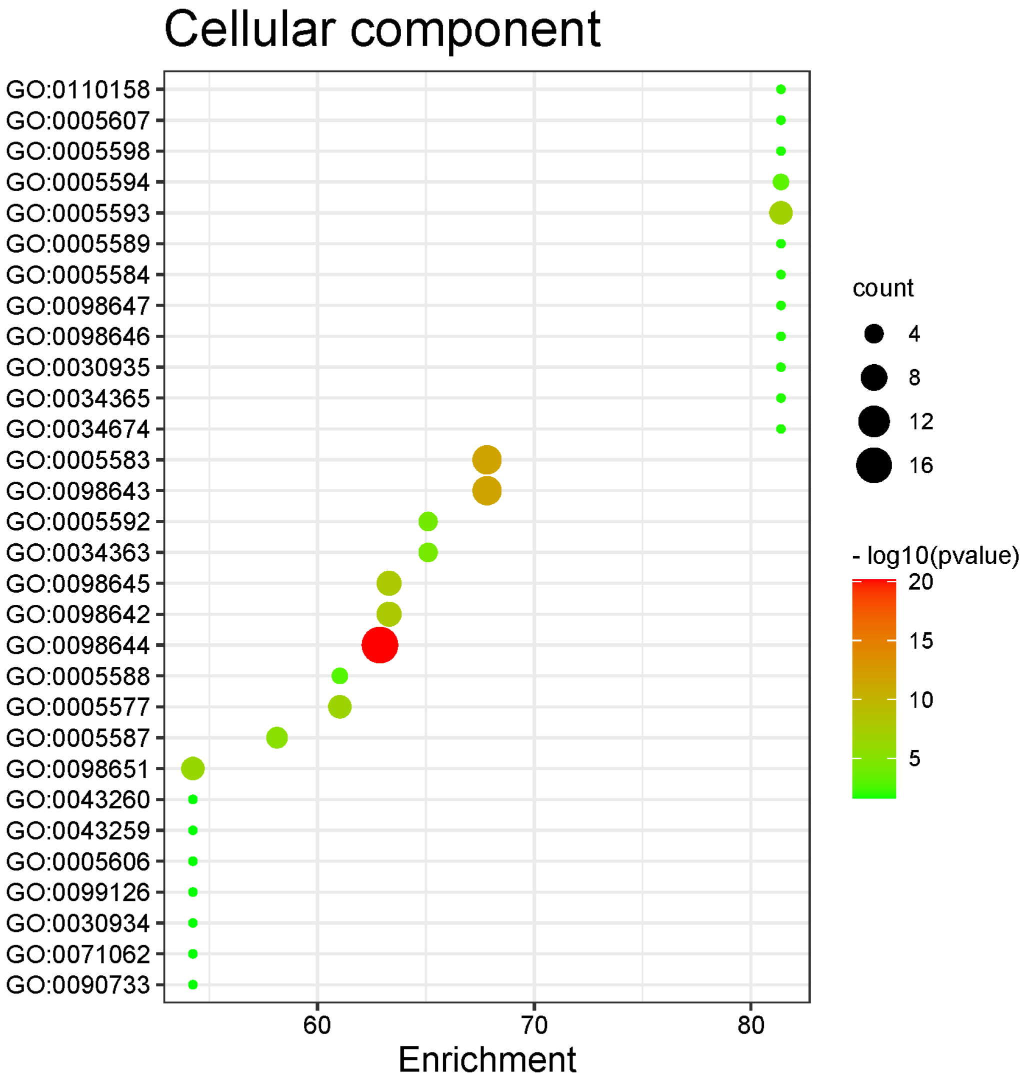


**Supplementary figure S1. Representation of cellular components GO categories with higher enrichment in bone ECM proteins of *Homo sapiens*.**
GO:0110158, calpain complex; GO:0005607, laminin-2 complex; GO:0005598, short-chain collagen trimer; GO:0005594, collagen type IX trimer; GO:0005593, FACIT collagen trimer; GO:0005589, collagen type VI trimer; GO:0005584, collagen type I trimer; GO:0098647, collagen beaded filament; GO:0098646, collagen sheet; GO:0030935, sheet-forming collagen trimer; GO:0034365, discoidal high-density lipoprotein particle; GO:0034674, integrin alpha5-beta1 complex; GO:0005583, fibrillar collagen trimer; GO:0098643, banded collagen fibril; GO:0005592, collagen type XI trimer; GO:0034363, intermediate-density lipoprotein particle; GO:0098645, collagen network; GO:0098642, network-forming collagen trimer; GO:0098644, complex of collagen trimers; GO:0005588, collagen type V trimer; GO:0005577, fibrinogen complex; GO:0005587, collagen type IV trimer; GO:0098651, basement membrane collagen trimer; GO:0043260, laminin-11 complex; GO:0043259, laminin-10 complex; GO:0005606, laminin-1 complex; GO:0099126, transforming growth factor beta complex; GO:0030934, anchoring collagen complex; GO:0071062, alphav-beta3 integrin-vitronectin complex; and GO:0090733, tenascin complex.

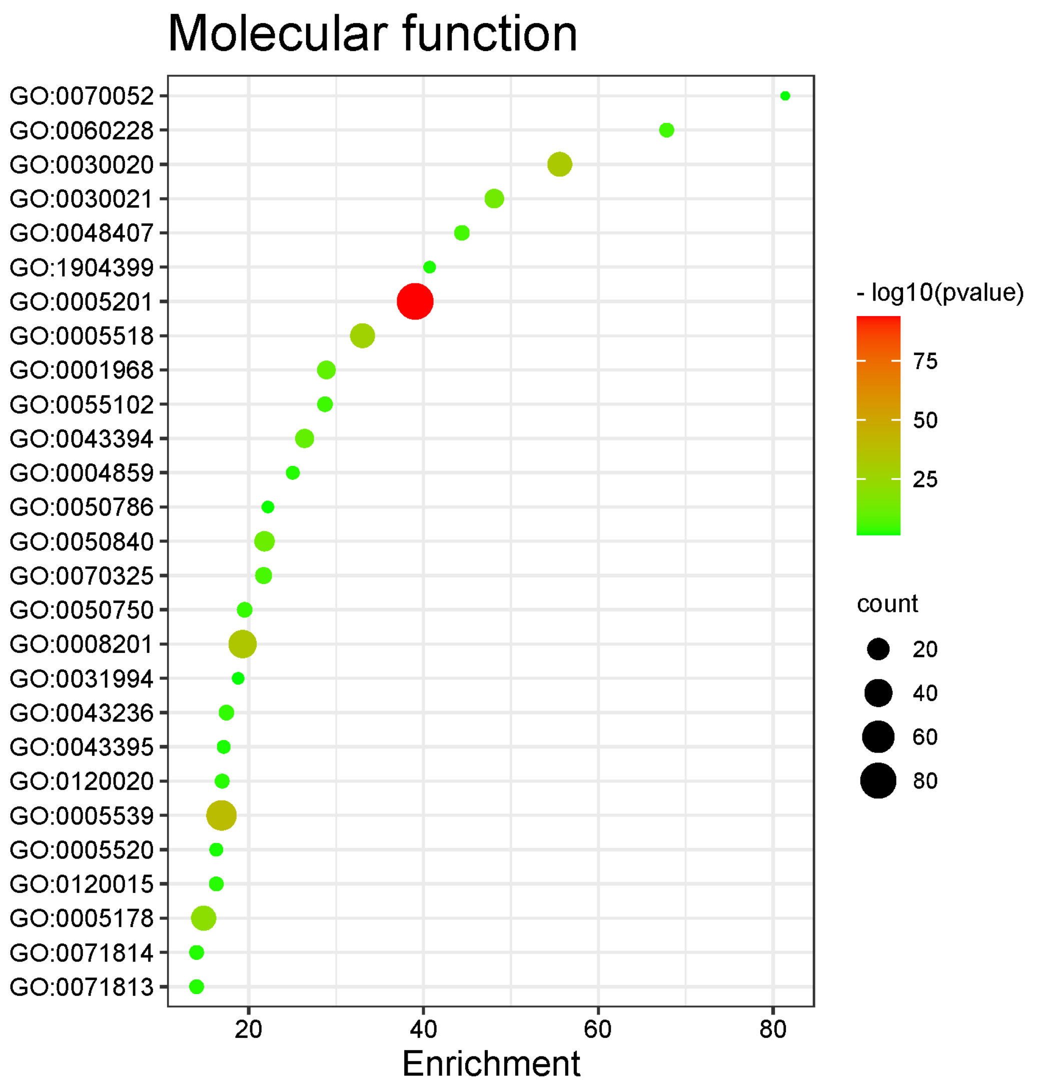


**Supplementary figure S2. Representation of molecular function GO categories with higher enrichment in bone ECM proteins of *Homo sapiens*.**
GO:0070052, collagen V binding; GO:0060228, phosphatidylcholine-sterol O-acyltransferase activator activity; GO:0030020, extracellular matrix structural constituent conferring tensile strength; GO:0030021, extracellular matrix structural constituent conferring compression resistance; GO:0048407, platelet-derived growth factor binding; GO:1904399, heparan sulfate binding; GO:0005201, extracellular matrix structural constituent; GO:0005518, collagen binding; GO:0001968, fibronectin binding; GO:0055102, lipase inhibitor activity; GO:0043394, proteoglycan binding; GO:0004859, phospholipase inhibitor activity; GO:0050786, RAGE receptor binding; GO:0050840, extracellular matrix binding; GO:0070325, lipoprotein particle receptor binding; GO:0050750, low-density lipoprotein particle receptor binding; GO:0008201, heparin binding; GO:0031994, insulin-like growth factor I binding; GO:0043236, laminin binding; GO:0043395, heparan sulfate proteoglycan binding; GO:0120020, cholesterol transfer activity; GO:0005539, glycosaminoglycan binding; GO:0005520, insulin-like growth factor binding; GO:0120015, sterol transfer activity; GO:0005178, integrin binding; GO:0071814, protein-lipid complex binding; and GO:0071813, lipoprotein particle binding.

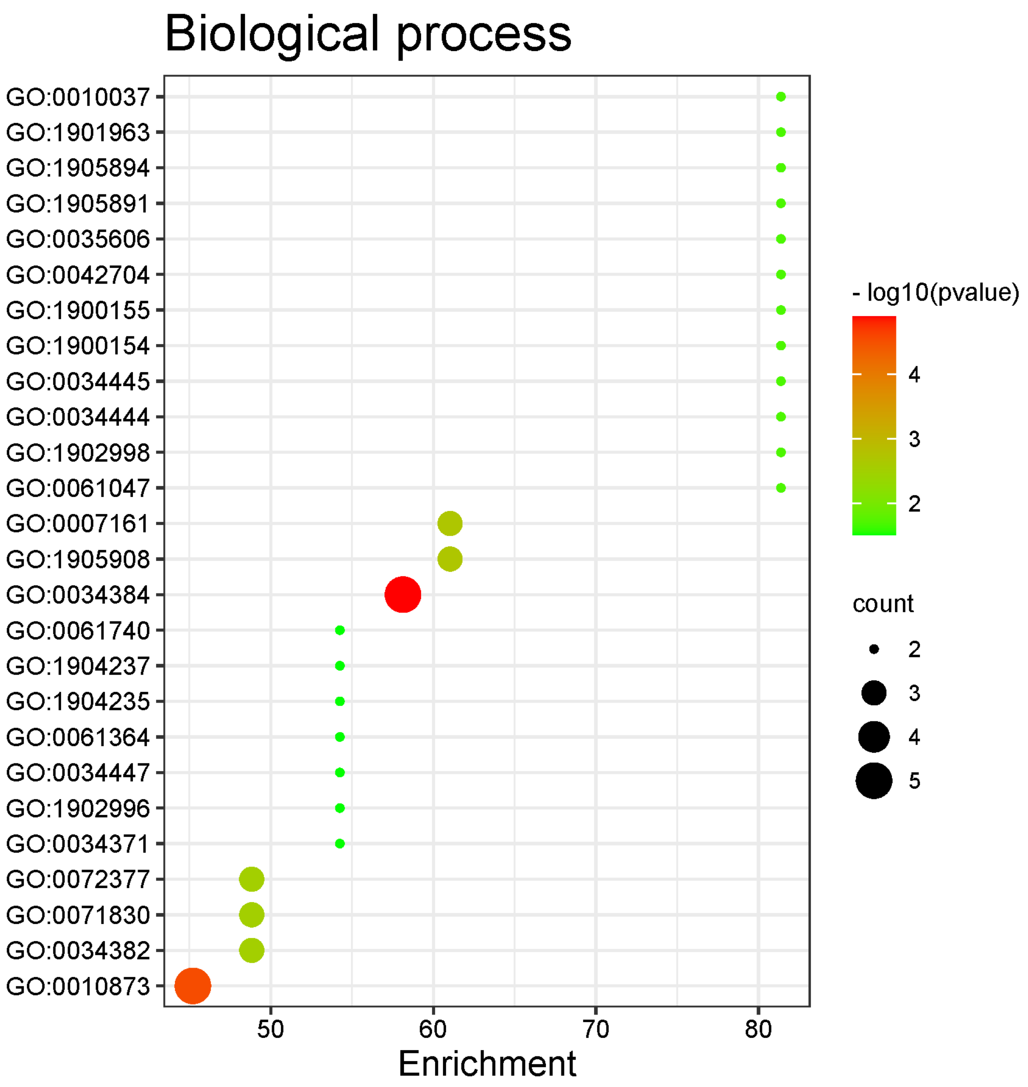


**Supplementary figure S3. Representation of biological process GO categories with higher enrichment in bone ECM proteins of *Homo sapiens*.**

GO:0010037, response to carbon dioxide; GO:1901963, regulation of cell proliferation involved in outflow tract morphogenesis; GO:1905894, regulation of cellular response to tunicamycin; GO:1905891, regulation of cellular response to thapsigargin; GO:0035606, peptidyl-cysteine S-trans-nitrosylation; GO:0042704, uterine wall breakdown; GO:1900155, negative regulation of bone trabecula formation; GO:1900154, regulation of bone trabecula formation; GO:0034445, negative regulation of plasma lipoprotein oxidation; GO:0034444, regulation of plasma lipoprotein oxidation; GO:1902998, positive regulation of neurofibrillary tangle assembly; GO:0061047, positive regulation of branching involved in lung morphogenesis; GO:0007161, calcium-independent cell-matrix adhesion; GO:1905908, positive regulation of amyloid fibril formation; GO:0034384, high-density lipoprotein particle clearance; GO:0061740, protein targeting to lysosome involved in chaperone-mediated autophagy; GO:1904237, positive regulation of substrate-dependent cell migration, cell attachment to substrate; GO:1904235, regulation of substrate-dependent cell migration, cell attachment to substrate; GO:0061364, apoptotic process involved in luteolysis; GO:0034447, very-low-density lipoprotein particle clearance; GO:1902996, regulation of neurofibrillary tangle assembly; GO:0034371, chylomicron remodeling; GO:0072377, blood coagulation, common pathway; GO:0071830, triglyceride-rich lipoprotein particle clearance; GO:0034382, chylomicron remnant clearance; and GO:0010873, positive regulation of cholesterol esterification.

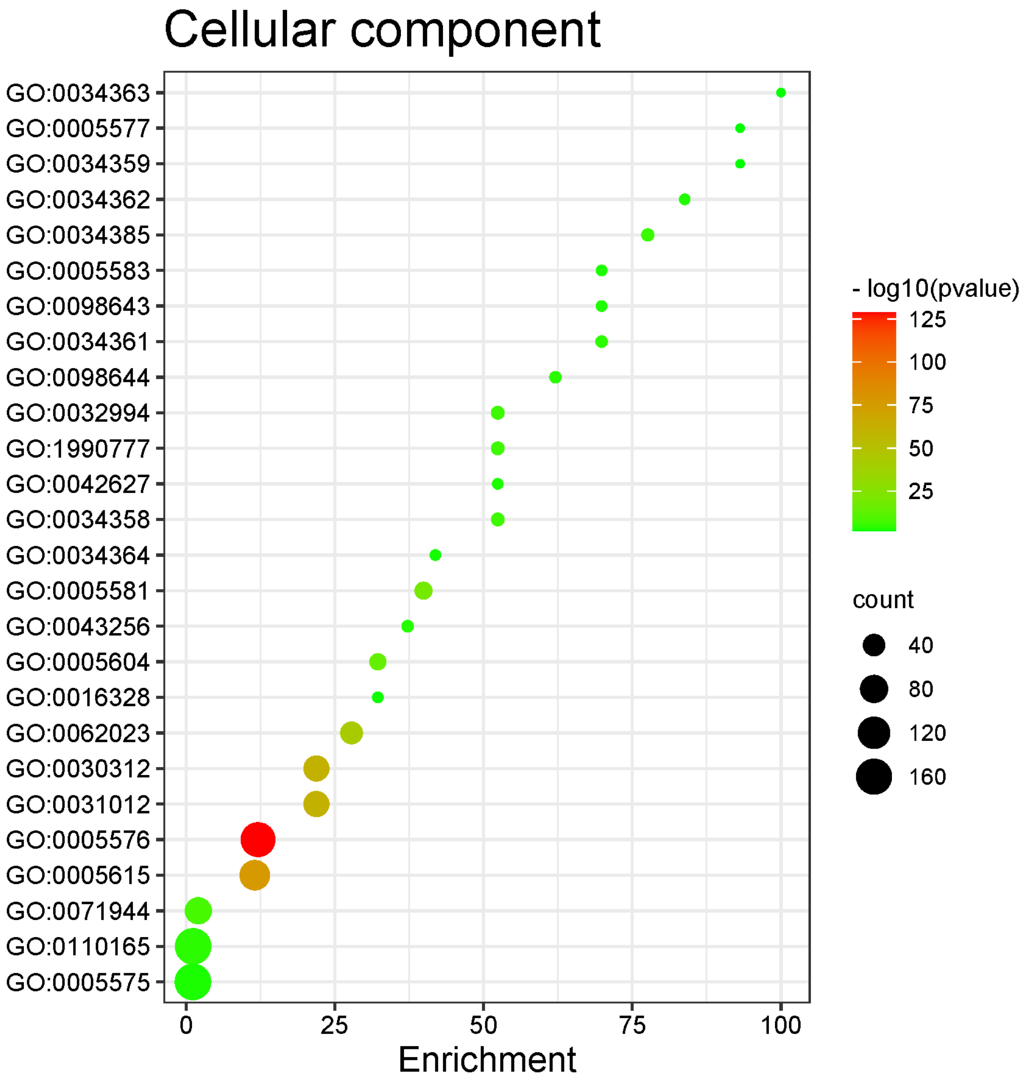


**Supplementary figure S4. Representation of cellular components GO categories with higher enrichment in bone ECM proteins of *Danio rerio*.**
GO:0034363, intermediate-density lipoprotein particle; GO:0005577, fibrinogen complex; GO:0034359, mature chylomicron; GO:0034362, low-density lipoprotein particle; GO:0034385, triglyceride-rich plasma lipoprotein particle; GO:0005583, fibrillar collagen trimer; GO:0098643, banded collagen fibril; GO:0034361, very-low-density lipoprotein particle; GO:0098644, complex of collagen trimers; GO:0032994, protein-lipid complex; GO:1990777, lipoprotein particle; GO:0042627, chylomicron; GO:0034358, plasma lipoprotein particle; GO:0034364, high-density lipoprotein particle; GO:0005581, collagen trimer; GO:0043256, laminin complex; GO:0005604, basement membrane; GO:0016328, lateral plasma membrane; GO:0062023, collagen-containing extracellular matrix; GO:0030312, external encapsulating structure; GO:0031012, extracellular matrix; GO:0005576, extracellular region; GO:0005615, extracellular space; GO:0071944, cell periphery; GO:0110165, cellular anatomical entity; and GO:0005575, cellular_component. Points with Enrichment 100 mean 100 or more.

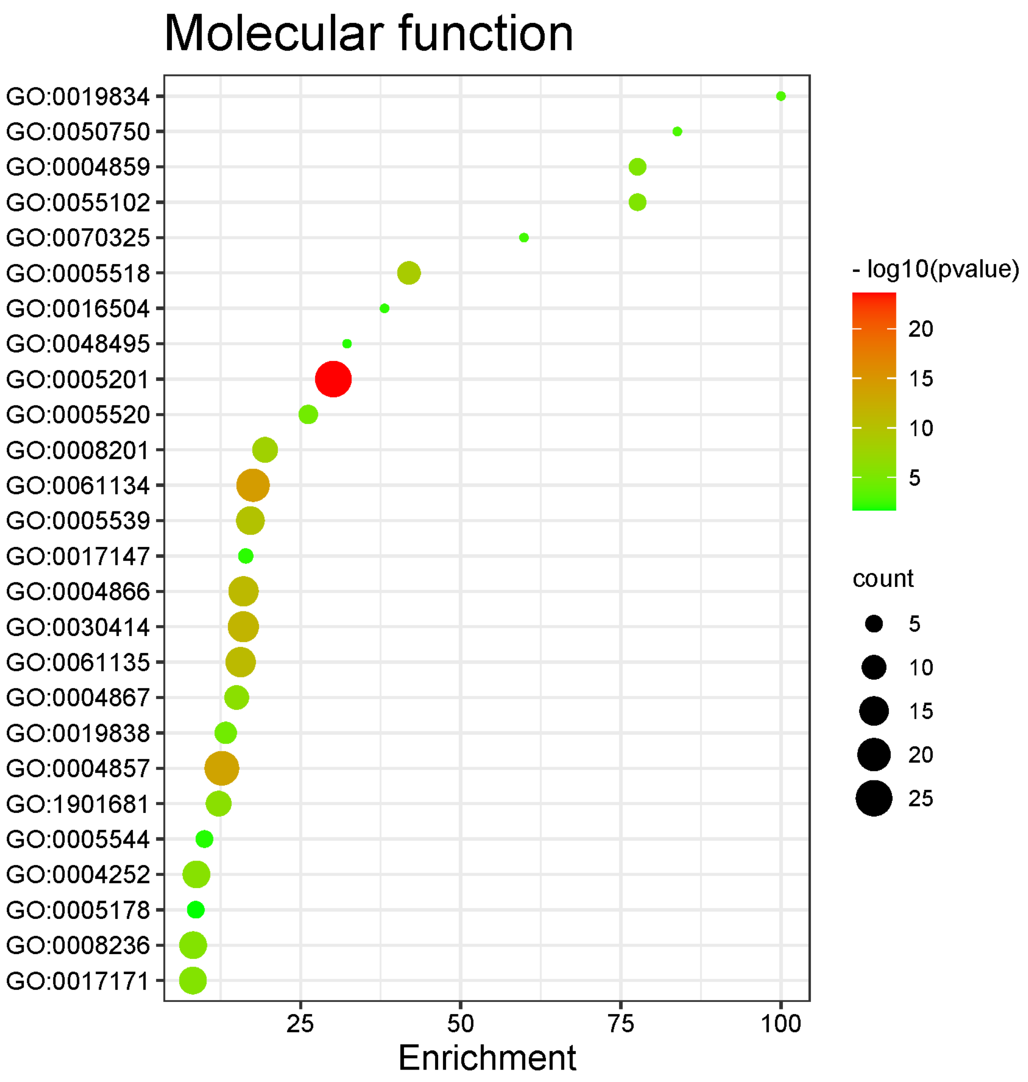


**Supplementary figure S5. Representation of molecular function GO categories with higher enrichment in bone ECM proteins of *Danio rerio*.**

GO:0019834, phospholipase A2 inhibitor activity; GO:0050750, low-density lipoprotein particle receptor binding; GO:0004859, phospholipase inhibitor activity; GO:0055102, lipase inhibitor activity; GO:0070325, lipoprotein particle receptor binding; GO:0005518, collagen binding; GO:0016504, peptidase activator activity; GO:0048495, Roundabout binding; GO:0005201, extracellular matrix structural constituent; GO:0005520, insulin-like growth factor binding; GO:0008201, heparin binding; GO:0061134, peptidase regulator activity; GO:0005539, glycosaminoglycan binding; GO:0017147, Wnt-protein binding; GO:0004866, endopeptidase inhibitor activity; GO:0030414, peptidase inhibitor activity; GO:0061135, endopeptidase regulator activity; GO:0004867, serine-type endopeptidase inhibitor activity; GO:0019838, growth factor binding; GO:0004857, enzyme inhibitor activity; GO:1901681, sulfur compound binding; GO:0005544, calcium-dependent phospholipid binding; GO:0004252, serine-type endopeptidase activity; GO:0005178, integrin binding; GO:0008236, serine-type peptidase activity; and GO:0017171, serine hydrolase activity. Points with Enrichment 100 mean 100 or more.

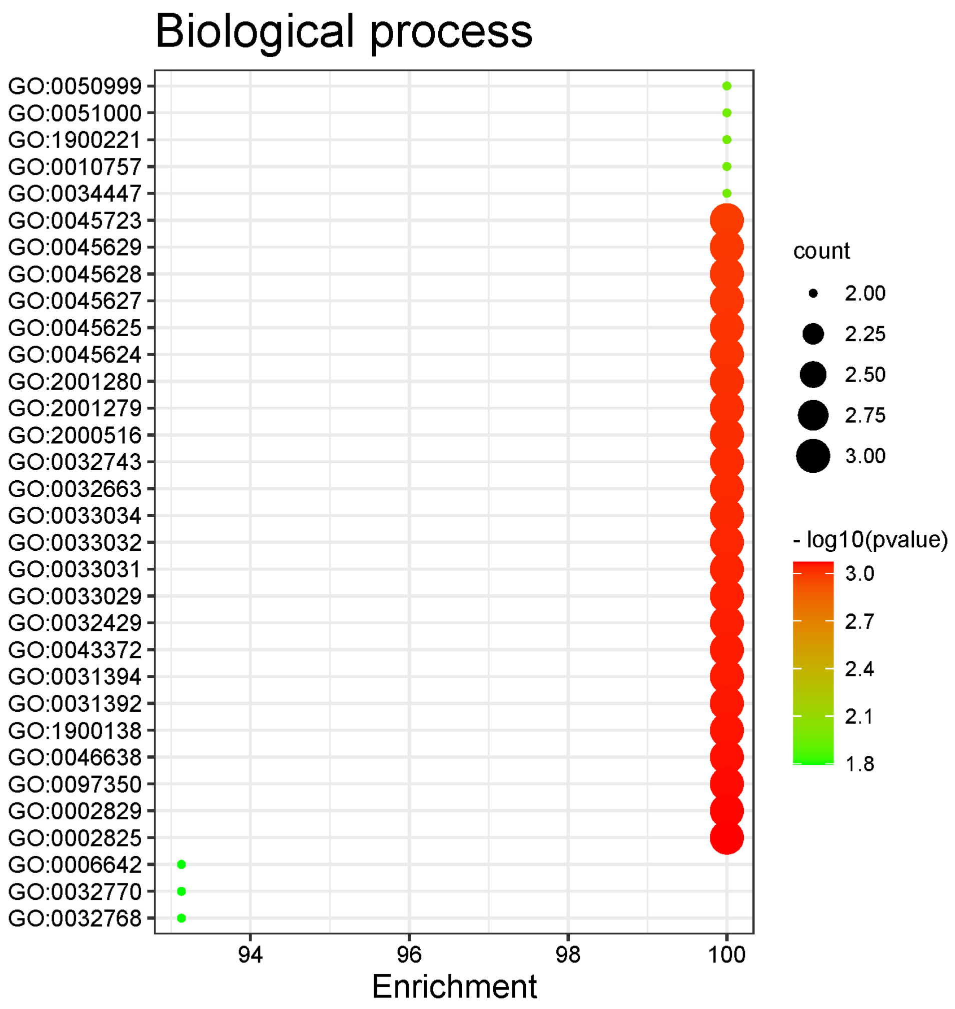


**Supplementary figure S6. Representation of biological process GO categories with higher enrichment in bone ECM proteins of *Danio rerio*.**

GO:0050999, regulation of nitric-oxide synthase activity; GO:0051000, positive regulation of nitric-oxide synthase activity; GO:1900221, regulation of amyloid-beta clearance; GO:0010757, negative regulation of plasminogen activation; GO:0034447, very-low-density lipoprotein particle clearance; GO:0045723, positive regulation of fatty acid biosynthetic process; GO:0045629, negative regulation of T-helper 2 cell differentiation; GO:0045628, regulation of T-helper 2 cell differentiation; GO:0045627, positive regulation of T-helper 1 cell differentiation; GO:0045625, regulation of T-helper 1 cell differentiation; GO:0045624, positive regulation of T-helper cell differentiation; GO:2001280, positive regulation of unsaturated fatty acid biosynthetic process; GO:2001279, regulation of unsaturated fatty acid biosynthetic process; GO:2000516, positive regulation of CD4-positive, alpha-beta T cell activation; GO:0032743, positive regulation of interleukin-2 production; GO:0032663, regulation of interleukin-2 production; GO:0033034, positive regulation of myeloid cell apoptotic process; GO:0033032, regulation of myeloid cell apoptotic process; GO:0033031, positive regulation of neutrophil apoptotic process; GO:0033029, regulation of neutrophil apoptotic process; GO:0032429, regulation of phospholipase A2 activity; GO:0043372, positive regulation of CD4-positive, alpha-beta T cell differentiation; GO:0031394, positive regulation of prostaglandin biosynthetic process; GO:0031392, regulation of prostaglandin biosynthetic process; GO:1900138, negative regulation of phospholipase A2 activity; GO:0046638, positive regulation of alpha-beta T cell differentiation; GO:0097350, neutrophil clearance; GO:0002829, negative regulation of type 2 immune response; GO:0002825, regulation of T-helper 1 type immune response; GO:0006642, triglyceride mobilization; GO:0032770, positive regulation of monooxygenase activity; and GO:0032768, regulation of monooxygenase activity. Points with Enrichment 100 mean 100 or more.

### SUPPLEMENTARY TABLES

**Supplementary table S1.** Proteins used as seed to build the Phylobone database.

| PB ID | Uniprot *D. rerio* | Gene name *D. rerio* | Uniprot *H. sapiens* | Gene name *H. sapiens* | Uniprot C. nippon | Gene Name C. nippon | Ref. |
|---|---|---|---|---|---|---|---|
| **PB0001** | A5WWI5 | ahsg | P02765 | AHSG | | | (1) |
| **PB0002** | B0UYS1 | b2m | P61769 | B2M | | | (1) |
| **PB0003** | E7FAN5 | f2 | P00734 | F2 | | | (1) |
| **PB0004** | Q6YND0 | mgp | P08493 | MGP | | | (1) |
| **PB0005** | B7ZDB5 | bgnb | P21810 | BGN | | BGN | (1) |
| **PB0005** | Q6GMI5 | bgna | P21810 | BGN | | BGN | (1) |
| **PB0006** | Q6PVV8 | sparc | P09486 | SPARC | | SPARC | (1) |
| **PB0007** | F1Q7R2 | chad | O15335 | CHAD | | | (1) |
| **PB0008** | F1R6R2 | APCS | P02743 | APCS | | | (1) |
| **PB0009** | Q66I20 | serpinf1 | P36955 | SERPINF1 | | | (1) |
| **PB0010** | | | P02792 | FTL | | | (1) |
| **PB0011** | | | P04406 | GAPDH | | | (1) |
| **PB0012** | | | P06702 | S100A9 | | | (1) |
| **PB0013** | | | P08311 | CTSG | | | (1) |
| **PB0014** | | | P24158 | PRTN3 | | | (1) |
| **PB0015** | | | P28300 | LOX | | LOX | (1) |
| **PB0016** | | | P62805 | HIST1H4A | | | (1) |
| **PB0017** | | | P69905 | HBA1 | | | (1) |
| **PB0018** | Q6IQX2 | col1a2 | P08123 | COL1A2 | | COL1A2 | (1) |
| **PB0019** | F1QDL1 | col1a1b | P02452 | COL1A1 | | COL1A1 | (1) |
| **PB0019** | F1QJC9 | col1a1a | P02452 | COL1A1 | | COL1A1 | (1) |
| **PB0020** | A0JML7 | adpgk | Q9BRR6 | ADPGK | | | (1) |
| **PB0021** | A1IGX5 | fibin | Q8TAL6 | FIBIN | | | (1) |
| **PB0022** | A1L1V4 | loxl2b | Q9Y4K0 | LOXL2 | | | (1) |
| **PB0023** | A2AVJ4 | bmp3 | P12645 | BMP3 | | | (1) |
| **PB0024** | A2CEN1 | efemp2a | Q6FH22 | EFEMP2 | | | (1) |
| **PB0025** | A2CEW3 | fn1b | P02751 | FN1 | | FN1 | (1) |
| **PB0026** | A2RRT1 | mfge8b | Q08431 | MFGE8 | | | (1) |
| **PB0027** | A2RUX5 | col11a1a | P12107 | COL11A1 | | COL11A1 | (1) |
| **PB0027** | D6MUD3 | col11a1b | P12107 | COL11A1 | | COL11A1 | (1) |
| **PB0028** | A3KNP6 | sfrp1b | Q8N474 | SFRP1 | | | (1) |
| **PB0028** | Q7T2K9 | sfrp1a | Q8N474 | SFRP1 | | | (1) |
| **PB0029** | A3KQQ9 | scg3 | Q8WXD2 | SCG3 | | | (1) |
| **PB0030** | A5PMP6 | c3b.1 | P01024 | C3 | | | (1) |
| **PB0030** | B8JKW4 | c3a | P01024 | C3 | | | (1) |
| **PB0030** | F1QX13 | c3c | P01024 | C3 | | | (1) |
| **PB0031** | A5WWJ4 | matn1 | P21941 | MATN1 | | | (1) |
| **PB0032** | A7MCK2 | hsd17b12b | Q53GQ0 | HSD17B12 | | | (1) |
| **PB0033** | A8E5G1 | anxa2b | P07355 | ANXA2 | | | (1) |
| **PB0034** | E2FHP6 | cfhl2 | P36980 | CFHR2 | | | (1) |
| **PB0035** | A8KC28 | proc | P04070 | PROC | | | (1) |
| **PB0035** | B8JLG2 | proc | P04070 | PROC | | | (1) |
| **PB0036** | A9JRB3 | htra1b | Q92743 | HTRA1 | | | (1) |
| **PB0036** | Q6GMI0 | htra1a | Q92743 | HTRA1 | | | (1) |
| **PB0037** | B0JZL7 | enpp1 | P22413 | ENPP1 | | | (1) |
| **PB0038** | B0S525 | igfbp5b | P24593 | IGFBP5 | | | (1) |
| **PB0039** | P58239 | lect1 | O75829 | LECT1 | | | (1) |

| | | | | | | |
|---|---|---|---|---|---|---|
| **PB0040** | B0S5W0 | col8a1a | P27658 | COL8A1 | | (1) |
| **PB0041** | B0S5X0 | epyc | Q99645 | EPYC | | (1) |
| **PB0042** | B0S6C1 | tfpia | P48307 | TFPI2 | | (1) |
| **PB0043** | B0S6K5 | tnn | Q9UQP3 | TNN | | (1) |
| **PB0044** | B0UXF7 | col4a5 | P29400 | COL4A5 | | (1) |
| **PB0045** | B0UXN0 | qsox1 | O00391 | QSOX1 | | (1) |
| **PB0046** | B0UXR7 | olfml2a | Q68BL7 | OLFML2A | | (1) |
| **PB0047** | B3DFP9 | apoa2 | P02652 | APOA2 | | (1) |
| **PB0048** | B3DFV8 | coch | O43405 | COCH | | (1) |
| **PB0049** | B3DHC5 | apoa4b.2 | P06727 | APOA4 | | (1) |
| **PB0050** | B3DJG2 | pcolce (2/2) | Q15113 | PCOLCE | | (1) |
| **PB0050** | F1QUX0 | pcolce (1/2) | Q15113 | PCOLCE | | (1) |
| **PB0051** | E7FG81 | col12a1 | Q99715 | COL12A1 | | (1) |
| **PB0051** | E7FG81 | col12a1 | Q99715 | COL12A1 | | (1) |
| **PB0052** | B7SDQ7 | ccdc80 | Q76M96 | CCDC80 | | (1) |
| **PB0053** | B7ZDA6 | lamb2 | P55268 | LAMB2 | | (1) |
| **PB0053** | F1QKX8 | lamb2l | P55268 | LAMB2P1 | | (1) |
| **PB0054** | B8A4T6 | col9a1 | P20849 | COL9A1 | | (1) |
| **PB0054** | F1QTQ2 | col9a1 | P20849 | COL9A1 | | (1) |
| **PB0055** | B8JHU4 | actn1 | P12814 | ACTN1 | | (1) |
| **PB0056** | E9QIM2 | sod3b | P08294 | SOD3 | | (1) |
| **PB0057** | B8JL29 | hmgb2a | P26583 | HMGB2 | | (1) |
| **PB0057** | Q66IB6 | hmgb2b | P26583 | HMGB2 | | (1) |
| **PB0058** | B8JL43 | tfa | P02787 | TF | | (1) |
| **PB0059** | B8JLZ3 | anxa1b | P04083 | ANXA1 | | (1) |
| **PB0059** | Q804H0 | anxa1c | P04083 | ANXA1 | | (1) |
| **PB0059** | Q804H2 | anxa1a | P04083 | ANXA1 | | (1) |
| **PB0060** | B8XY56 | rnaset2 | O00584 | RNASET2 | | (1) |
| **PB0061** | C1IHU8 | itln1 | Q8WWA0 | ITLN1 | | (1) |
| **PB0062** | E7F7K2 | c1qtnf9 | B2RNN3 | C1QTNF9B | | (1) |
| **PB0063** | E7EXE8 | tnxb | P22105 | TNXB | | (1) |
| **PB0064** | E7EYJ8 | clec3ba | P05452 | CLEC3B | | (1) |
| **PB0065** | E7F1S4 | thbs2b | P35442 | THBS2 | | (1) |
| **PB0065** | F8W5Z7 | thbs2a | P35442 | THBS2 | | (1) |
| **PB0066** | E7F2E3 | mfi2 | P08582 | MFI2 | | (1) |
| **PB0067** | E7F301 | f9a | P00740 | F9 | | (1) |
| **PB0067** | Q1RLV2 | f9b | P00740 | F9 | | (1) |
| **PB0068** | E7F537 | acana | P16112 | ACAN | | (1) |
| **PB0068** | F1R511 | acanb | P16112 | ACAN | | (1) |
| **PB0069** | E7F5I3 | postna | Q15063 | POSTN | POSTN | (1) |
| **PB0069** | Q75U66 | postnb | Q15063 | POSTN | POSTN | (1) |
| **PB0070** | X1WE42 | nid2 | Q14112 | NID2 | NID2 | (1) |
| **PB0071** | B8A4S4 | col17a1a | Q9UMD9 | COL17A1 | | (1) |
| **PB0072** | E7F829 | aebp1 | Q8IUX7 | AEBP1 | | (1) |
| **PB0073** | A0A286Y9F3 | otos | Q8NHW6 | OTOS | | (1) |
| **PB0074** | E7F8X0 | SRPX2 | O60687 | SRPX2 | | (1) |
| **PB0075** | F1RAG3 | nid1a | P14543 | NID1 | NID1 | (1) |
| **PB0076** | E7FB76 | col21a1 | Q96P44 | COL21A1 | | (1) |
| **PB0077** | E7FBD3 | apob | P04114 | APOB | | (1) |
| **PB0077** | Q5TZ29 | apobb | P04114 | APOB | | (1) |
| **PB0078** | E7FBW8 | hapln1b | P10915 | HAPLN1 | | (1) |
| **PB0078** | Q1LXE1 | hapln1a | P10915 | HAPLN1 | | (1) |
| **PB0079** | D6MUD6 | col5a3a | P25940 | COL5A3 | | (1) |
| **PB0079** | D6MUD7 | col5a3b | P25940 | COL5A3 | | (1) |

| | | | | | | |
|---|---|---|---|---|---|---|
| **PB0080** | E7FCV2 | c5 | P01031 | C5 | | (1) |
| **PB0081** | E7FCV8 | col6a2 | P12110 | COL6A2 | COL6A2 | (1) |
| **PB0082** | E7FDK4 | podn | Q7Z5L7 | PODN | | (1) |
| **PB0083** | X1WG18 | muc5b | Q9HC84 | MUC5B | | (1) |
| **PB0084** | E7FE47 | col5a2b | P05997 | COL5A2 | COL5A2 | (1) |
| **PB0085** | E7FE90 | fetub | Q9UGM5 | FETUB | | (1) |
| **PB0086** | A0A0R4IKF0 | apoa1b | P02647 | APOA1 | | (1) |
| **PB0087** | E9QDI1 | apoc1l | P02654 | APOC1 | | (1) |
| **PB0088** | E9QEQ1 | apoc2 | P02655 | APOC2 | | (1) |
| **PB0089** | E9QG22 | spp2 | Q13103 | SPP2 | | (1) |
| **PB0090** | F1Q4X1 | col6a3 | P12111 | COL6A3 | COL6A3 | (1) |
| **PB0091** | K7DYI0 | col16a1 | Q07092 | COL16A1 | | (1) |
| **PB0092** | F1Q5B5 | alpl | P05186 | ALPL | | (1) |
| **PB0093** | A2AR69 | itih6 | Q6UXX5 | ITIH6 | | (1) |
| **PB0094** | F1Q5N7 | cilp2 | Q8IUL8 | CILP2 | | (1) |
| **PB0095** | F1Q5S2 | proza | P22891 | PROZ | | (1) |
| **PB0095** | F1QZU6 | proz | P22891 | PROZ | | (1) |
| **PB0096** | F1Q6P3 | col6a1 | P12109 | COL6A1 | COL6A1 | (1) |
| **PB0097** | | | Q8N2E2 | VWDE | | (1) |
| **PB0098** | F1QG51 | fmoda | Q06828 | FMOD | | (1) |
| **PB0099** | F1Q775 | cilp | O75339 | CILP | | (1) |
| **PB0100** | F1Q8A0 | ctsk | P43235 | CTSK | | (1) |
| **PB0101** | X1WEZ4 | col6a4a | A6NMZ7 | col6a6 | | (1) |
| **PB0102** | F1Q9F4 | spp1 | P10451 | SPP1 | | (1) |
| **PB0103** | Q7ZV46 | metrnl | Q641Q3 | METRNL | | (1) |
| **PB0104** | A0A286Y830 | emid1 | Q96A84 | EMID1 | | (1) |
| **PB0104** | F1QR06 | emid1 | Q96A84 | EMID1 | | (1) |
| **PB0105** | Q8JHV6 | lamb4 | A4D0S4 | LAMB4 | | (1) |
| **PB0106** | F1QBE9 | ccl25b | O15444 | CCL25 | | (1) |
| **PB0107** | F1QC76 | mmp9 | P14780 | MMP9 | | (1) |
| **PB0108** | F1QEE7 | thbs1b | P07996 | THBS1 | THBS1 | (1) |
| **PB0109** | F1QFH8 | apom | O95445 | APOM | | (1) |
| **PB0110** | F1QFP3 | f7 | P08709 | F7 | | (1) |
| **PB0111** | F1QFX6 | ltbp3 | Q9NS15 | LTBP3 | | (1) |
| **PB0112** | Q71G59 | mmp13b | P45452 | MMP13 | | (1) |
| **PB0113** | F1QG79 | col4a1 | P02462 | COL4A1 | | (1) |
| **PB0114** | F1QGG9 | gas6 | Q14393 | GAS6 | | (1) |
| **PB0115** | F1QHI8 | pdgfrl | Q15198 | PDGFRL | | (1) |
| **PB0116** | | | P04004 | VTN | VTN | (1) |
| **PB0117** | F1QIJ3 | lamc2 | Q13753 | LAMC2 | | (1) |
| **PB0118** | F1QJB3 | c2 | P06681 | C2 | | (1) |
| **PB0119** | F1QJG6 | sfrp5 | Q5T4F7 | SFRP5 | | (1) |
| **PB0120** | F1QK29 | pros1 | P07225 | PROS1 | | (1) |
| **PB0121** | F1QK57 | tecta | O75443 | TECTA | | (1) |
| **PB0122** | F1QKL1 | thbs3a | P49746 | THBS3 | | (1) |
| **PB0123** | F1QKW3 | lama5 | O15230 | LAMA5 | | (1) |
| **PB0124** | F1QLC3 | f10 | P00742 | F10 | | (1) |
| **PB0125** | F1QLH9 | nucb1 | Q02818 | NUCB1 | | (1) |
| **PB0126** | F1QLT3 | fgfbp2 | Q9BYJ0 | FGFBP2 | | (1) |
| **PB0126** | F1R8L1 | fgfbp2 | Q9BYJ0 | FGFBP2 | | (1) |
| **PB0127** | F1QLW6 | ogn (2/2) | P20774 | OGN | | (1) |
| **PB0127** | Q1LV51 | ogn | P20774 | OGN | | (1) |
| **PB0128** | Q8JHV7 | lamb1a | P07942 | LAMB1 | LAMB1 | (1) |
| **PB0128** | F1REL9 | lamb1b | P07942 | LAMB1 | LAMB1 | (1) |

| | | | | | | | |
|---|---|---|---|---|---|---|---|
| **PB0129** | L0S5L0 | col2a1b | P02458 | COL2A1 | | | (1) |
| **PB0129** | Q2LDA1 | col2a1a | P02458 | COL2A1 | | | (1) |
| **PB0130** | F1QPX0 | ndnf | Q8TB73 | NDNF | | | (1) |
| **PB0131** | F1QQE4 | slit2 | O94813 | SLIT2 | | | (1) |
| **PB0132** | F1QRB8 | serp1 | P05121 | SERPINE1 | | | (1) |
| **PB0133** | B3DKC3 | mfap2 | P55001 | MFAP2 | | | (1) |
| **PB0134** | F1QTG2 | vwa2 | Q5GFL6 | VWA2 | | | (1) |
| **PB0135** | A0A2R8Q7H6 | lama3 | Q16787 | LAMA3 | | | (1) |
| **PB0136** | F1QW52 | habp2 | Q14520 | HABP2 | | | (1) |
| **PB0137** | F1QXD5 | col10a1 | Q03692 | COL10A1 | | | (1) |
| **PB0138** | F1QY29 | prelp | P51888 | PRELP | | | (1) |
| **PB0139** | F1QYE2 | tnc | P24821 | TNC | | TNC | (1) |
| **PB0140** | F1QYN1 | col4a6 | Q14031 | COL4A6 | | | (1) |
| **PB0141** | F1QYU0 | lama4 | Q16363 | LAMA4 | | LAMA4 | (1) |
| **PB0142** | F1R074 | agrn | O00468 | AGRN | | | (1) |
| **PB0143** | F1R0N0 | lrrc17 | Q8N6Y2 | LRRC17 | | | (1) |
| **PB0144** | F1R147 | pla2g12b | Q9BX93 | PLA2G12B | | | (1) |
| **PB0145** | F1R1F1 | col9a3 | Q14050 | COL9A3 | | | (1) |
| **PB0146** | F1R1P9 | thbs4b | P35443 | THBS4 | | | (1) |
| **PB0147** | A0A0H2UKP2 | col4a2 | P08572 | COL4A2 | | | (1) |
| **PB0148** | F1R2S9 | ecm2 | O94769 | ECM2 | | ECM2 | (1) |
| **PB0149** | F1R2W1 | dhrs13a.3 | Q6UX07 | DHRS13 | | | (1) |
| **PB0150** | F1R2Z9 | spon1b | Q9HCB6 | SPON1 | | | (1) |
| **PB0151** | D6MUD4 | col11a2 | P13942 | COL11A2 | | | (1) |
| **PB0152** | F1R5K7 | hspg2 | P98160 | HSPG2 | | HSPG2 | (1) |
| **PB0153** | F1R5V4 | col4a3 | Q01955 | COL4A3 | | | (1) |
| **PB0154** | F1R5Z2 | fndc1 | Q4ZHG4 | FNDC1 | | | (1) |
| **PB0155** | Q6IQQ7 | lum | P51884 | LUM | | LUM | (1) |
| **PB0156** | | | P35555 | FBN1 | | | (1) |
| **PB0157** | F1R774 | pcolce2b | Q9UKZ9 | PCOLCE2 | | | (1) |
| **PB0158** | Q5KQU0 | angptl1 | O95841 | ANGPTL1 | | | (1) |
| **PB0158** | Q5RH53 | angptl1 | O95841 | ANGPTL1 | | | (1) |
| **PB0159** | F1R886 | cfb | P00751 | CFB | | | (1) |
| **PB0160** | Q7SZV4 | tgfb2 | P61812 | TGFB2 | | | (1) |
| **PB0161** | E9QI62 | flna | P21333 | FLNA | | | (1) |
| **PB0162** | A0A0G2L5F9 | cfi | P05156 | CFI | | | (1) |
| **PB0163** | F1R9Y1 | col14a1 | Q05707 | COL14A1 | | COL14A1 | (1) |
| **PB0164** | A6H8K2 | bmper | Q8N8U9 | BMPER | | | (1) |
| **PB0165** | F1RAP6 | serpinf2b | P08697 | SERPINF2 | | | (1) |
| **PB0166** | Q9PWH3 | dkk1a | O94907 | DKK1 | | | (1) |
| **PB0167** | F6P711 | cpz | Q66K79 | CPZ | | | (1) |
| **PB0168** | F1RCV3 | slit3 | O75094 | SLIT3 | | | (1) |
| **PB0169** | F1RDU8 | FCGBP | Q9Y6R7 | FCGBP | | | (1) |
| **PB0170** | A0A0G2KQ62 | fbn2b | Q75N90 | FBN3 | | | (1) |
| **PB0171** | F1REI0 | htra3 | P83110 | HTRA3 | | | (1) |
| **PB0172** | A0A0R4IF38 | esm1 | Q9NQ30 | ESM1 | | | (1) |
| **PB0173** | F1REM3 | bmp1a | P13497 | BMP1 | | | (1) |
| **PB0174** | Q8AV83 | try | P07477 | PRSS1 | | | (1) |
| **PB0175** | Q5SPJ4 | serpina1 | P01009 | SERPINA1 | | | (1) |
| **PB0176** | Q503K1 | tgfbi | Q15582 | TGFBI | | TGFBI | (1) |
| **PB0177** | F2Z4S3 | tcn2l | P20062 | TCN2 | | | (1) |
| **PB0178** | O42222 | mstnb | O14793 | MSTN | | | (1) |
| **PB0179** | O42364 | apoeb | P02649 | APOE | | | (1) |
| **PB0179** | Q503V2 | apoea | P02649 | APOE | | | (1) |

| | | | | | | | |
|---|---|---|---|---|---|---|---|
| **PB0180** | Q15JE7 | optc | Q9UBM4 | OPTC | | | (1) |
| **PB0181** | Q1L8M3 | fgf23 | Q9GZV9 | FGF23 | | | (1) |
| **PB0182** | Q1L8P0 | crtap | O75718 | CRTAP | | | (1) |
| **PB0183** | Q1LUC6 | tinagl1 | Q9GZM7 | TINAGL1 | | | (1) |
| **PB0184** | Q1LVF0 | lamc1 | P11047 | LAMC1 | | LAMC1 | (1) |
| **PB0185** | Q1LXA7 | aspn | Q9BXN1 | ASPN | | | (1) |
| **PB0186** | Q1LXU2 | col9a2 | Q14055 | COL9A2 | | | (1) |
| **PB0187** | Q1LYJ7 | kng1 | P01042 | KNG1 | | | (1) |
| **PB0188** | Q1RLX2 | timp2a | P16035 | TIMP2 | | | (1) |
| **PB0189** | Q24JW2 | lyz | P61626 | LYZ | | | (1) |
| **PB0190** | Q29RB4 | olfml3a | Q9NRN5 | OLFML3 | | | (1) |
| **PB0191** | Q29RF0 | dpt | Q07507 | DPT | | | (1) |
| **PB0192** | Q2PMI2 | fstl1b | Q12841 | FSTL1 | | | (1) |
| **PB0193** | Q5F0G5 | myoc | Q99972 | MYOC | | | (1) |
| **PB0194** | Q5NJJ5 | matn4 | O95460 | MATN4 | | | (1) |
| **PB0195** | Q5NJJ7 | matn3b | O15232 | MATN3 | | | (1) |
| **PB0195** | Q68EK6 | matn3a | O15232 | MATN3 | | | (1) |
| **PB0196** | Q5RI33 | ctgfa | P29279 | CTGF | | | (1) |
| **PB0197** | Q5RI43 | kera | O60938 | KERA | | | (1) |
| **PB0198** | Q5RI45 | dcn | P07585 | DCN | | DCN | (1) |
| **PB0199** | Q5RIH8 | serpina10a | Q9UK55 | SERPINA10 | | | (1) |
| **PB0200** | Q5SPR2 | clu | P10909 | CLU | | | (1) |
| **PB0201** | Q66I23 | tgfb3 | P10600 | TGFB3 | | | (1) |
| **PB0202** | Q6AXL0 | cthrc1a | Q96CG8 | CTHRC1 | | | (1) |
| **PB0203** | Q6AZC0 | cela1 | Q9UNI1 | CELA1 | | | (1) |
| **PB0204** | Q6DG10 | mmp2 | P08253 | MMP2 | | MMP2 | (1) |
| **PB0205** | Q6IQP3 | calua | O43852 | CALU | | | (1) |
| **PB0206** | Q6NWB6 | ucmab | Q8WVF2 | UCMA | | | (1) |
| **PB0207** | Q6NXA5 | pon2 | Q15165 | PON2 | | | (1) |
| **PB0208** | Q6NYE1 | fgb | P02675 | FGB | | FGB | (1) |
| **PB0209** | Q6NYR4 | hspa8 | P11142 | HSPA8 | | | (1) |
| **PB0210** | | | P78380 | OLR1 | | | (1) |
| **PB0211** | Q6P0V8 | anxa5b | P08758 | ANXA5 | | | (1) |
| **PB0212** | Q6PC07 | rbp4l | P02753 | RBP4 | | | (1) |
| **PB0213** | Q6T937 | grem1a | O60565 | GREM1 | | | (1) |
| **PB0214** | Q7ZVG7 | fgg | P02679 | FGG | | FGG | (1) |
| **PB0215** | Q7ZVL5 | serpine2 | P07093 | SERPINE2 | | | (1) |
| **PB0216** | Q803H7 | itm2ba | Q9Y287 | ITM2B | | | (1) |
| **PB0217** | Q8QGV5 | ptgdsb | Q6UWW0 | LCN15 | | | (1) |
| **PB0218** | Q9DDG2 | mdkb | P21741 | MDK | | | (1) |
| **PB0218** | Q9W767 | mdka | P21741 | MDK | | | (1) |
| **PB0219** | Q9PTH3 | igfbp2a | P18065 | IGFBP2 | | | (1) |
| **PB0220** | Q9W6E0 | frzb | Q92765 | FRZB | | | (1) |
| **PB0221** | Q9YHV4 | fsta | P19883 | FST | | | |
| **PB0222** | | | P01023 | A2M | | A2M | (2) |
| **PB0223** | | | P05067 | APP | | APP | (2) |
| **PB0224** | | | P40121 | CAPG | | CAPG | (2) |
| **PB0225** | | | P07384 | CAPN1 | | CAPN1 | (2) |
| **PB0226** | | | P17655 | CAPN2 | | CAPN2 | (2) |
| **PB0227** | | | P04632 | CAPNS1 | | CAPNS1 | (2) |
| **PB0228** | | | P16070 | CD44 | | CD44 | (2) |
| **PB0229** | | | P39060 | COL18A1 | | COL18A1 | (2) |
| **PB0230** | | | P02461 | COL3A1 | | COL3A1 | (2) |
| **PB0231** | | | P23142 | FBLN1 | | FBLN1 | (2) |

| | | | | | |
|---|---|---|---|---|---|
| **PB0232** | P02671 | FGA | | FGA | (2) |
| **PB0233** | P06396 | GSN | | GSN | (2) |
| **PB0234** | P08648 | ITGA5 | | ITGA5 | (2) |
| **PB0235** | P06756 | ITGAV | | ITGAV | (2) |
| **PB0236** | P05556 | ITGB1 | | ITGB1 | (2) |
| **PB0237** | P13796 | LCP1 | | LCP1 | (2) |
| **PB0238** | P00747 | PLG | | PLG | (2) |
| **PB0239** | Q13162 | PRDX4 | | PRDX4 | (2) |
| **PB0240** | P36952 | SERPINB5 | | SERPINB5 | (2) |
| **PB0241** | A1X283 | SH3PXD2B | | SH3PXD2B | (2) |
| **PB0242** | P02766 | TTR | | TTR | (2) |
| **PB0243** | P13611 | VCAN | | VCAN | (2) |
| **PB0244** | Q15942 | ZYX | | ZYX | (2) |
| **PB0245** | P35222 | CTNNB1 | | CTNNB1 | (2) |
| **PB0246** | P63000 | RAC1 | | RAC1 | (2) |
| **PB0247** | Q96AC1 | FERMT2 | | FERMT2 | (2) |
| **PB0248** | P18206 | VCL | | VCL | (2) |
| **PB0249** | P21815 | IBSP | E7D7Z0 | BSPII | (3) |
| **PB0250** | Q99217 | AMELX | | | (4) |
| **PB0251** | P12644 | BMP4 | | | (5) |
| **PB0252** | Q2MKA7 | RSPO1 | | | (6) |
| **PB0253** | Q6UXX9 | RSPO2 | | | (6) |
| **PB0254** | Q9BXY4 | RSPO3 | | | (6) |
| **PB0255** | Q2I0M5 | RSPO4 | | | (6) |

**Supplementary table S2. List of selected species included in the Phylobone database.**

| N | Species | | Taxonomic group | | Proteins |
|---|---|---|---|---|---|
| 1 | *Homo sapiens (*)* | Human | Vertebrates | Primates | 255 |
| 2 | *Pan troglodytes* | Chimpanzee | Vertebrates | Primates | 255 |
| 3 | *Pan paniscus* | Pygmy chimpanzee | Vertebrates | Primates | 255 |
| 4 | *Gorilla gorilla* | Western gorilla | Vertebrates | Primates | 247 |
| 5 | *Pongo abelii* | Sumatran orangutan | Vertebrates | Primates | 255 |
| 6 | *Oryctolagus cuniculus* | Rabbit | Vertebrates | Rabbits | 244 |
| 7 | *Mus musculus* | Mouse | Vertebrates | Rodents | 253 |
| 8 | *Rattus norvegicus* | Rat | Vertebrates | Rodents | 253 |
| 9 | *Felis catus* | Cat | Vertebrates | Carnivores | 250 |
| 10 | *Canis lupus familiaris* | Dog | Vertebrates | Carnivores | 250 |
| 11 | *Cervus elaphus* | Red deer | Vertebrates | Even-toed ungulates | 168 |
| 12 | *Cervus hanglu* | Central Asian red deer | Vertebrates | Even-toed ungulates | 171 |
| 13 | *Cervus canadensis* | Elk | Vertebrates | Even-toed ungulates | 245 |
| 14 | *Odocoileus virginianus* | White-tailed deer | Vertebrates | Even-toed ungulates | 243 |
| 15 | *Muntiacus muntjak* | Indian muntjac | Vertebrates | Even-toed ungulates | 202 |
| 16 | *Muntiacus reevesi* | Chinese muntjac | Vertebrates | Even-toed ungulates | 186 |
| 17 | *Capra hircus* | Goat | Vertebrates | Even-toed ungulates | 251 |
| 18 | *Ovis aries* | Sheep | Vertebrates | Even-toed ungulates | 251 |
| 19 | *Bos taurus* | Bull | Vertebrates | Even-toed ungulates | 253 |
| 20 | *Sus scrofa* | Pig | Vertebrates | Even-toed ungulates | 251 |
| 21 | *Alligator mississippiensis* | American alligator | Vertebrates | Reptiles | 243 |
| 22 | *Alligator sinensis* | Chinese alligator | Vertebrates | Reptiles | 238 |

| | | | | | |
|---|---|---|---|---|---|
| 23 | *Columba livia* | Rock pigeon | Vertebrates | Birds | 224 |
| 24 | *Gallus gallus* | Chicken | Vertebrates | Birds | 244 |
| 25 | *Meleagris gallopavo* | Turkey | Vertebrates | Birds | 216 |
| 26 | *Xenopus laevis* | African clawed frog | Vertebrates | Frogs and toads | 248 |
| 27 | *Rana temporaria* | common frog | Vertebrates | Frogs and toads | 245 |
| 28 | *Danio rerio (*)* | Zebrafish | Vertebrates | Bony fishes | 251 |
| 29 | *Nothobranchius furzeri* | Turquoise killifish | Vertebrates | Bony fishes | 245 |
| 30 | *Oryzias latipes* | Japanese medaka | Vertebrates | Bony fishes | 246 |
| 31 | *Sparus aurata* | Gilthead seabream | Vertebrates | Bony fishes | 249 |
| 32 | *Styela clava* | Asian tunicate | Chordata | Tunicata | 188 |
| 33 | *Patiria miniata* | Bat star | Invertebrates | Echinoderm | 179 |
| 34 | *Strongylocentrotus purpuratus* | Purple sea urchin | Invertebrates | Echinoderm | 182 |
| 35 | *Drosophila melanogaster* | Fruit fly | Invertebrates | Insects | 135 |
| 36 | *Apis mellifera* | Honey bee | Invertebrates | Insects | 151 |
| 37 | *Caenorhabditis elegans* | Worm | Invertebrates | Nematoda | 115 |
| 38 | *Sepia pharaonis* | Cuttlefish | Invertebrates | Cephalopoda | 120 |
| 39 | *Actinia tenebrosa* | Anemone | Invertebrates | Cnidaria | 158 |

*(*) Used as seed to identify proteins in other organisms*

**Supplementary table S3. Phyletic patterns.**

| Groups | Pr | Ra | Ro | Ca | De | Bo | Pi | Re | Bi | Am | Fi | Tu | Ec | In | Ne | Ce | Cn |
|---|---|---|---|---|---|---|---|---|---|---|---|---|---|---|---|---|---|
| N | 5 | 1 | 2 | 2 | 6 | 3 | 1 | 2 | 3 | 2 | 4 | 1 | 2 | 2 | 1 | 1 | 1 |
| PB0001 | 1 | 1 | 1 | 1 | 1 | 1 | 1 | 1 | 1 | 1 | 1 | 0 | 0 | 0 | 0 | 0 | 0 |
| PB0002 | 1 | 1 | 1 | 1 | 1 | 1 | 1 | 1 | 1 | 1 | 1 | 0 | 0 | 0 | 0 | 0 | 0 |
| PB0003 | 1 | 1 | 1 | 1 | 1 | 1 | 1 | 1 | 1 | 1 | 1 | 1 | 1 | 1 | 1 | 1 | 1 |
| PB0004 | 1 | 1 | 1 | 1 | 1 | 1 | 1 | 1 | 1 | 1 | 1 | 0 | 0 | 0 | 0 | 0 | 0 |
| PB0005 | 1 | 1 | 1 | 1 | 1 | 1 | 1 | 1 | 1 | 1 | 1 | 1 | 1 | 1 | 1 | 1 | 0 |
| PB0006 | 1 | 1 | 1 | 1 | 1 | 1 | 1 | 1 | 1 | 1 | 1 | 1 | 1 | 1 | 1 | 0 | 1 |
| PB0007 | 1 | 1 | 1 | 1 | 1 | 1 | 1 | 1 | 1 | 1 | 1 | 1 | 1 | 1 | 1 | 1 | 1 |
| PB0008 | 1 | 1 | 1 | 1 | 1 | 1 | 1 | 1 | 1 | 1 | 1 | 1 | 1 | 1 | 0 | 0 | 1 |
| PB0009 | 1 | 1 | 1 | 1 | 1 | 1 | 1 | 1 | 1 | 1 | 1 | 1 | 1 | 1 | 1 | 0 | 1 |
| PB0010 | 1 | 1 | 1 | 1 | 1 | 1 | 1 | 1 | 1 | 1 | 1 | 1 | 1 | 1 | 1 | 1 | 1 |
| PB0011 | 1 | 1 | 1 | 1 | 1 | 1 | 1 | 1 | 1 | 1 | 1 | 1 | 1 | 1 | 1 | 1 | 1 |
| PB0012 | 1 | 1 | 1 | 1 | 1 | 1 | 1 | 1 | 1 | 1 | 1 | 0 | 0 | 0 | 0 | 0 | 0 |
| PB0013 | 1 | 1 | 1 | 1 | 1 | 1 | 1 | 1 | 1 | 1 | 1 | 1 | 1 | 1 | 1 | 1 | 1 |
| PB0014 | 1 | 1 | 1 | 1 | 1 | 1 | 1 | 1 | 1 | 1 | 1 | 1 | 1 | 1 | 1 | 1 | 1 |
| PB0015 | 1 | 1 | 1 | 1 | 1 | 1 | 1 | 1 | 1 | 1 | 1 | 1 | 1 | 1 | 0 | 1 | 1 |
| PB0016 | 1 | 1 | 1 | 1 | 1 | 1 | 1 | 1 | 1 | 1 | 1 | 1 | 1 | 1 | 1 | 0 | 1 |
| PB0017 | 1 | 1 | 1 | 1 | 1 | 1 | 1 | 1 | 1 | 1 | 1 | 0 | 0 | 0 | 0 | 0 | 0 |
| PB0018 | 1 | 1 | 1 | 1 | 1 | 1 | 1 | 1 | 1 | 1 | 1 | 1 | 1 | 1 | 0 | 1 | 1 |
| PB0019 | 1 | 1 | 1 | 1 | 1 | 1 | 1 | 1 | 1 | 1 | 1 | 1 | 1 | 1 | 0 | 1 | 1 |
| PB0020 | 1 | 1 | 1 | 1 | 1 | 1 | 1 | 1 | 1 | 1 | 1 | 1 | 1 | 1 | 1 | 1 | 1 |
| PB0021 | 1 | 1 | 1 | 1 | 1 | 1 | 1 | 0 | 1 | 1 | 1 | 0 | 0 | 0 | 0 | 0 | 0 |
| PB0022 | 1 | 1 | 1 | 1 | 1 | 1 | 1 | 1 | 1 | 1 | 1 | 1 | 1 | 1 | 0 | 1 | 1 |
| PB0023 | 1 | 1 | 1 | 1 | 1 | 1 | 1 | 1 | 1 | 1 | 1 | 1 | 1 | 1 | 1 | 1 | 1 |
| PB0024 | 1 | 1 | 1 | 1 | 1 | 1 | 1 | 1 | 1 | 1 | 1 | 1 | 1 | 1 | 1 | 1 | 1 |
| PB0025 | 1 | 1 | 1 | 1 | 1 | 1 | 1 | 1 | 1 | 1 | 1 | 0 | 0 | 0 | 0 | 0 | 0 |
| PB0026 | 1 | 1 | 1 | 1 | 1 | 1 | 1 | 1 | 1 | 1 | 1 | 1 | 1 | 0 | 0 | 1 | 0 |
| PB0027 | 1 | 1 | 1 | 1 | 1 | 1 | 1 | 1 | 1 | 1 | 1 | 1 | 1 | 1 | 1 | 1 | 1 |
| PB0028 | 1 | 1 | 1 | 1 | 1 | 1 | 1 | 1 | 1 | 1 | 1 | 1 | 1 | 1 | 1 | 1 | 1 |
| PB0029 | 1 | 1 | 1 | 1 | 1 | 1 | 1 | 1 | 1 | 1 | 1 | 0 | 0 | 0 | 0 | 0 | 0 |

| | | | | | | | | | | | | | | | | | |
|---|---|---|---|---|---|---|---|---|---|---|---|---|---|---|---|---|---|
| PB0030 | 1 | 1 | 1 | 1 | 1 | 1 | 1 | 1 | 1 | 1 | 1 | 1 | 1 | 1 | 0 | 1 | 1 |
| PB0031 | 1 | 1 | 1 | 1 | 1 | 1 | 1 | 1 | 1 | 1 | 1 | 0 | 0 | 0 | 0 | 0 | 0 |
| PB0032 | 1 | 1 | 1 | 1 | 1 | 1 | 1 | 1 | 1 | 1 | 1 | 1 | 1 | 1 | 1 | 1 | 1 |
| PB0033 | 1 | 1 | 1 | 1 | 1 | 1 | 1 | 1 | 1 | 1 | 1 | 1 | 1 | 1 | 1 | 1 | 1 |
| PB0034 | 1 | 1 | 1 | 1 | 1 | 1 | 1 | 1 | 1 | 1 | 1 | 0 | 1 | 0 | 0 | 0 | 1 |
| PB0035 | 1 | 1 | 1 | 1 | 1 | 1 | 1 | 1 | 1 | 1 | 1 | 0 | 1 | 1 | 1 | 1 | 1 |
| PB0036 | 1 | 1 | 1 | 1 | 1 | 1 | 1 | 1 | 1 | 1 | 1 | 1 | 1 | 1 | 0 | 1 | 0 |
| PB0037 | 1 | 1 | 1 | 1 | 1 | 1 | 1 | 1 | 1 | 1 | 1 | 1 | 1 | 1 | 1 | 1 | 1 |
| PB0038 | 1 | 1 | 1 | 1 | 1 | 1 | 1 | 1 | 1 | 1 | 1 | 1 | 0 | 0 | 0 | 0 | 0 |
| PB0039 | 1 | 1 | 1 | 1 | 1 | 1 | 1 | 1 | 1 | 1 | 1 | 1 | 0 | 0 | 0 | 0 | 0 |
| PB0040 | 1 | 1 | 1 | 1 | 1 | 1 | 1 | 1 | 1 | 1 | 1 | 0 | 1 | 0 | 0 | 0 | 0 |
| PB0041 | 1 | 1 | 1 | 1 | 1 | 1 | 1 | 1 | 1 | 1 | 1 | 0 | 1 | 0 | 0 | 0 | 0 |
| PB0042 | 1 | 1 | 1 | 1 | 1 | 1 | 1 | 1 | 1 | 1 | 1 | 0 | 0 | 0 | 0 | 0 | 1 |
| PB0043 | 1 | 1 | 1 | 1 | 1 | 1 | 1 | 1 | 1 | 1 | 1 | 1 | 1 | 1 | 1 | 1 | 1 |
| PB0044 | 1 | 1 | 0 | 1 | 1 | 1 | 1 | 1 | 1 | 1 | 1 | 1 | 1 | 1 | 1 | 1 | 1 |
| PB0045 | 1 | 1 | 1 | 1 | 1 | 1 | 1 | 1 | 1 | 1 | 1 | 1 | 1 | 1 | 1 | 1 | 1 |
| PB0046 | 1 | 1 | 1 | 1 | 1 | 1 | 1 | 1 | 1 | 1 | 1 | 1 | 1 | 1 | 1 | 0 | 0 |
| PB0047 | 1 | 1 | 1 | 1 | 1 | 1 | 1 | 1 | 1 | 0 | 1 | 0 | 0 | 0 | 0 | 0 | 0 |
| PB0048 | 1 | 1 | 1 | 1 | 1 | 1 | 1 | 1 | 1 | 1 | 1 | 0 | 0 | 0 | 1 | 0 | 0 |
| PB0049 | 1 | 1 | 1 | 1 | 1 | 1 | 1 | 1 | 1 | 1 | 1 | 0 | 0 | 0 | 0 | 0 | 0 |
| PB0050 | 1 | 1 | 1 | 1 | 1 | 1 | 1 | 1 | 1 | 1 | 1 | 1 | 1 | 1 | 1 | 1 | 1 |
| PB0051 | 1 | 1 | 1 | 1 | 1 | 1 | 1 | 1 | 1 | 1 | 1 | 1 | 1 | 1 | 0 | 1 | 1 |
| PB0052 | 1 | 1 | 1 | 1 | 1 | 1 | 1 | 1 | 1 | 1 | 1 | 0 | 0 | 0 | 0 | 0 | 0 |
| PB0053 | 1 | 1 | 1 | 1 | 1 | 1 | 1 | 1 | 1 | 1 | 1 | 1 | 1 | 1 | 1 | 1 | 1 |
| PB0054 | 1 | 1 | 1 | 1 | 1 | 1 | 1 | 1 | 1 | 1 | 1 | 1 | 1 | 1 | 1 | 1 | 1 |
| PB0055 | 1 | 1 | 1 | 1 | 1 | 1 | 1 | 1 | 1 | 1 | 1 | 1 | 1 | 1 | 1 | 0 | 1 |
| PB0056 | 1 | 1 | 1 | 1 | 1 | 1 | 1 | 1 | 1 | 1 | 1 | 1 | 1 | 1 | 1 | 1 | 1 |
| PB0057 | 1 | 1 | 1 | 1 | 1 | 1 | 1 | 1 | 1 | 1 | 1 | 1 | 1 | 1 | 1 | 1 | 1 |
| PB0058 | 1 | 1 | 1 | 1 | 1 | 1 | 1 | 1 | 1 | 1 | 1 | 1 | 1 | 1 | 0 | 1 | 1 |
| PB0059 | 1 | 1 | 1 | 1 | 1 | 1 | 1 | 1 | 1 | 1 | 1 | 1 | 1 | 1 | 1 | 1 | 1 |
| PB0060 | 1 | 1 | 1 | 1 | 1 | 1 | 1 | 1 | 1 | 1 | 1 | 1 | 1 | 1 | 1 | 1 | 1 |
| PB0061 | 1 | 1 | 1 | 0 | 1 | 1 | 1 | 1 | 1 | 1 | 1 | 1 | 0 | 0 | 0 | 0 | 1 |
| PB0062 | 1 | 1 | 1 | 1 | 1 | 1 | 1 | 1 | 1 | 1 | 1 | 1 | 0 | 0 | 0 | 0 | 0 |
| PB0063 | 1 | 1 | 1 | 1 | 1 | 1 | 1 | 1 | 1 | 1 | 1 | 0 | 1 | 1 | 1 | 1 | 0 |
| PB0064 | 1 | 1 | 1 | 1 | 1 | 1 | 1 | 1 | 1 | 1 | 1 | 0 | 1 | 0 | 1 | 0 | 1 |
| PB0065 | 1 | 1 | 1 | 1 | 1 | 1 | 1 | 1 | 1 | 1 | 1 | 1 | 1 | 1 | 1 | 1 | 1 |
| PB0066 | 1 | 1 | 1 | 1 | 1 | 1 | 1 | 1 | 1 | 1 | 1 | 1 | 1 | 1 | 0 | 1 | 1 |
| PB0067 | 1 | 1 | 1 | 1 | 1 | 1 | 1 | 1 | 1 | 1 | 1 | 1 | 1 | 1 | 1 | 1 | 1 |
| PB0068 | 1 | 1 | 1 | 1 | 1 | 1 | 1 | 1 | 1 | 1 | 1 | 0 | 1 | 0 | 0 | 0 | 0 |
| PB0069 | 1 | 1 | 1 | 1 | 1 | 1 | 1 | 1 | 1 | 1 | 1 | 0 | 1 | 1 | 1 | 1 | 1 |
| PB0070 | 1 | 1 | 1 | 1 | 1 | 1 | 1 | 1 | 1 | 1 | 1 | 1 | 1 | 1 | 1 | 1 | 1 |
| PB0071 | 1 | 1 | 1 | 1 | 1 | 1 | 1 | 1 | 1 | 1 | 1 | 0 | 0 | 0 | 0 | 0 | 0 |
| PB0072 | 1 | 1 | 1 | 1 | 1 | 1 | 1 | 1 | 1 | 1 | 1 | 1 | 1 | 1 | 1 | 1 | 1 |
| PB0073 | 1 | 0 | 1 | 1 | 1 | 1 | 1 | 1 | 1 | 1 | 1 | 0 | 0 | 0 | 0 | 0 | 0 |
| PB0074 | 1 | 1 | 1 | 1 | 1 | 1 | 1 | 1 | 1 | 1 | 1 | 1 | 1 | 1 | 1 | 1 | 1 |
| PB0075 | 1 | 1 | 1 | 1 | 1 | 1 | 1 | 1 | 1 | 1 | 1 | 1 | 1 | 1 | 1 | 1 | 1 |
| PB0076 | 1 | 1 | 1 | 1 | 1 | 1 | 1 | 1 | 1 | 1 | 1 | 1 | 1 | 1 | 1 | 1 | 1 |
| PB0077 | 1 | 1 | 1 | 1 | 1 | 1 | 1 | 1 | 1 | 1 | 1 | 1 | 1 | 1 | 0 | 1 | 1 |
| PB0078 | 1 | 1 | 1 | 1 | 1 | 1 | 1 | 1 | 1 | 1 | 1 | 0 | 0 | 0 | 0 | 0 | 1 |
| PB0079 | 1 | 1 | 1 | 1 | 1 | 1 | 1 | 1 | 1 | 1 | 1 | 1 | 1 | 1 | 0 | 1 | 1 |
| PB0080 | 1 | 1 | 1 | 1 | 1 | 1 | 1 | 1 | 1 | 1 | 1 | 1 | 1 | 1 | 1 | 1 | 1 |
| PB0081 | 1 | 1 | 1 | 1 | 1 | 1 | 1 | 1 | 1 | 1 | 1 | 1 | 0 | 0 | 0 | 0 | 0 |
| PB0082 | 1 | 1 | 1 | 1 | 1 | 1 | 1 | 1 | 1 | 1 | 1 | 1 | 1 | 1 | 1 | 1 | 1 |
| PB0083 | 1 | 0 | 1 | 1 | 1 | 1 | 1 | 1 | 1 | 1 | 1 | 1 | 1 | 1 | 1 | 1 | 1 |
| PB0084 | 1 | 1 | 1 | 1 | 1 | 1 | 1 | 1 | 1 | 1 | 1 | 1 | 1 | 1 | 0 | 1 | 1 |
| PB0085 | 1 | 1 | 1 | 1 | 1 | 1 | 1 | 1 | 1 | 1 | 1 | 0 | 0 | 0 | 0 | 0 | 0 |
| PB0086 | 1 | 1 | 1 | 1 | 1 | 1 | 1 | 1 | 1 | 1 | 1 | 0 | 0 | 0 | 0 | 0 | 0 |
| PB0087 | 1 | 1 | 1 | 1 | 0 | 0 | 0 | 1 | 1 | 0 | 1 | 0 | 0 | 0 | 0 | 0 | 0 |

| | | | | | | | | | | | | | | | | | |
|---|---|---|---|---|---|---|---|---|---|---|---|---|---|---|---|---|---|
| PB0088 | 1 | 1 | 1 | 1 | 1 | 1 | 1 | 0 | 0 | 1 | 1 | 0 | 0 | 0 | 0 | 0 | 0 |
| PB0089 | 1 | 1 | 1 | 1 | 1 | 1 | 1 | 1 | 1 | 1 | 1 | 0 | 0 | 0 | 0 | 0 | 0 |
| PB0090 | 1 | 1 | 1 | 1 | 1 | 1 | 1 | 1 | 1 | 1 | 1 | 1 | 0 | 0 | 0 | 1 | 0 |
| PB0091 | 1 | 1 | 1 | 1 | 1 | 1 | 1 | 1 | 1 | 1 | 1 | 0 | 1 | 1 | 0 | 0 | 0 |
| PB0092 | 1 | 1 | 1 | 1 | 1 | 1 | 1 | 1 | 1 | 1 | 1 | 1 | 1 | 1 | 0 | 0 | 1 |
| PB0093 | 1 | 1 | 1 | 1 | 1 | 1 | 1 | 1 | 1 | 1 | 1 | 1 | 1 | 0 | 0 | 1 | 0 |
| PB0094 | 1 | 1 | 1 | 1 | 1 | 1 | 1 | 1 | 1 | 1 | 1 | 1 | 0 | 0 | 0 | 0 | 0 |
| PB0095 | 1 | 1 | 1 | 0 | 1 | 1 | 0 | 1 | 1 | 1 | 1 | 1 | 1 | 1 | 0 | 0 | 1 |
| PB0096 | 1 | 1 | 1 | 1 | 1 | 1 | 1 | 1 | 1 | 1 | 1 | 1 | 0 | 0 | 0 | 0 | 1 |
| PB0097 | 1 | 0 | 1 | 1 | 1 | 1 | 1 | 1 | 1 | 1 | 1 | 1 | 1 | 1 | 1 | 1 | 1 |
| PB0098 | 1 | 1 | 1 | 1 | 1 | 1 | 1 | 1 | 1 | 1 | 1 | 1 | 1 | 1 | 1 | 1 | 1 |
| PB0099 | 1 | 1 | 1 | 1 | 1 | 1 | 1 | 1 | 1 | 1 | 1 | 1 | 0 | 0 | 0 | 0 | 0 |
| PB0100 | 1 | 1 | 1 | 1 | 1 | 1 | 1 | 1 | 1 | 1 | 1 | 1 | 1 | 1 | 1 | 1 | 1 |
| PB0101 | 1 | 1 | 1 | 1 | 1 | 1 | 1 | 1 | 1 | 1 | 1 | 1 | 0 | 0 | 0 | 1 | 1 |
| PB0102 | 1 | 1 | 1 | 1 | 1 | 1 | 1 | 1 | 1 | 0 | 1 | 0 | 0 | 0 | 0 | 0 | 0 |
| PB0103 | 1 | 1 | 1 | 1 | 1 | 1 | 1 | 1 | 1 | 1 | 1 | 1 | 1 | 1 | 0 | 0 | 1 |
| PB0104 | 1 | 1 | 1 | 1 | 1 | 1 | 1 | 1 | 1 | 1 | 1 | 1 | 1 | 1 | 1 | 1 | 1 |
| PB0105 | 1 | 1 | 1 | 1 | 1 | 1 | 1 | 1 | 1 | 1 | 1 | 1 | 1 | 1 | 1 | 1 | 1 |
| PB0106 | 1 | 1 | 1 | 1 | 1 | 1 | 1 | 1 | 1 | 1 | 1 | 0 | 0 | 0 | 0 | 0 | 0 |
| PB0107 | 1 | 1 | 1 | 1 | 1 | 1 | 1 | 1 | 1 | 1 | 1 | 1 | 1 | 1 | 1 | 1 | 1 |
| PB0108 | 1 | 1 | 1 | 1 | 1 | 1 | 1 | 1 | 1 | 1 | 1 | 1 | 1 | 1 | 0 | 1 | 1 |
| PB0109 | 1 | 1 | 1 | 1 | 1 | 1 | 1 | 1 | 1 | 1 | 1 | 0 | 0 | 0 | 0 | 0 | 0 |
| PB0110 | 1 | 1 | 1 | 1 | 1 | 1 | 1 | 1 | 1 | 1 | 1 | 1 | 1 | 1 | 1 | 1 | 1 |
| PB0111 | 1 | 1 | 1 | 1 | 1 | 1 | 1 | 1 | 1 | 1 | 1 | 1 | 1 | 1 | 0 | 1 | 1 |
| PB0112 | 1 | 1 | 1 | 1 | 1 | 1 | 1 | 1 | 1 | 1 | 1 | 1 | 1 | 1 | 1 | 1 | 1 |
| PB0113 | 1 | 1 | 1 | 1 | 1 | 1 | 1 | 1 | 1 | 1 | 1 | 1 | 1 | 1 | 1 | 1 | 1 |
| PB0114 | 1 | 1 | 1 | 1 | 1 | 1 | 1 | 1 | 1 | 1 | 1 | 1 | 0 | 0 | 0 | 0 | 1 |
| PB0115 | 1 | 1 | 1 | 1 | 1 | 1 | 1 | 1 | 1 | 1 | 1 | 0 | 0 | 0 | 0 | 0 | 0 |
| PB0116 | 1 | 1 | 1 | 1 | 1 | 1 | 1 | 1 | 1 | 1 | 1 | 1 | 1 | 1 | 1 | 1 | 1 |
| PB0117 | 1 | 1 | 1 | 1 | 1 | 1 | 1 | 1 | 1 | 1 | 1 | 1 | 1 | 1 | 1 | 1 | 1 |
| PB0118 | 1 | 1 | 1 | 1 | 1 | 1 | 1 | 1 | 1 | 1 | 1 | 1 | 1 | 0 | 0 | 0 | 1 |
| PB0119 | 1 | 1 | 1 | 1 | 1 | 1 | 1 | 1 | 1 | 1 | 1 | 1 | 1 | 1 | 1 | 1 | 1 |
| PB0120 | 1 | 1 | 1 | 1 | 1 | 1 | 1 | 1 | 1 | 1 | 1 | 1 | 0 | 0 | 0 | 0 | 0 |
| PB0121 | 1 | 1 | 1 | 1 | 1 | 1 | 1 | 1 | 1 | 1 | 1 | 1 | 1 | 0 | 0 | 0 | 0 |
| PB0122 | 1 | 1 | 1 | 1 | 1 | 1 | 1 | 1 | 1 | 1 | 1 | 1 | 1 | 1 | 0 | 1 | 1 |
| PB0123 | 1 | 1 | 1 | 1 | 1 | 1 | 1 | 1 | 1 | 1 | 1 | 1 | 1 | 1 | 1 | 0 | 1 |
| PB0124 | 1 | 1 | 1 | 1 | 1 | 1 | 1 | 1 | 1 | 1 | 1 | 1 | 1 | 1 | 1 | 1 | 1 |
| PB0125 | 1 | 1 | 1 | 1 | 1 | 1 | 1 | 1 | 1 | 1 | 1 | 1 | 1 | 1 | 1 | 0 | 1 |
| PB0126 | 1 | 1 | 0 | 0 | 0 | 0 | 0 | 1 | 1 | 1 | 1 | 0 | 0 | 0 | 0 | 0 | 0 |
| PB0127 | 1 | 1 | 1 | 1 | 1 | 1 | 1 | 1 | 1 | 1 | 1 | 0 | 1 | 1 | 0 | 0 | 0 |
| PB0128 | 1 | 1 | 1 | 1 | 1 | 1 | 1 | 1 | 1 | 1 | 1 | 1 | 1 | 1 | 1 | 1 | 1 |
| PB0129 | 1 | 1 | 1 | 1 | 1 | 1 | 1 | 1 | 1 | 1 | 1 | 1 | 1 | 1 | 1 | 1 | 1 |
| PB0130 | 1 | 1 | 1 | 1 | 1 | 1 | 1 | 1 | 1 | 1 | 1 | 0 | 1 | 1 | 1 | 0 | 0 |
| PB0131 | 1 | 1 | 1 | 1 | 1 | 1 | 1 | 1 | 1 | 1 | 1 | 1 | 1 | 1 | 1 | 0 | 0 |
| PB0132 | 1 | 1 | 1 | 1 | 1 | 1 | 1 | 1 | 1 | 1 | 1 | 1 | 1 | 1 | 1 | 0 | 1 |
| PB0133 | 1 | 0 | 1 | 1 | 1 | 1 | 1 | 1 | 1 | 1 | 1 | 0 | 0 | 0 | 0 | 0 | 0 |
| PB0134 | 1 | 1 | 1 | 1 | 1 | 1 | 1 | 1 | 1 | 1 | 1 | 1 | 0 | 0 | 0 | 1 | 1 |
| PB0135 | 1 | 1 | 1 | 1 | 1 | 1 | 1 | 1 | 1 | 1 | 1 | 1 | 1 | 1 | 1 | 0 | 1 |
| PB0136 | 1 | 1 | 1 | 1 | 1 | 1 | 1 | 1 | 1 | 1 | 1 | 1 | 1 | 1 | 0 | 0 | 1 |
| PB0137 | 1 | 1 | 1 | 1 | 1 | 1 | 1 | 1 | 1 | 1 | 1 | 1 | 1 | 1 | 1 | 1 | 1 |
| PB0138 | 1 | 1 | 1 | 1 | 1 | 1 | 1 | 1 | 1 | 1 | 1 | 1 | 1 | 1 | 1 | 1 | 0 |
| PB0139 | 1 | 1 | 1 | 1 | 1 | 1 | 1 | 1 | 1 | 1 | 1 | 0 | 0 | 1 | 0 | 0 | 1 |
| PB0140 | 1 | 1 | 1 | 1 | 1 | 1 | 1 | 1 | 1 | 1 | 1 | 1 | 1 | 1 | 1 | 1 | 1 |
| PB0141 | 1 | 1 | 1 | 1 | 1 | 1 | 1 | 1 | 1 | 1 | 1 | 1 | 1 | 1 | 1 | 0 | 0 |
| PB0142 | 1 | 1 | 1 | 1 | 1 | 1 | 1 | 1 | 1 | 1 | 1 | 1 | 1 | 1 | 1 | 1 | 0 |
| PB0143 | 1 | 1 | 1 | 1 | 1 | 1 | 1 | 1 | 1 | 1 | 1 | 1 | 1 | 1 | 1 | 0 | 0 |
| PB0144 | 1 | 1 | 1 | 1 | 1 | 1 | 1 | 1 | 1 | 1 | 1 | 0 | 1 | 1 | 0 | 1 | 1 |
| PB0145 | 1 | 0 | 1 | 1 | 1 | 1 | 1 | 1 | 1 | 1 | 1 | 0 | 1 | 1 | 1 | 1 | 1 |

| | | | | | | | | | | | | | | | | | |
|---|---|---|---|---|---|---|---|---|---|---|---|---|---|---|---|---|---|
| PB0146 | 1 | 1 | 1 | 1 | 1 | 1 | 1 | 1 | 1 | 1 | 1 | 1 | 1 | 1 | 0 | 1 | 1 |
| PB0147 | 1 | 1 | 1 | 1 | 1 | 1 | 1 | 1 | 1 | 1 | 1 | 1 | 1 | 1 | 1 | 1 | 1 |
| PB0148 | 1 | 1 | 1 | 1 | 1 | 1 | 1 | 1 | 1 | 1 | 1 | 1 | 1 | 1 | 1 | 0 | 1 |
| PB0149 | 1 | 1 | 1 | 1 | 1 | 1 | 1 | 1 | 1 | 1 | 1 | 1 | 1 | 1 | 1 | 1 | 1 |
| PB0150 | 1 | 1 | 1 | 1 | 1 | 1 | 1 | 1 | 1 | 1 | 1 | 1 | 1 | 1 | 1 | 0 | 1 |
| PB0151 | 1 | 1 | 1 | 1 | 1 | 1 | 1 | 1 | 1 | 1 | 1 | 1 | 1 | 1 | 0 | 1 | 1 |
| PB0152 | 1 | 1 | 1 | 1 | 1 | 1 | 1 | 1 | 1 | 1 | 1 | 1 | 1 | 1 | 1 | 0 | 1 |
| PB0153 | 1 | 1 | 1 | 1 | 1 | 1 | 1 | 1 | 1 | 1 | 1 | 1 | 1 | 1 | 1 | 1 | 1 |
| PB0154 | 1 | 1 | 1 | 1 | 1 | 1 | 1 | 1 | 1 | 1 | 1 | 0 | 0 | 1 | 0 | 0 | 0 |
| PB0155 | 1 | 1 | 1 | 1 | 1 | 1 | 1 | 1 | 1 | 1 | 1 | 1 | 0 | 1 | 1 | 0 | 1 |
| PB0156 | 1 | 1 | 1 | 1 | 1 | 1 | 1 | 1 | 1 | 1 | 1 | 1 | 1 | 1 | 0 | 1 | 1 |
| PB0157 | 1 | 1 | 1 | 1 | 1 | 1 | 1 | 1 | 1 | 1 | 1 | 1 | 0 | 0 | 0 | 0 | 0 |
| PB0158 | 1 | 1 | 1 | 1 | 1 | 1 | 1 | 1 | 1 | 1 | 1 | 1 | 0 | 0 | 0 | 0 | 0 |
| PB0159 | 1 | 1 | 1 | 1 | 1 | 1 | 1 | 1 | 1 | 1 | 1 | 1 | 1 | 0 | 0 | 0 | 1 |
| PB0160 | 1 | 1 | 1 | 1 | 1 | 1 | 1 | 1 | 1 | 1 | 1 | 1 | 1 | 0 | 0 | 0 | 1 |
| PB0161 | 1 | 1 | 1 | 1 | 1 | 1 | 1 | 1 | 1 | 1 | 1 | 1 | 1 | 1 | 1 | 1 | 1 |
| PB0162 | 1 | 1 | 1 | 1 | 1 | 1 | 1 | 1 | 1 | 1 | 1 | 1 | 1 | 0 | 0 | 0 | 0 |
| PB0163 | 1 | 1 | 1 | 1 | 1 | 1 | 1 | 1 | 1 | 1 | 1 | 1 | 0 | 0 | 0 | 0 | 0 |
| PB0164 | 1 | 1 | 1 | 1 | 1 | 1 | 1 | 1 | 1 | 1 | 1 | 1 | 1 | 1 | 0 | 1 | 1 |
| PB0165 | 1 | 1 | 1 | 1 | 1 | 1 | 1 | 1 | 1 | 1 | 1 | 1 | 1 | 1 | 1 | 0 | 1 |
| PB0166 | 1 | 1 | 1 | 1 | 1 | 1 | 1 | 1 | 1 | 1 | 1 | 1 | 1 | 0 | 0 | 0 | 0 |
| PB0167 | 1 | 0 | 1 | 1 | 1 | 1 | 1 | 1 | 1 | 1 | 1 | 1 | 1 | 1 | 1 | 1 | 1 |
| PB0168 | 1 | 1 | 1 | 1 | 1 | 1 | 1 | 1 | 1 | 1 | 1 | 1 | 1 | 1 | 1 | 0 | 0 |
| PB0169 | 1 | 1 | 1 | 1 | 1 | 1 | 1 | 1 | 1 | 1 | 1 | 1 | 1 | 1 | 0 | 0 | 1 |
| PB0170 | 1 | 1 | 1 | 1 | 1 | 1 | 1 | 1 | 1 | 1 | 1 | 1 | 1 | 1 | 0 | 1 | 1 |
| PB0171 | 1 | 1 | 1 | 1 | 1 | 1 | 1 | 1 | 1 | 1 | 1 | 1 | 1 | 1 | 0 | 1 | 0 |
| PB0172 | 1 | 1 | 1 | 1 | 1 | 1 | 1 | 1 | 1 | 1 | 1 | 0 | 0 | 0 | 0 | 0 | 0 |
| PB0173 | 1 | 0 | 1 | 1 | 1 | 1 | 1 | 1 | 1 | 1 | 1 | 1 | 1 | 1 | 1 | 1 | 1 |
| PB0174 | 1 | 1 | 1 | 1 | 1 | 1 | 1 | 1 | 1 | 1 | 1 | 1 | 1 | 1 | 0 | 0 | 0 |
| PB0175 | 1 | 1 | 1 | 1 | 1 | 1 | 1 | 1 | 1 | 1 | 1 | 1 | 1 | 1 | 1 | 0 | 1 |
| PB0176 | 1 | 1 | 1 | 1 | 1 | 1 | 1 | 1 | 1 | 1 | 1 | 0 | 0 | 1 | 0 | 1 | 1 |
| PB0177 | 1 | 1 | 1 | 1 | 1 | 1 | 1 | 1 | 1 | 1 | 1 | 1 | 0 | 0 | 0 | 0 | 1 |
| PB0178 | 1 | 1 | 1 | 1 | 1 | 1 | 1 | 1 | 1 | 1 | 1 | 1 | 1 | 1 | 1 | 1 | 1 |
| PB0179 | 1 | 1 | 1 | 1 | 1 | 1 | 1 | 1 | 1 | 1 | 1 | 0 | 0 | 0 | 0 | 0 | 0 |
| PB0180 | 1 | 1 | 1 | 1 | 1 | 1 | 1 | 1 | 1 | 1 | 1 | 1 | 0 | 0 | 0 | 0 | 1 |
| PB0181 | 1 | 1 | 1 | 1 | 1 | 1 | 1 | 1 | 1 | 1 | 1 | 1 | 0 | 0 | 0 | 0 | 0 |
| PB0182 | 1 | 1 | 1 | 1 | 1 | 1 | 1 | 1 | 1 | 1 | 1 | 1 | 1 | 0 | 0 | 0 | 0 |
| PB0183 | 1 | 0 | 1 | 1 | 1 | 1 | 1 | 1 | 1 | 1 | 1 | 1 | 1 | 1 | 1 | 1 | 0 |
| PB0184 | 1 | 1 | 1 | 1 | 1 | 1 | 1 | 1 | 1 | 1 | 1 | 1 | 1 | 1 | 1 | 0 | 1 |
| PB0185 | 1 | 1 | 1 | 1 | 1 | 1 | 1 | 1 | 1 | 1 | 1 | 1 | 1 | 1 | 0 | 0 | 0 |
| PB0186 | 1 | 1 | 1 | 1 | 1 | 1 | 1 | 1 | 1 | 1 | 1 | 1 | 1 | 1 | 1 | 1 | 1 |
| PB0187 | 1 | 1 | 1 | 1 | 1 | 1 | 1 | 1 | 1 | 1 | 1 | 0 | 0 | 0 | 0 | 0 | 0 |
| PB0188 | 1 | 1 | 1 | 1 | 1 | 1 | 1 | 1 | 1 | 1 | 1 | 1 | 1 | 1 | 0 | 0 | 1 |
| PB0189 | 1 | 1 | 1 | 1 | 1 | 1 | 1 | 1 | 1 | 1 | 1 | 0 | 0 | 1 | 0 | 0 | 1 |
| PB0190 | 1 | 1 | 1 | 1 | 1 | 1 | 1 | 1 | 1 | 1 | 1 | 1 | 1 | 0 | 0 | 0 | 0 |
| PB0191 | 1 | 1 | 1 | 1 | 1 | 1 | 1 | 1 | 1 | 1 | 1 | 0 | 0 | 0 | 0 | 0 | 0 |
| PB0192 | 1 | 1 | 1 | 1 | 1 | 1 | 1 | 1 | 1 | 1 | 1 | 1 | 1 | 1 | 0 | 0 | 1 |
| PB0193 | 1 | 1 | 1 | 1 | 1 | 1 | 1 | 1 | 1 | 1 | 1 | 1 | 1 | 1 | 1 | 0 | 0 |
| PB0194 | 1 | 1 | 1 | 1 | 1 | 1 | 1 | 1 | 1 | 1 | 1 | 1 | 0 | 1 | 0 | 0 | 0 |
| PB0195 | 1 | 1 | 1 | 1 | 1 | 1 | 1 | 1 | 1 | 1 | 1 | 1 | 1 | 1 | 0 | 0 | 1 |
| PB0196 | 1 | 1 | 1 | 1 | 1 | 1 | 1 | 1 | 1 | 1 | 1 | 1 | 0 | 1 | 0 | 0 | 0 |
| PB0197 | 1 | 1 | 1 | 1 | 1 | 1 | 1 | 1 | 1 | 1 | 1 | 1 | 1 | 1 | 0 | 0 | 1 |
| PB0198 | 1 | 1 | 1 | 1 | 1 | 1 | 1 | 1 | 1 | 1 | 1 | 1 | 1 | 1 | 1 | 0 | 1 |
| PB0199 | 1 | 0 | 1 | 0 | 1 | 1 | 0 | 1 | 1 | 1 | 1 | 1 | 1 | 1 | 1 | 0 | 1 |
| PB0200 | 1 | 1 | 1 | 1 | 1 | 1 | 1 | 1 | 1 | 1 | 1 | 0 | 0 | 0 | 0 | 0 | 0 |
| PB0201 | 1 | 1 | 1 | 1 | 1 | 1 | 1 | 1 | 1 | 1 | 1 | 1 | 1 | 1 | 1 | 0 | 1 |
| PB0202 | 1 | 1 | 1 | 1 | 1 | 1 | 1 | 1 | 1 | 1 | 1 | 1 | 1 | 0 | 0 | 0 | 1 |
| PB0203 | 1 | 1 | 1 | 1 | 1 | 1 | 1 | 0 | 1 | 1 | 1 | 1 | 1 | 1 | 1 | 0 | 1 |

| | | | | | | | | | | | | | | | | | |
|---|---|---|---|---|---|---|---|---|---|---|---|---|---|---|---|---|---|
| PB0204 | 1 | 1 | 1 | 1 | 1 | 1 | 1 | 1 | 1 | 1 | 1 | 1 | 1 | 1 | 0 | 1 | 1 |
| PB0205 | 1 | 1 | 1 | 1 | 1 | 1 | 1 | 1 | 1 | 1 | 1 | 1 | 1 | 1 | 1 | 0 | 1 |
| PB0206 | 1 | 1 | 1 | 1 | 1 | 1 | 1 | 1 | 1 | 1 | 1 | 0 | 0 | 0 | 0 | 0 | 0 |
| PB0207 | 1 | 1 | 1 | 1 | 1 | 1 | 1 | 1 | 1 | 1 | 1 | 0 | 1 | 0 | 1 | 1 | 1 |
| PB0208 | 1 | 1 | 1 | 1 | 1 | 1 | 1 | 1 | 1 | 1 | 1 | 1 | 1 | 1 | 1 | 1 | 1 |
| PB0209 | 1 | 1 | 1 | 1 | 1 | 1 | 1 | 1 | 1 | 1 | 1 | 1 | 1 | 1 | 1 | 1 | 1 |
| PB0210 | 1 | 1 | 1 | 1 | 1 | 1 | 1 | 1 | 1 | 0 | 0 | 0 | 0 | 0 | 0 | 0 | 0 |
| PB0211 | 1 | 1 | 1 | 1 | 1 | 1 | 1 | 1 | 1 | 1 | 1 | 1 | 1 | 1 | 1 | 1 | 1 |
| PB0212 | 1 | 1 | 1 | 1 | 1 | 1 | 1 | 1 | 1 | 1 | 1 | 0 | 0 | 0 | 0 | 0 | 0 |
| PB0213 | 1 | 1 | 1 | 1 | 1 | 1 | 1 | 1 | 1 | 1 | 1 | 1 | 1 | 0 | 1 | 1 | 1 |
| PB0214 | 1 | 1 | 1 | 1 | 1 | 1 | 1 | 1 | 1 | 1 | 1 | 1 | 1 | 1 | 1 | 1 | 1 |
| PB0215 | 1 | 1 | 1 | 1 | 1 | 1 | 1 | 1 | 1 | 1 | 1 | 1 | 1 | 1 | 1 | 0 | 1 |
| PB0216 | 1 | 1 | 1 | 1 | 1 | 1 | 1 | 1 | 1 | 1 | 1 | 0 | 1 | 1 | 0 | 1 | 0 |
| PB0217 | 1 | 0 | 1 | 1 | 1 | 1 | 1 | 1 | 1 | 1 | 1 | 0 | 0 | 0 | 0 | 0 | 0 |
| PB0218 | 1 | 1 | 1 | 1 | 1 | 1 | 1 | 1 | 1 | 1 | 1 | 0 | 0 | 0 | 0 | 0 | 0 |
| PB0219 | 1 | 1 | 1 | 1 | 1 | 1 | 1 | 1 | 1 | 1 | 1 | 1 | 0 | 0 | 0 | 0 | 0 |
| PB0220 | 1 | 1 | 1 | 1 | 1 | 1 | 1 | 1 | 1 | 1 | 1 | 1 | 1 | 0 | 0 | 0 | 1 |
| PB0221 | 1 | 1 | 1 | 1 | 1 | 1 | 1 | 1 | 1 | 1 | 1 | 0 | 1 | 1 | 0 | 1 | 0 |
| PB0222 | 1 | 1 | 1 | 1 | 1 | 1 | 1 | 1 | 1 | 1 | 1 | 0 | 0 | 1 | 0 | 0 | 0 |
| PB0223 | 1 | 1 | 1 | 1 | 1 | 1 | 1 | 1 | 1 | 1 | 1 | 1 | 1 | 1 | 1 | 0 | 1 |
| PB0224 | 1 | 1 | 1 | 1 | 1 | 1 | 1 | 1 | 1 | 1 | 1 | 1 | 0 | 1 | 0 | 0 | 0 |
| PB0225 | 1 | 1 | 1 | 1 | 1 | 1 | 1 | 1 | 1 | 1 | 1 | 1 | 1 | 1 | 0 | 1 | 1 |
| PB0226 | 1 | 1 | 1 | 1 | 1 | 1 | 1 | 1 | 1 | 1 | 1 | 1 | 1 | 1 | 0 | 1 | 1 |
| PB0227 | 1 | 1 | 1 | 1 | 1 | 1 | 1 | 1 | 1 | 1 | 1 | 1 | 1 | 1 | 0 | 1 | 0 |
| PB0228 | 1 | 1 | 1 | 1 | 1 | 1 | 1 | 1 | 1 | 1 | 1 | 0 | 0 | 0 | 0 | 0 | 0 |
| PB0229 | 1 | 0 | 1 | 1 | 1 | 1 | 1 | 1 | 1 | 1 | 1 | 0 | 0 | 1 | 0 | 0 | 0 |
| PB0230 | 1 | 1 | 1 | 1 | 1 | 1 | 1 | 1 | 1 | 1 | 1 | 0 | 0 | 1 | 0 | 0 | 1 |
| PB0231 | 1 | 1 | 1 | 1 | 1 | 1 | 1 | 1 | 1 | 1 | 1 | 1 | 1 | 1 | 1 | 1 | 1 |
| PB0232 | 1 | 1 | 1 | 1 | 1 | 1 | 1 | 1 | 1 | 1 | 1 | 1 | 0 | 0 | 0 | 0 | 1 |
| PB0233 | 1 | 1 | 1 | 1 | 1 | 1 | 1 | 1 | 1 | 1 | 1 | 1 | 0 | 1 | 0 | 0 | 1 |
| PB0234 | 1 | 1 | 1 | 1 | 1 | 1 | 1 | 1 | 1 | 1 | 1 | 1 | 1 | 1 | 1 | 1 | 1 |
| PB0235 | 1 | 1 | 1 | 1 | 1 | 1 | 1 | 1 | 1 | 1 | 1 | 1 | 1 | 1 | 1 | 1 | 1 |
| PB0236 | 1 | 1 | 1 | 1 | 1 | 1 | 1 | 1 | 1 | 1 | 1 | 1 | 1 | 1 | 1 | 1 | 1 |
| PB0237 | 1 | 1 | 1 | 1 | 1 | 1 | 1 | 1 | 1 | 1 | 1 | 1 | 1 | 1 | 1 | 1 | 1 |
| PB0238 | 1 | 1 | 1 | 1 | 1 | 1 | 1 | 1 | 1 | 1 | 1 | 1 | 0 | 0 | 0 | 0 | 0 |
| PB0239 | 1 | 1 | 1 | 1 | 1 | 1 | 1 | 1 | 1 | 1 | 1 | 1 | 1 | 1 | 1 | 1 | 1 |
| PB0240 | 1 | 1 | 1 | 1 | 1 | 1 | 1 | 1 | 1 | 1 | 1 | 1 | 1 | 1 | 1 | 0 | 1 |
| PB0241 | 1 | 1 | 1 | 1 | 1 | 1 | 1 | 1 | 1 | 1 | 1 | 1 | 1 | 0 | 0 | 1 | 1 |
| PB0242 | 1 | 1 | 1 | 1 | 1 | 1 | 1 | 1 | 1 | 1 | 1 | 0 | 0 | 0 | 0 | 0 | 0 |
| PB0243 | 1 | 1 | 1 | 1 | 1 | 1 | 1 | 1 | 1 | 1 | 1 | 0 | 1 | 0 | 0 | 0 | 0 |
| PB0244 | 1 | 1 | 1 | 1 | 1 | 1 | 1 | 1 | 1 | 1 | 1 | 0 | 1 | 1 | 1 | 0 | 0 |
| PB0245 | 1 | 1 | 1 | 1 | 1 | 1 | 1 | 1 | 1 | 1 | 1 | 1 | 1 | 0 | 1 | 0 | 1 |
| PB0246 | 1 | 1 | 1 | 1 | 1 | 1 | 1 | 1 | 1 | 1 | 1 | 1 | 1 | 1 | 1 | 1 | 1 |
| PB0247 | 1 | 1 | 1 | 1 | 1 | 1 | 1 | 1 | 1 | 1 | 1 | 1 | 1 | 1 | 0 | 0 | 1 |
| PB0248 | 1 | 1 | 1 | 1 | 1 | 1 | 1 | 1 | 1 | 1 | 1 | 1 | 1 | 1 | 1 | 1 | 1 |
| PB0249 | 1 | 1 | 1 | 1 | 1 | 1 | 1 | 1 | 1 | 0 | 0 | 0 | 0 | 0 | 0 | 0 | 0 |
| PB0250 | 1 | 1 | 1 | 1 | 1 | 1 | 1 | 1 | 1 | 1 | 0 | 0 | 0 | 0 | 0 | 0 | 0 |
| PB0251 | 1 | 1 | 1 | 1 | 1 | 1 | 1 | 1 | 1 | 1 | 1 | 1 | 1 | 1 | 0 | 0 | 1 |
| PB0252 | 1 | 1 | 1 | 1 | 1 | 1 | 1 | 1 | 1 | 1 | 1 | 1 | 1 | 0 | 0 | 1 | 0 |
| PB0253 | 1 | 1 | 1 | 1 | 1 | 1 | 1 | 1 | 1 | 1 | 1 | 1 | 1 | 0 | 0 | 1 | 0 |
| PB0254 | 1 | 1 | 1 | 1 | 1 | 1 | 1 | 1 | 1 | 1 | 1 | 1 | 1 | 0 | 0 | 1 | 0 |
| PB0255 | 1 | 1 | 1 | 1 | 1 | 1 | 1 | 1 | 1 | 1 | 1 | 1 | 1 | 0 | 0 | 1 | 0 |

*N: Number of species; Pm: Primate; Ra: Rabbit; Ro: Rodent; Ca: Carnivore; De: Deer; Bo: Bovid; Pi: Pig; Re: Reptile; Bi: Bird; Am: amphibian; Fi: Bony fishes; Tu: Tunicates; Ec: Echinoderm; In: Insects; Ne: Nematode; Ce: Cephalopod; and Cn: Cnidarian*

**Supplementary table S4. Domain present in human bone extracellular matrix proteins dataset.** Data of domains in other species is available from the database.

| Domain | N | Type | Description |
|---|---|---|---|
| PF01391_Collagen | 30 | Repeat | Collagen triple helix repeat |
| PF13855_LRR_8 | 16 | Repeat | Leucine rich repeat |
| PF00089_Trypsin | 15 | Domain | Trypsin |
| PF00092_VWA | 15 | Domain | von Willebrand factor type A domain |
| PF00008_EGF | 14 | Domain | EGF-like domain |
| PF07645_EGF_CA | 13 | Domain | Calcium-binding EGF domain |
| PF00053_Laminin_EGF | 10 | Domain | Laminin EGF domain |
| PF00093_VWC | 9 | Domain | von Willebrand factor type C domain |
| PF12662_cEGF | 9 | Domain | Complement Clr-like EGF-like |
| PF00594_Gla | 8 | Domain | Vitamin K-dependent carboxylation/gamma-carboxyglutamic (GLA) domain |
| PF01410_COLFI | 8 | Family | Fibrillar collagen C-terminal domain |
| PF01462_LRRNT | 8 | Family | Leucine rich repeat N-terminal domain |
| PF02210_Laminin_G_2 | 8 | Domain | Laminin G domain |
| PF00041_fn3 | 7 | Domain | Fibronectin type III domain |
| PF00079_Serpin | 7 | Domain | Serpin (serine protease inhibitor) |
| PF00147_Fibrinogen_C | 7 | Domain | Fibrinogen beta and gamma chains, C-terminal globular domain |
| PF01759_NTR | 7 | Domain | UNC-6/NTR/C345C module |
| PF00054_Laminin_G_1 | 6 | Domain | Laminin G domain |
| PF00055_Laminin_N | 6 | Domain | Laminin N-terminal (Domain VI) |
| PF00084_Sushi | 6 | Domain | Sushi repeat (SCR repeat) |
| PF00090_TSP_1 | 6 | Domain | Thrombospondin type 1 domain |
| PF00219_IGFBP | 6 | Domain | Insulin-like growth factor binding protein |
| PF00019_TGF_beta | 5 | Domain | Transforming growth factor beta like domain |
| PF00094_VWD | 5 | Domain | von Willebrand factor type D domain |
| PF01392_Fz | 5 | Domain | Fz domain |
| PF01413_C4 | 5 | Domain | C-terminal tandem repeated domain in type 4 procollagen |
| PF07648_Kazal_2 | 5 | Domain | Kazal-type serine protease inhibitor domain |
| PF00045_Hemopexin | 4 | Repeat | Hemopexin |
| PF00052_Laminin_B | 4 | Domain | Laminin B (Domain IV) |
| PF00059_Lectin_C | 4 | Domain | Lectin C-type domain |
| PF00086_Thyroglobulin_1 | 4 | Domain | Thyroglobulin type-1 repeat |
| PF00193_Xlink | 4 | Domain | Extracellular link domain |
| PF00688_TGFb_propeptide | 4 | Family | TGF-beta propeptide |
| PF01826_TIL | 4 | Domain | Trypsin Inhibitor like cysteine rich domain |
| PF02412_TSP_3 | 4 | Domain | Thrombospondin type 3 repeat |
| PF05735_TSP_C | 4 | Family | Thrombospondin C-terminal region |
| PF08742_C8 | 4 | Domain | C8 domain |
| PF15913_Furin-like_2 | 4 | Domain | Furin-like repeat, cysteine-rich |
| PF00014_Kunitz_BPTI | 3 | Domain | Kunitz/Bovine pancreatic trypsin inhibitor domain |
| PF00031_Cystatin | 3 | Domain | Cystatin domain |
| PF00040_fn2 | 3 | Domain | Fibronectin type II domain |
| PF00051_Kringle | 3 | Domain | Kringle domain |
| PF00191_Annexin | 3 | Domain | Annexind |
| PF00207_A2M | 3 | Family | Alpha-2-macroglobulin family |
| PF00307_CH | 3 | Domain | Calponin homology (CH) domain |
| PF00386_C1q | 3 | Domain | C1q domain |
| PF00413_Peptidase_M10 | 3 | Domain | Matrixin |
| PF00431_CUB | 3 | Domain | CUB domain |
| PF00683_TB | 3 | Family | TB domain |
| PF01442_Apolipoprotein | 3 | Domain | Apolipoprotein A1/A4/E domain |
| PF01471_PG_binding_1 | 3 | Domain | Putative peptidoglycan binding domain |
| PF01835_MG2 | 3 | Domain | MG2 domain |
| PF02191_OLF | 3 | Repeat | Olfactomedin-like domain |

| | | | |
|---|---|---|---|
| PF06008_Laminin_I | 3 | Coiled-coil | Laminin Domain I |
| PF06009_Laminin_II | 3 | Coiled-coil | Laminin Domain II |
| PF07677_A2M_recep | 3 | Domain | A-macroglobulin receptor binding domain |
| PF07678_TED_complement | 3 | Repeat | A-macroglobulin TED domain |
| PF07686_V-set | 3 | Domain | Immunoglobulin V-set domain |
| PF07703_A2M_BRD | 3 | Domain | Alpha-2-macroglobulin bait region domain |
| PF07974_EGF_2 | 3 | Domain | EGF-like domain |
| PF08702_Fib_alpha | 3 | Coiled-coil | Fibrinogen alpha/beta chain family |
| PF09289_FOLN | 3 | Domain | Follistatin/Osteonectin-like EGF domain |
| PF10393_Matrilin_ccoil | 3 | Coiled-coil | Trimeric coiled-coil oligomerisation domain of matrilin |
| PF12661_hEGF | 3 | Domain | Human growth factor-like EGF |
| PF12947_EGF_3 | 3 | Domain | EGF domain |
| PF13330_Mucin2_WxxW | 3 | Family | Mucin-2 protein WxxW repeating region |
| PF13833_EF-hand_8 | 3 | Domain | EF-hand domain pair |
| PF17789_MG4 | 3 | Domain | Macroglobulin domain MG4 |
| PF17791_MG3 | 3 | Domain | Macroglobulin domain MG3 |
| PF18720_EGF_Tenascin | 3 | Domain | Tenascin EGF domain |
| PF06119_NIDO | 3 | Family | Nidogen-like |
| PF00050_Kazal_1 | 2 | Domain | Kazal-type serine protease inhibitor domain |
| PF00057_Ldl_recept_a | 2 | Repeat | Low-density lipoprotein receptor domain class A |
| PF00058_Ldl_recept_b | 2 | Repeat | Low-density lipoprotein receptor repeat class B |
| PF00061_Lipocalin | 2 | Domain | Lipocalin / cytosolic fatty-acid binding protein family |
| PF00106_adh_short | 2 | Domain | Short chain dehydrogenase |
| PF00405_Transferrin | 2 | Domain | Transferrin |
| PF00530_SRCR | 2 | Domain | Scavenger receptor cysteine-rich domain |
| PF00560_LRR_1 | 2 | Repeat | Leucine Rich Repeat |
| PF00626_Gelsolin | 2 | Domain | Gelsolin repeat |
| PF00648_Peptidase_C2 | 2 | Family | Calpain family cysteine protease |
| PF01033_Somatomedin_B | 2 | Family | Somatomedin B domain |
| PF01067_Calpain_III | 2 | Domain | Calpain large subunit, domain III |
| PF01186_Lysyl_oxidase | 2 | Family | Lysyl oxidase |
| PF01463_LRRCT | 2 | Family | Leucine rich repeat C-terminal domain |
| PF01821_ANATO | 2 | Domain | Anaphylotoxin-like domain |
| PF01839_FG-GAP | 2 | Repeat | FG-GAP repeat |
| PF02469_Fasciclin | 2 | Domain | Fasciclin domain |
| PF04089_BRICHOS | 2 | Domain | BRICHOS domain |
| PF07474_G2F | 2 | Domain | G2F domain |
| PF08441_Integrin_alpha2 | 2 | Family | Integrin alpha |
| PF11598_COMP | 2 | Family | Cartilage oligomeric matrix protein |
| PF12714_TILa | 2 | Domain | TILa domain |
| PF13499_EF-hand_7 | 2 | Domain | EF-hand domain pair |
| PF13516_LRR_6 | 2 | Repeat | Leucine Rich repeat |
| PF13778_DUF4174 | 2 | Family | Domain of unknown function (DUF4174) |
| PF17790_MG1 | 2 | Domain | Macroglobulin domain MG1 |
| PF17820_PDZ_6 | 2 | Domain | PDZ domain |
| PF18193_Fibrillin_U_N | 2 | Domain | Fibrillin 1 unique N-terminal domain |
| PF00754_F5_F8_type_C | 2 | Domain | F5/8 type C domain |
| PF00246_Peptidase_M14 | 2 | Domain | Zinc carboxypeptidase |
| PF19028_TSP1_spondin | 2 | Domain | Spondin-like TSP1 domain |
| PF00007_Cys_knot | 1 | Domain | Cystine-knot domain |
| PF00012_HSP70 | 1 | Family | Hsp70 protein |
| PF00018_SH3_1 | 1 | Domain | SH3 domain |
| PF00024_PAN_1 | 1 | Domain | PAN domain |
| PF00039_fn1 | 1 | Domain | Fibronectin type I domain |
| PF00042_Globin | 1 | Domain | Globin |
| PF00044_Gp_dh_N | 1 | Domain | Glyceraldehyde 3-phosphate dehydrogenase, NAD binding domain |
| PF00048_IL8 | 1 | Domain | Small cytokines (intecrine/chemokine), interleukin-8 like |
| PF00062_Lys | 1 | Domain | C-type lysozyme/alpha-lactalbumin family |

| | | | |
|---|---|---|---|
| PF00071_Ras | 1 | Domain | Ras family |
| PF00080_Sod_Cu | 1 | Domain | Copper/zinc superoxide dismutase (SODC) |
| PF00085_Thioredoxin | 1 | Domain | Thioredoxin |
| PF00100_Zona_pellucida | 1 | Family | Zona pellucida-like domain |
| PF00112_Peptidase_C1 | 1 | Domain | Papain family cysteine protease |
| PF00167_FGF | 1 | Domain | Fibroblast growth factor |
| PF00169_PH | 1 | Domain | PH domain |
| PF00210_Ferritin | 1 | Domain | Ferritin-like domain |
| PF00245_Alk_phosphatase | 1 | Domain | Alkaline phosphatase |
| PF00354_Pentaxin | 1 | Domain | Pentaxin family |
| PF00357_Integrin_alpha | 1 | Family | Integrin alpha cytoplasmic region |
| PF00362_Integrin_beta | 1 | Domain | Integrin beta chain VWA domain |
| PF00373_FERM_M | 1 | Domain | FERM central domain |
| PF00412_LIM | 1 | Domain | LIM domain |
| PF00435_Spectrin | 1 | Domain | Spectrin repeat |
| PF00445_Ribonuclease_T2 | 1 | Domain | Ribonuclease T2 family |
| PF00505_HMG_box | 1 | Domain | HMG (high mobility group) box |
| PF00514_Arm | 1 | Repeat | Armadillo |
| PF00576_Transthyretin | 1 | Domain | HIUase/Transthyretin family |
| PF00578_AhpC-TSA | 1 | Domain | AhpC/TSA family |
| PF00594_GLA | 1 | Domain | Vitamin K-dependent carboxylation/gamma-carboxyglutamic (GLA) domain |
| PF00630_Filamin | 1 | Domain | Filamin/ABP280 repeat |
| PF00787_PX | 1 | Domain | PX domain |
| PF00865_Osteopontin | 1 | Family | Osteopontin |
| PF00965_TIMP | 1 | Domain | Tissue inhibitor of metalloproteinase |
| PF01023_S_100 | 1 | Domain | S-100/ICaBP type calcium binding domain |
| PF01044_Vinculin | 1 | Family | Vinculin family |
| PF01091_PTN_MK_C | 1 | Domain | PTN/MK heparin-binding protein family, C-terminal domain |
| PF01093_Clusterin | 1 | Family | Clusterin |
| PF01122_Cobalamin_bind | 1 | Repeat | Eukaryotic cobalamin-binding protein |
| PF01223_Endonuclease_NS | 1 | Domain | DNA/RNA non-specific endonuclease |
| PF01347_Vitellogenin_N | 1 | Repeat | Lipoprotein amino terminal region |
| PF01390_SEA | 1 | Family | SEA domain |
| PF01400_Astacin | 1 | Domain | Astacin (Peptidase family M12A) |
| PF01549_ShK | 1 | Domain | ShK domain-like |
| PF01663_Phosphodiest | 1 | Family | Type I phosphodiesterase / nucleotide pyrophosphatase |
| PF01731_Arylesterase | 1 | Repeat | Arylesterase |
| PF02014_Reeler | 1 | Family | Reeler domain |
| PF02177_APP_N | 1 | Domain | Amyloid A4 N-terminal heparin-binding |
| PF02494_HYR | 1 | Domain | HYR domain |
| PF02800_Gp_dh_C | 1 | Domain | lyceraldehyde 3-phosphate dehydrogenase, C-terminal domain |
| PF03045_DAN | 1 | Domain | DAN domain |
| PF03146_NtA | 1 | Domain | Agrin NtA domain |
| PF03494_Beta-APP | 1 | Family | Beta-amyloid peptide (beta-APP) |
| PF03815_LCCL | 1 | Domain | LCCL domain |
| PF04587_ADP_PFK_GK | 1 | Family | ADP-specific Phosphofructokinase/Glucokinase conserved region |
| PF04691_ApoC-I | 1 | Family | Apolipoprotein C-I (ApoC-1) |
| PF04706_Dickkopf_N | 1 | Family | Dickkopf N-terminal cysteine-rich region |
| PF04711_ApoA-II | 1 | Family | Apolipoprotein A-II (ApoA-II) |
| PF04777_Evr1_Alr | 1 | Family | Erv1 / Alr family |
| PF05196_PTN_MK_N | 1 | Domain | PTN/MK heparin-binding protein family, N-terminal domain |
| PF05355_Apo-CII | 1 | Family | Apolipoprotein C-II |
| PF05432_BSP_II | 1 | Family | Bone sialoprotein II (BSP-II) |
| PF05507_MAGP | 1 | Family | Microfibril-associated glycoprotein (MAGP) |
| PF06121_DUF959 | 1 | Domain | Domain of Unknown Function (DUF959) |
| PF06448_DUF1081 | 1 | Family | Domain of Unknown Function (DUF1081) |
| PF06468_Spond_N | 1 | Domain | Spondin_N |
| PF06473_FGF-BP1 | 1 | Family | FGF binding protein 1 (FGF-BP1) |

| | | | |
|---|---|---|---|
| PF06482_Endostatin | 1 | Domain | Collagenase NC10 and Endostatin |
| PF06668_ITI_HC_C | 1 | Family | Inter-alpha-trypsin inhibitor heavy chain C-terminus |
| PF06951_PLA2G12 | 1 | Domain | Group XII secretory phospholipase A2 precursor (PLA2G12) |
| PF07448_Spp-24 | 1 | Domain | Secreted phosphoprotein 24 (Spp-24) cystatin-like domain |
| PF07546_EMI | 1 | Domain | EMI domain |
| PF07653_SH3_2 | 1 | Domain | Variant SH3 domain |
| PF07654_C1-set | 1 | Domain | Immunoglobulin C1-set domain |
| PF07679_I-set | 1 | Domain | Immunoglobulin I-set domain |
| PF07965_Integrin_B_tail | 1 | Domain | Integrin beta tail domain |
| PF08246_Inhibitor_I29 | 1 | Domain | Cathepsin propeptide inhibitor domain (I29) |
| PF08391_Ly49 | 1 | Family | Ly49-like protein, N-terminal region |
| PF08487_VIT | 1 | Family | Vault protein inter-alpha-trypsin domain |
| PF08725_Integrin_b_cyt | 1 | Domain | Integrin beta cytoplasmic domain |
| PF08726_EFhand_Ca_insen | 1 | Domain | Ca2+ insensitive EF hand |
| PF09011_HMG_box_2 | 1 | Domain | HMG-box domain |
| PF09172_Vit_open_b-sht | 1 | Domain | Vitellinogen, open beta-sheet |
| PF09396_Thrombin_light | 1 | Domain | Thrombin light chain |
| PF10179_NDNF | 1 | Domain | Neuron-derived neurotrophic factor, first Fn(III) domain |
| PF10417_1-cysPrx_C | 1 | Domain | C-terminal domain of 1-Cys peroxiredoxin |
| PF10515_APP_amyloid | 1 | Family | Beta-amyloid precursor protein C-terminus |
| PF10591_SPARC_Ca_bdg | 1 | Domain | Secreted protein acidic and rich in cysteine Ca binding region |
| PF11032_ApoM | 1 | Domain | ApoM domain |
| PF12160_Fibrinogen_aC | 1 | Domain | Fibrinogen alpha C domain |
| PF12491_ApoB100_C | 1 | Family | Apolipoprotein B100 C terminal |
| PF12924_APP_Cu_bd | 1 | Domain | Copper-binding of amyloid precursor, CuBD |
| PF12925_APP_E2 | 1 | Domain | E2 domain of amyloid precursor protein |
| PF13202_EF-hand_5 | 1 | Domain | EF hand |
| PF13306_LRR_5 | 1 | Repeat | BspA type Leucine rich repeat region (6 copies) |
| PF13895_Ig_2 | 1 | Domain | Immunoglobulin domain |
| PF13927_IG_3 | 1 | Domain | Immunoglobulin domain |
| PF14478_DUF4430 | 1 | Domain | Domain of unknown function (DUF4430) |
| PF14704_DERM | 1 | Family | Dermatopontin |
| PF15182_OTOS | 1 | Family | Otospiralin |
| PF15467_SGIII | 1 | Family | Secretogranin-3 |
| PF15511_CENP-T_C | 1 | Domain | Centromere kinetochore component CENP-T histone fold |
| PF15819_Fibin | 1 | Family | Fin bud initiation factor homologue |
| PF17085_UCMA | 1 | Family | Unique cartilage matrix associated protein |
| PF17205_PSI_integrin | 1 | Domain | Integrin plexin domain |
| PF17517_IgGFc_binding | 1 | Family | IgGFc binding protein |
| PF18108_QSOX_Trx1 | 1 | Domain | QSOX Trx-like domain |
| PF18124_Kindlin_2_N | 1 | Domain | Kindlin-2 N-terminal domain |
| PF18371_FAD_SOX | 1 | Domain | Flavin adenine dinucleotide (FAD)-dependent sulfhydryl oxidase |
| PF18372_I-EGF_1 | 1 | Domain | Integrin beta epidermal growth factor like domain 1 |
| PF19035_TSP1_CCN | 1 | Domain | CCN3 Nov like TSP1 domain |
| PF20010_Collagen_trimer | 1 | Domain | Collagen trimerization domain |

*N = Number of proteins.*

**Supplementary table 5.** List of proteins that are target of existing drugs in the market.

| Drugs \ Uniprot codes | O14793 | P00734 | P00740 | P00742 | P00747 | P00751 | P01009 | P01024 | P01031 | P01042 | P02743 | P02753 | P02776 | P04114 | P05121 | P05186 | P05556 | P06756 | P07477 | P08253 | P08648 | P10600 | P10909 | P14780 | P21333 | P29279 | P43235 | P45452 | P61812 | P63000 | P69905 | Q92743 | Q9BX93 | Q9BXY4 | Q9GZV9 | Q9UNI1 |
|---|---|---|---|---|---|---|---|---|---|---|---|---|---|---|---|---|---|---|---|---|---|---|---|---|---|---|---|---|---|---|---|---|---|---|---|---|
| D00160 - Aminocaproic acid (USP/INN); Epsilon-Aminocaproic acid (JAN); Amicar (TN) | 0 | 0 | 0 | 0 | 1 | 0 | 0 | 0 | 0 | 0 | 0 | 0 | 0 | 0 | 0 | 0 | 0 | 0 | 0 | 0 | 0 | 0 | 0 | 0 | 0 | 0 | 0 | 0 | 0 | 0 | 0 | 0 | 0 | 0 | 0 | 0 |
| D00181 - Argatroban (USP); Argatroban hydrate (JP18); Novastan (TN) | 0 | 1 | 0 | 0 | 0 | 0 | 0 | 0 | 0 | 0 | 0 | 0 | 0 | 0 | 0 | 0 | 0 | 0 | 0 | 0 | 0 | 0 | 0 | 0 | 0 | 0 | 0 | 0 | 0 | 0 | 0 | 0 | 0 | 0 | 0 | 0 |
| D01136 - Tranexamic acid (JP18/USP/INN); Cyklokapron (TN); Rikavarin (TN); Transamin (TN) | 0 | 0 | 0 | 0 | 1 | 0 | 0 | 0 | 0 | 0 | 0 | 0 | 0 | 0 | 0 | 0 | 0 | 0 | 0 | 0 | 0 | 0 | 0 | 0 | 0 | 0 | 0 | 0 | 0 | 0 | 0 | 0 | 0 | 0 | 0 | 0 |
| D01844 - Fondaparinux sodium (JAN/USP/INN); Arixtra (TN) | 0 | 0 | 0 | 1 | 0 | 0 | 0 | 0 | 0 | 0 | 0 | 0 | 0 | 0 | 0 | 0 | 0 | 0 | 0 | 0 | 0 | 0 | 0 | 0 | 0 | 0 | 0 | 0 | 0 | 0 | 0 | 0 | 0 | 0 | 0 | 0 |
| D01981 - Ximelagatran (JAN/USAN/INN); Exanta (TN) | 0 | 1 | 0 | 0 | 0 | 0 | 0 | 0 | 0 | 0 | 0 | 0 | 0 | 0 | 0 | 0 | 0 | 0 | 0 | 0 | 0 | 0 | 0 | 0 | 0 | 0 | 0 | 0 | 0 | 0 | 0 | 0 | 0 | 0 | 0 | 0 |
| D03061 - Batimastat (USAN/INN) | 0 | 0 | 0 | 0 | 0 | 0 | 0 | 0 | 0 | 0 | 0 | 0 | 0 | 0 | 0 | 0 | 0 | 0 | 0 | 1 | 0 | 0 | 0 | 1 | 0 | 0 | 0 | 0 | 0 | 0 | 0 | 0 | 0 | 0 | 0 | 0 |
| D03136 - Bivalirudin (USAN/INN) | 0 | 1 | 0 | 0 | 0 | 0 | 0 | 0 | 0 | 0 | 0 | 0 | 0 | 0 | 0 | 0 | 0 | 0 | 0 | 0 | 0 | 0 | 0 | 0 | 0 | 0 | 0 | 0 | 0 | 0 | 0 | 0 | 0 | 0 | 0 | 0 |
| D03213 - Apixaban (JAN/USAN/INN); Eliquis (TN) | 0 | 0 | 0 | 1 | 0 | 0 | 0 | 0 | 0 | 0 | 0 | 0 | 0 | 0 | 0 | 0 | 0 | 0 | 0 | 0 | 0 | 0 | 0 | 0 | 0 | 0 | 0 | 0 | 0 | 0 | 0 | 0 | 0 | 0 | 0 | 0 |
| D03330 - Kallidinogenase (JP18/INN); Cacelmin A (TN) | 0 | 0 | 0 | 0 | 0 | 0 | 0 | 0 | 0 | 1 | 0 | 0 | 0 | 0 | 0 | 0 | 0 | 0 | 0 | 0 | 0 | 0 | 0 | 0 | 0 | 0 | 0 | 0 | 0 | 0 | 0 | 0 | 0 | 0 | 0 | 0 |
| D03436 - Cellulose; oxidized (JAN/USP); Surgicel (TN) | 0 | 0 | 0 | 0 | 0 | 0 | 0 | 0 | 0 | 0 | 0 | 0 | 0 | 0 | 0 | 0 | 0 | 0 | 0 | 0 | 0 | 0 | 0 | 0 | 0 | 0 | 0 | 0 | 0 | 0 | 1 | 0 | 0 | 0 | 0 | 0 |
| D03497 - Cilengitide (USAN/INN) | 0 | 0 | 0 | 0 | 0 | 0 | 0 | 0 | 0 | 0 | 0 | 0 | 0 | 0 | 0 | 0 | 0 | 1 | 0 | 0 | 0 | 0 | 0 | 0 | 0 | 0 | 0 | 0 | 0 | 0 | 0 | 0 | 0 | 0 | 0 | 0 |
| D03517 - Cipemastat (USAN/INN); Trocade (TN) | 0 | 0 | 0 | 0 | 0 | 0 | 0 | 0 | 0 | 0 | 0 | 0 | 0 | 0 | 0 | 0 | 0 | 0 | 0 | 0 | 0 | 0 | 0 | 1 | 0 | 0 | 0 | 1 | 0 | 0 | 0 | 0 | 0 | 0 | 0 | 0 |
| D03692 - Desirudin (USAN/INN); Iprivask (TN) | 0 | 1 | 0 | 0 | 0 | 0 | 0 | 0 | 0 | 0 | 0 | 0 | 0 | 0 | 0 | 0 | 0 | 0 | 0 | 0 | 0 | 0 | 0 | 0 | 0 | 0 | 0 | 0 | 0 | 0 | 0 | 0 | 0 | 0 | 0 | 0 |
| D03722 - Efegatran sulfate (USAN) | 0 | 1 | 0 | 0 | 0 | 0 | 0 | 0 | 0 | 0 | 0 | 0 | 0 | 0 | 0 | 0 | 0 | 0 | 0 | 0 | 0 | 0 | 0 | 0 | 0 | 0 | 0 | 0 | 0 | 0 | 0 | 0 | 0 | 0 | 0 | 0 |
| D03728 - Napsagatran (USAN) | 0 | 1 | 0 | 0 | 0 | 0 | 0 | 0 | 0 | 0 | 0 | 0 | 0 | 0 | 0 | 0 | 0 | 0 | 0 | 0 | 0 | 0 | 0 | 0 | 0 | 0 | 0 | 0 | 0 | 0 | 0 | 0 | 0 | 0 | 0 | 0 |
| D03766 - Diaplasinin (USAN) | 0 | 0 | 0 | 0 | 0 | 0 | 0 | 0 | 0 | 0 | 0 | 0 | 0 | 0 | 1 | 0 | 0 | 0 | 0 | 0 | 0 | 0 | 0 | 0 | 0 | 0 | 0 | 0 | 0 | 0 | 0 | 0 | 0 | 0 | 0 | 0 |
| D03793 - Ilomastat (USAN/INN) | 0 | 0 | 0 | 0 | 0 | 0 | 0 | 0 | 0 | 0 | 0 | 0 | 0 | 0 | 0 | 0 | 0 | 0 | 0 | 1 | 0 | 0 | 0 | 1 | 0 | 0 | 0 | 0 | 0 | 0 | 0 | 0 | 0 | 0 | 0 | 0 |
| D03795 - Marimastat (USAN/INN) | 0 | 0 | 0 | 0 | 0 | 0 | 0 | 0 | 0 | 0 | 0 | 0 | 0 | 0 | 0 | 0 | 0 | 0 | 0 | 1 | 0 | 0 | 0 | 1 | 0 | 0 | 0 | 0 | 0 | 0 | 0 | 0 | 0 | 0 | 0 | 0 |
| D03797 - Prinomastat (USAN/INN) | 0 | 0 | 0 | 0 | 0 | 0 | 0 | 0 | 0 | 0 | 0 | 0 | 0 | 0 | 0 | 0 | 0 | 0 | 0 | 1 | 0 | 0 | 0 | 1 | 0 | 0 | 0 | 1 | 0 | 0 | 0 | 0 | 0 | 0 | 0 | 0 |
| D03800 - Rebimastat (USAN/INN) | 0 | 0 | 0 | 0 | 0 | 0 | 0 | 0 | 0 | 0 | 0 | 0 | 0 | 0 | 0 | 0 | 0 | 0 | 0 | 1 | 0 | 0 | 0 | 1 | 0 | 0 | 0 | 0 | 0 | 0 | 0 | 0 | 0 | 0 | 0 | 0 |
| D03802 - Tanomastat (USAN/INN) | 0 | 0 | 0 | 0 | 0 | 0 | 0 | 0 | 0 | 0 | 0 | 0 | 0 | 0 | 0 | 0 | 0 | 0 | 0 | 1 | 0 | 0 | 0 | 1 | 0 | 0 | 0 | 0 | 0 | 0 | 0 | 0 | 0 | 0 | 0 | 0 |
| D03940 - Eculizumab (USAN/INN); Eculizumab (genetical recombination) (JAN); Soliris (TN) | 0 | 0 | 0 | 0 | 0 | 0 | 0 | 0 | 1 | 0 | 0 | 0 | 0 | 0 | 0 | 0 | 0 | 0 | 0 | 0 | 0 | 0 | 0 | 0 | 0 | 0 | 0 | 0 | 0 | 0 | 0 | 0 | 0 | 0 | 0 | 0 |
| D04029 - Razaxaban hydrochloride (USAN) | 0 | 0 | 0 | 1 | 0 | 0 | 0 | 0 | 0 | 0 | 0 | 0 | 0 | 0 | 0 | 0 | 0 | 0 | 0 | 0 | 0 | 0 | 0 | 0 | 0 | 0 | 0 | 0 | 0 | 0 | 0 | 0 | 0 | 0 | 0 | 0 |
| D04183 - Fidexaban (USAN/INN) | 0 | 0 | 0 | 1 | 0 | 0 | 0 | 0 | 0 | 0 | 0 | 0 | 0 | 0 | 0 | 0 | 0 | 0 | 0 | 0 | 0 | 0 | 0 | 0 | 0 | 0 | 0 | 0 | 0 | 0 | 0 | 0 | 0 | 0 | 0 | 0 |
| D05183 - Ulinastatin (JP18); Miraclid (TN) | 0 | 0 | 0 | 0 | 0 | 0 | 0 | 0 | 0 | 0 | 0 | 0 | 0 | 0 | 0 | 0 | 0 | 0 | 1 | 0 | 0 | 0 | 0 | 0 | 0 | 0 | 0 | 0 | 0 | 0 | 0 | 0 | 0 | 0 | 0 | 0 |
| D05917 - Stamulumab (USAN) | 1 | 0 | 0 | 0 | 0 | 0 | 0 | 0 | 0 | 0 | 0 | 0 | 0 | 0 | 0 | 0 | 0 | 0 | 0 | 0 | 0 | 0 | 0 | 0 | 0 | 0 | 0 | 0 | 0 | 0 | 0 | 0 | 0 | 0 | 0 | 0 |
| D06142 - Tifacogin (USAN/INN) | 0 | 0 | 0 | 1 | 0 | 0 | 0 | 0 | 0 | 0 | 0 | 0 | 0 | 0 | 0 | 0 | 0 | 0 | 0 | 0 | 0 | 0 | 0 | 0 | 0 | 0 | 0 | 0 | 0 | 0 | 0 | 0 | 0 | 0 | 0 | 0 |
| D06160 - Tiplasinin (USAN) | 0 | 0 | 0 | 0 | 0 | 0 | 0 | 0 | 0 | 0 | 0 | 0 | 0 | 0 | 1 | 0 | 0 | 0 | 0 | 0 | 0 | 0 | 0 | 0 | 0 | 0 | 0 | 0 | 0 | 0 | 0 | 0 | 0 | 0 | 0 | 0 |
| D06283 - Varespladib sodium (USAN) | 0 | 0 | 0 | 0 | 0 | 0 | 0 | 0 | 0 | 0 | 0 | 0 | 0 | 0 | 0 | 0 | 0 | 0 | 0 | 0 | 0 | 0 | 0 | 0 | 0 | 0 | 0 | 0 | 0 | 0 | 0 | 0 | 1 | 0 | 0 | 0 |

| | | | | | | | | | | | | | | | | | | | | | | | | | | | | | | | | | | | | |
|---|---|---|---|---|---|---|---|---|---|---|---|---|---|---|---|---|---|---|---|---|---|---|---|---|---|---|---|---|---|---|---|---|---|---|---|---|
| D06319 - Volociximab (USAN) | 0 | 0 | 0 | 0 | 0 | 0 | 0 | 0 | 0 | 0 | 0 | 0 | 0 | 0 | 0 | 0 | 1 | 0 | 0 | 0 | 1 | 0 | 0 | 0 | 0 | 0 | 0 | 0 | 0 | 0 | 0 | 0 | 0 | 0 | 0 | 0 |
| D06410 - Thrombomodulin alfa (USAN/INN); Thrombomodulin alfa (genetical recombination) (JAN) | 0 | 1 | 0 | 0 | 0 | 0 | 0 | 0 | 0 | 0 | 0 | 0 | 0 | 0 | 0 | 0 | 0 | 0 | 0 | 0 | 0 | 0 | 0 | 0 | 0 | 0 | 0 | 0 | 0 | 0 | 0 | 0 | 0 | 0 | 0 | 0 |
| D06456 - Cellulose; oxidized regenerated (USP); Surgicel absorbable hemostat (TN) | 0 | 0 | 0 | 0 | 0 | 0 | 0 | 0 | 0 | 0 | 0 | 0 | 0 | 0 | 0 | 0 | 0 | 0 | 0 | 0 | 0 | 0 | 0 | 0 | 0 | 0 | 0 | 0 | 0 | 0 | 1 | 0 | 0 | 0 | 0 | 0 |
| D06563 - Belagenpumatucel-L (USAN); Lucanix (TN) | 0 | 0 | 0 | 0 | 0 | 0 | 0 | 0 | 0 | 0 | 0 | 0 | 0 | 0 | 0 | 0 | 0 | 0 | 0 | 0 | 0 | 0 | 0 | 0 | 0 | 0 | 0 | 0 | 1 | 0 | 0 | 0 | 0 | 0 | 0 | 0 |
| D06634 - Relacatib (USAN/INN) | 0 | 0 | 0 | 0 | 0 | 0 | 0 | 0 | 0 | 0 | 0 | 0 | 0 | 0 | 0 | 0 | 0 | 0 | 0 | 0 | 0 | 0 | 0 | 0 | 0 | 0 | 1 | 0 | 0 | 0 | 0 | 0 | 0 | 0 | 0 | 0 |
| D06880 - Lepirudin (INN); Refludan (TN) | 0 | 1 | 0 | 0 | 0 | 0 | 0 | 0 | 0 | 0 | 0 | 0 | 0 | 0 | 0 | 0 | 0 | 0 | 0 | 0 | 0 | 0 | 0 | 0 | 0 | 0 | 0 | 0 | 0 | 0 | 0 | 0 | 0 | 0 | 0 | 0 |
| D07082 - Dabigatran etexilate mesylate (USAN); Dabigatran etexilate methanesulfonate (JAN); Pradaxa (TN) | 0 | 1 | 0 | 0 | 0 | 0 | 0 | 0 | 0 | 0 | 0 | 0 | 0 | 0 | 0 | 0 | 0 | 0 | 0 | 0 | 0 | 0 | 0 | 0 | 0 | 0 | 0 | 0 | 0 | 0 | 0 | 0 | 0 | 0 | 0 | 0 |
| D07086 - Rivaroxaban (JAN/USAN/INN); Xarelto (TN) | 0 | 0 | 0 | 1 | 0 | 0 | 0 | 0 | 0 | 0 | 0 | 0 | 0 | 0 | 0 | 0 | 0 | 0 | 0 | 0 | 0 | 0 | 0 | 0 | 0 | 0 | 0 | 0 | 0 | 0 | 0 | 0 | 0 | 0 | 0 | 0 |
| D07143 - Melagatran (INN) | 0 | 1 | 0 | 0 | 0 | 0 | 0 | 0 | 0 | 0 | 0 | 0 | 0 | 0 | 0 | 0 | 0 | 0 | 0 | 0 | 0 | 0 | 0 | 0 | 0 | 0 | 0 | 0 | 0 | 0 | 0 | 0 | 0 | 0 | 0 | 0 |
| D07144 - Dabigatran etexilate (USAN/INN); Pradaxa (TN) | 0 | 1 | 0 | 0 | 0 | 0 | 0 | 0 | 0 | 0 | 0 | 0 | 0 | 0 | 0 | 0 | 0 | 0 | 0 | 0 | 0 | 0 | 0 | 0 | 0 | 0 | 0 | 0 | 0 | 0 | 0 | 0 | 0 | 0 | 0 | 0 |
| D07568 - Aminomethylbenzoic acid; p-Aminomethylbenzoic acid; Gumbix (TN) | 0 | 0 | 0 | 0 | 1 | 0 | 0 | 0 | 0 | 0 | 0 | 0 | 0 | 0 | 0 | 0 | 0 | 0 | 0 | 0 | 0 | 0 | 0 | 0 | 0 | 0 | 0 | 0 | 0 | 0 | 0 | 0 | 0 | 0 | 0 | 0 |
| D08656 - Aleplasinin (USAN/INN) | 0 | 0 | 0 | 0 | 0 | 0 | 0 | 0 | 0 | 0 | 0 | 0 | 0 | 0 | 1 | 0 | 0 | 0 | 0 | 0 | 0 | 0 | 0 | 0 | 0 | 0 | 0 | 0 | 0 | 0 | 0 | 0 | 0 | 0 | 0 | 0 |
| D08858 - Antithrombin alfa (INN/USAN); Atryn (TN) | 0 | 0 | 0 | 1 | 0 | 0 | 0 | 0 | 0 | 0 | 0 | 0 | 0 | 0 | 0 | 0 | 0 | 0 | 0 | 0 | 0 | 0 | 0 | 0 | 0 | 0 | 0 | 0 | 0 | 0 | 0 | 0 | 0 | 0 | 0 | 0 |
| D08859 - Apratastat (USAN/INN) | 0 | 0 | 0 | 0 | 0 | 0 | 0 | 0 | 0 | 0 | 0 | 0 | 0 | 0 | 0 | 0 | 0 | 0 | 0 | 0 | 0 | 0 | 0 | 0 | 0 | 0 | 0 | 1 | 0 | 0 | 0 | 0 | 0 | 0 | 0 | 0 |
| D08873 - Betrixaban (USAN); Bevyxxa (TN) | 0 | 0 | 0 | 1 | 0 | 0 | 0 | 0 | 0 | 0 | 0 | 0 | 0 | 0 | 0 | 0 | 0 | 0 | 0 | 0 | 0 | 0 | 0 | 0 | 0 | 0 | 0 | 0 | 0 | 0 | 0 | 0 | 0 | 0 | 0 | 0 |
| D08913 - Eribaxaban (USAN) | 0 | 0 | 0 | 1 | 0 | 0 | 0 | 0 | 0 | 0 | 0 | 0 | 0 | 0 | 0 | 0 | 0 | 0 | 0 | 0 | 0 | 0 | 0 | 0 | 0 | 0 | 0 | 0 | 0 | 0 | 0 | 0 | 0 | 0 | 0 | 0 |
| D08946 - Mipomersen sodium (USAN); Kynamro (TN) | 0 | 0 | 0 | 0 | 0 | 0 | 0 | 0 | 0 | 0 | 0 | 0 | 0 | 1 | 0 | 0 | 0 | 0 | 0 | 0 | 0 | 0 | 0 | 0 | 0 | 0 | 0 | 0 | 0 | 0 | 0 | 0 | 0 | 0 | 0 | 0 |
| D08955 - Odanacatib (JAN/USAN) | 0 | 0 | 0 | 0 | 0 | 0 | 0 | 0 | 0 | 0 | 0 | 0 | 0 | 0 | 0 | 0 | 0 | 0 | 0 | 0 | 0 | 0 | 0 | 0 | 0 | 0 | 1 | 0 | 0 | 0 | 0 | 0 | 0 | 0 | 0 | 0 |
| D09342 - Etaracizumab (USAN/INN); Abegrin (TN) | 0 | 0 | 0 | 0 | 0 | 0 | 0 | 0 | 0 | 0 | 0 | 0 | 0 | 0 | 0 | 0 | 0 | 1 | 0 | 0 | 0 | 0 | 0 | 0 | 0 | 0 | 0 | 0 | 0 | 0 | 0 | 0 | 0 | 0 | 0 | 0 |
| D09403 - Custirsen sodium (USAN) | 0 | 0 | 0 | 0 | 0 | 0 | 0 | 0 | 0 | 0 | 0 | 0 | 0 | 0 | 0 | 0 | 0 | 0 | 0 | 0 | 0 | 0 | 1 | 0 | 0 | 0 | 0 | 0 | 0 | 0 | 0 | 0 | 0 | 0 | 0 | 0 |
| D09546 - Edoxaban tosilate hydrate (JAN); Edoxaban tosylate monohydrate; Lixiana (TN); Savaysa (TN) | 0 | 0 | 0 | 1 | 0 | 0 | 0 | 0 | 0 | 0 | 0 | 0 | 0 | 0 | 0 | 0 | 0 | 0 | 0 | 0 | 0 | 0 | 0 | 0 | 0 | 0 | 0 | 0 | 0 | 0 | 0 | 0 | 0 | 0 | 0 | 0 |
| D09620 - Fresolimumab (USAN/INN) | 0 | 0 | 0 | 0 | 0 | 0 | 0 | 0 | 0 | 0 | 0 | 0 | 0 | 0 | 0 | 0 | 0 | 0 | 0 | 0 | 0 | 1 | 0 | 0 | 0 | 0 | 0 | 0 | 1 | 0 | 0 | 0 | 0 | 0 | 0 | 0 |
| D09631 - Intetumumab (USAN/INN) | 0 | 0 | 0 | 0 | 0 | 0 | 0 | 0 | 0 | 0 | 0 | 0 | 0 | 0 | 0 | 0 | 0 | 1 | 0 | 0 | 0 | 0 | 0 | 0 | 0 | 0 | 0 | 0 | 0 | 0 | 0 | 0 | 0 | 0 | 0 | 0 |
| D09673 - Tafamidis (JAN/USAN/INN); Vyndamax (TN); Vynmac (TN) | 0 | 0 | 0 | 0 | 0 | 0 | 0 | 0 | 0 | 0 | 0 | 0 | 1 | 0 | 0 | 0 | 0 | 0 | 0 | 0 | 0 | 0 | 0 | 0 | 0 | 0 | 0 | 0 | 0 | 0 | 0 | 0 | 0 | 0 | 0 | 0 |
| D09674 - Tafamidis meglumine (JAN/USAN); Vyndaqel (TN) | 0 | 0 | 0 | 0 | 0 | 0 | 0 | 0 | 0 | 0 | 0 | 0 | 1 | 0 | 0 | 0 | 0 | 0 | 0 | 0 | 0 | 0 | 0 | 0 | 0 | 0 | 0 | 0 | 0 | 0 | 0 | 0 | 0 | 0 | 0 | 0 |
| D09707 - Dabigatran (USAN/INN) | 0 | 1 | 0 | 0 | 0 | 0 | 0 | 0 | 0 | 0 | 0 | 0 | 0 | 0 | 0 | 0 | 0 | 0 | 0 | 0 | 0 | 0 | 0 | 0 | 0 | 0 | 0 | 0 | 0 | 0 | 0 | 0 | 0 | 0 | 0 | 0 |
| D09710 - Edoxaban (USAN/INN) | 0 | 0 | 0 | 1 | 0 | 0 | 0 | 0 | 0 | 0 | 0 | 0 | 0 | 0 | 0 | 0 | 0 | 0 | 0 | 0 | 0 | 0 | 0 | 0 | 0 | 0 | 0 | 0 | 0 | 0 | 0 | 0 | 0 | 0 | 0 | 0 |
| D09799 - Carotegrast methyl (JAN); Carogra (TN) | 0 | 0 | 0 | 0 | 0 | 0 | 0 | 0 | 0 | 0 | 0 | 0 | 0 | 0 | 0 | 0 | 1 | 0 | 0 | 0 | 0 | 0 | 0 | 0 | 0 | 0 | 0 | 0 | 0 | 0 | 0 | 0 | 0 | 0 | 0 | 0 |
| D09817 - Darexaban maleate (JAN); YM150 | 0 | 0 | 0 | 1 | 0 | 0 | 0 | 0 | 0 | 0 | 0 | 0 | 0 | 0 | 0 | 0 | 0 | 0 | 0 | 0 | 0 | 0 | 0 | 0 | 0 | 0 | 0 | 0 | 0 | 0 | 0 | 0 | 0 | 0 | 0 | 0 |
| D09903 - Fluciclatide F 18 (USAN/INN) | 0 | 0 | 0 | 0 | 0 | 0 | 0 | 0 | 0 | 0 | 0 | 0 | 0 | 0 | 0 | 0 | 0 | 1 | 0 | 0 | 0 | 0 | 0 | 0 | 0 | 0 | 0 | 0 | 0 | 0 | 0 | 0 | 0 | 0 | 0 | 0 |
| D09923 - Letaxaban (USAN/INN) | 0 | 0 | 0 | 1 | 0 | 0 | 0 | 0 | 0 | 0 | 0 | 0 | 0 | 0 | 0 | 0 | 0 | 0 | 0 | 0 | 0 | 0 | 0 | 0 | 0 | 0 | 0 | 0 | 0 | 0 | 0 | 0 | 0 | 0 | 0 | 0 |
| D09929 - Maraciclatide (USAN/INN) | 0 | 0 | 0 | 0 | 0 | 0 | 0 | 0 | 0 | 0 | 0 | 0 | 0 | 0 | 0 | 0 | 0 | 1 | 0 | 0 | 0 | 0 | 0 | 0 | 0 | 0 | 0 | 0 | 0 | 0 | 0 | 0 | 0 | 0 | 0 | 0 |
| D10023 - Pexelizumab (USAN/INN) | 0 | 0 | 0 | 0 | 0 | 0 | 0 | 0 | 1 | 0 | 0 | 0 | 0 | 0 | 0 | 0 | 0 | 0 | 0 | 0 | 0 | 0 | 0 | 0 | 0 | 0 | 0 | 0 | 0 | 0 | 0 | 0 | 0 | 0 | 0 | 0 |
| D10028 - Valategrast hydrochloride (USAN) | 0 | 0 | 0 | 0 | 0 | 0 | 0 | 0 | 0 | 0 | 0 | 0 | 0 | 0 | 0 | 0 | 1 | 0 | 0 | 0 | 0 | 0 | 0 | 0 | 0 | 0 | 0 | 0 | 0 | 0 | 0 | 0 | 0 | 0 | 0 | 0 |
| D10150 - Pegnivacogin sodium (USAN) | 0 | 0 | 1 | 0 | 0 | 0 | 0 | 0 | 0 | 0 | 0 | 0 | 0 | 0 | 0 | 0 | 0 | 0 | 0 | 0 | 0 | 0 | 0 | 0 | 0 | 0 | 0 | 0 | 0 | 0 | 0 | 0 | 0 | 0 | 0 | 0 |
| D10203 - Trabedersen (INN/BAN) | 0 | 0 | 0 | 0 | 0 | 0 | 0 | 0 | 0 | 0 | 0 | 0 | 0 | 0 | 0 | 0 | 0 | 0 | 0 | 0 | 0 | 0 | 0 | 0 | 0 | 0 | 0 | 0 | 1 | 0 | 0 | 0 | 0 | 0 | 0 | 0 |
| D10347 - Ramatercept (USAN); Ace-031 | 1 | 0 | 0 | 0 | 0 | 0 | 0 | 0 | 0 | 0 | 0 | 0 | 0 | 0 | 0 | 0 | 0 | 0 | 0 | 0 | 0 | 0 | 0 | 0 | 0 | 0 | 0 | 0 | 0 | 0 | 0 | 0 | 0 | 0 | 0 | 0 |

| | | | | | | | | | | | | | | | | | | | | | | | | | | | | | | | | | | | | |
|---|---|---|---|---|---|---|---|---|---|---|---|---|---|---|---|---|---|---|---|---|---|---|---|---|---|---|---|---|---|---|---|---|---|---|---|---|
| D10471 - Idraparinux sodium (USAN) | 0 | 0 | 0 | 1 | 0 | 0 | 0 | 0 | 0 | 0 | 0 | 0 | 0 | 0 | 0 | 0 | 0 | 0 | 0 | 0 | 0 | 0 | 0 | 0 | 0 | 0 | 0 | 0 | 0 | 0 | 0 | 0 | 0 | 0 | 0 | 0 |
| D10484 - Pegnivacogin (USAN) | 0 | 0 | 1 | 0 | 0 | 0 | 0 | 0 | 0 | 0 | 0 | 0 | 0 | 0 | 0 | 0 | 0 | 0 | 0 | 0 | 0 | 0 | 0 | 0 | 0 | 0 | 0 | 0 | 0 | 0 | 0 | 0 | 0 | 0 | 0 | 0 |
| D10595 - Asfotase alfa (USAN/INN) | 0 | 0 | 0 | 0 | 0 | 0 | 0 | 0 | 0 | 0 | 0 | 0 | 0 | 0 | 0 | 0 | 0 | 1 | 0 | 0 | 0 | 0 | 0 | 0 | 0 | 0 | 0 | 0 | 0 | 0 | 0 | 0 | 0 | 0 | 0 | 0 |
| D10794 - Patisiran sodium (JAN); Onpattro (TN) | 0 | 0 | 0 | 0 | 0 | 0 | 0 | 0 | 0 | 0 | 0 | 0 | 1 | 0 | 0 | 0 | 0 | 0 | 0 | 0 | 0 | 0 | 0 | 0 | 0 | 0 | 0 | 0 | 0 | 0 | 0 | 0 | 0 | 0 | 0 | 0 |
| D10821 - Emicizumab (USAN/INN) | 0 | 0 | 1 | 1 | 0 | 0 | 0 | 0 | 0 | 0 | 0 | 0 | 0 | 0 | 0 | 0 | 0 | 0 | 0 | 0 | 0 | 0 | 0 | 0 | 0 | 0 | 0 | 0 | 0 | 0 | 0 | 0 | 0 | 0 | 0 | 0 |
| D10913 - Burosumab (USAN/INN) | 0 | 0 | 0 | 0 | 0 | 0 | 0 | 0 | 0 | 0 | 0 | 0 | 0 | 0 | 0 | 0 | 0 | 0 | 0 | 0 | 0 | 0 | 0 | 0 | 0 | 0 | 0 | 0 | 0 | 0 | 0 | 0 | 0 | 0 | 1 | 0 |
| D10940 - Inotersen (USAN/INN) | 0 | 0 | 0 | 0 | 0 | 0 | 0 | 0 | 0 | 0 | 0 | 0 | 1 | 0 | 0 | 0 | 0 | 0 | 0 | 0 | 0 | 0 | 0 | 0 | 0 | 0 | 0 | 0 | 0 | 0 | 0 | 0 | 0 | 0 | 0 | 0 |
| D10941 - Inotersen sodium (USAN); Tegsedi (TN) | 0 | 0 | 0 | 0 | 0 | 0 | 0 | 0 | 0 | 0 | 0 | 0 | 1 | 0 | 0 | 0 | 0 | 0 | 0 | 0 | 0 | 0 | 0 | 0 | 0 | 0 | 0 | 0 | 0 | 0 | 0 | 0 | 0 | 0 | 0 | 0 |
| D10969 - Pamrevlumab (USAN/INN) | 0 | 0 | 0 | 0 | 0 | 0 | 0 | 0 | 0 | 0 | 0 | 0 | 0 | 0 | 0 | 0 | 0 | 0 | 0 | 0 | 0 | 0 | 0 | 0 | 1 | 0 | 0 | 0 | 0 | 0 | 0 | 0 | 0 | 0 | 0 | 0 |
| D11029 - Andexanet alfa (USAN/INN) | 0 | 0 | 0 | 1 | 0 | 0 | 0 | 0 | 0 | 0 | 0 | 0 | 0 | 0 | 0 | 0 | 0 | 0 | 0 | 0 | 0 | 0 | 0 | 0 | 0 | 0 | 0 | 0 | 0 | 0 | 0 | 0 | 0 | 0 | 0 | 0 |
| D11032 - Plitidepsin (INN) | 0 | 0 | 0 | 0 | 0 | 0 | 0 | 0 | 0 | 0 | 0 | 0 | 0 | 0 | 0 | 0 | 0 | 0 | 0 | 0 | 0 | 0 | 0 | 0 | 0 | 0 | 0 | 0 | 1 | 0 | 0 | 0 | 0 | 0 | 0 | 0 |
| D11047 - Domagrozumab (USAN/INN) | 1 | 0 | 0 | 0 | 0 | 0 | 0 | 0 | 0 | 0 | 0 | 0 | 0 | 0 | 0 | 0 | 0 | 0 | 0 | 0 | 0 | 0 | 0 | 0 | 0 | 0 | 0 | 0 | 0 | 0 | 0 | 0 | 0 | 0 | 0 | 0 |
| D11054 - Ravulizumab (USAN) | 0 | 0 | 0 | 0 | 0 | 0 | 0 | 0 | 1 | 0 | 0 | 0 | 0 | 0 | 0 | 0 | 0 | 0 | 0 | 0 | 0 | 0 | 0 | 0 | 0 | 0 | 0 | 0 | 0 | 0 | 0 | 0 | 0 | 0 | 0 | 0 |
| D11116 - Patisiran (USAN/INN) | 0 | 0 | 0 | 0 | 0 | 0 | 0 | 0 | 0 | 0 | 0 | 0 | 1 | 0 | 0 | 0 | 0 | 0 | 0 | 0 | 0 | 0 | 0 | 0 | 0 | 0 | 0 | 0 | 0 | 0 | 0 | 0 | 0 | 0 | 0 | 0 |
| D11235 - Landogrozumab (USAN/INN) | 1 | 0 | 0 | 0 | 0 | 0 | 0 | 0 | 0 | 0 | 0 | 0 | 0 | 0 | 0 | 0 | 0 | 0 | 0 | 0 | 0 | 0 | 0 | 0 | 0 | 0 | 0 | 0 | 0 | 0 | 0 | 0 | 0 | 0 | 0 | 0 |
| D11242 - Trevogrumab (USAN/INN) | 1 | 0 | 0 | 0 | 0 | 0 | 0 | 0 | 0 | 0 | 0 | 0 | 0 | 0 | 0 | 0 | 0 | 0 | 0 | 0 | 0 | 0 | 0 | 0 | 0 | 0 | 0 | 0 | 0 | 0 | 0 | 0 | 0 | 0 | 0 | 0 |
| D11262 - Andecaliximab (USAN/INN) | 0 | 0 | 0 | 0 | 0 | 0 | 0 | 0 | 0 | 0 | 0 | 0 | 0 | 0 | 0 | 0 | 0 | 0 | 0 | 0 | 0 | 0 | 1 | 0 | 0 | 0 | 0 | 0 | 0 | 0 | 0 | 0 | 0 | 0 | 0 | 0 |
| D11265 - Taldefgrobep alfa (USAN); Talditercept alfa (USAN) | 1 | 0 | 0 | 0 | 0 | 0 | 0 | 0 | 0 | 0 | 0 | 0 | 0 | 0 | 0 | 0 | 0 | 0 | 0 | 0 | 0 | 0 | 0 | 0 | 0 | 0 | 0 | 0 | 0 | 0 | 0 | 0 | 0 | 0 | 0 | 0 |
| D11280 - Bivalirudin trifluoroacetate (USAN); Angiomax (TN) | 0 | 1 | 0 | 0 | 0 | 0 | 0 | 0 | 0 | 0 | 0 | 0 | 0 | 0 | 0 | 0 | 0 | 0 | 0 | 0 | 0 | 0 | 0 | 0 | 0 | 0 | 0 | 0 | 0 | 0 | 0 | 0 | 0 | 0 | 0 | 0 |
| D11331 - Olendalizumab (USAN); Lendalizumab (USAN) | 0 | 0 | 0 | 0 | 0 | 0 | 0 | 0 | 1 | 0 | 0 | 0 | 0 | 0 | 0 | 0 | 0 | 0 | 0 | 0 | 0 | 0 | 0 | 0 | 0 | 0 | 0 | 0 | 0 | 0 | 0 | 0 | 0 | 0 | 0 | 0 |
| D11369 - Rosmantuzumab (USAN) | 0 | 0 | 0 | 0 | 0 | 0 | 0 | 0 | 0 | 0 | 0 | 0 | 0 | 0 | 0 | 0 | 0 | 0 | 0 | 0 | 0 | 0 | 0 | 0 | 0 | 0 | 0 | 0 | 0 | 0 | 0 | 0 | 0 | 1 | 0 | 0 |
| D11464 - Miridesap (USAN/INN) | 0 | 0 | 0 | 0 | 0 | 0 | 0 | 0 | 0 | 0 | 1 | 0 | 0 | 0 | 0 | 0 | 0 | 0 | 0 | 0 | 0 | 0 | 0 | 0 | 0 | 0 | 0 | 0 | 0 | 0 | 0 | 0 | 0 | 0 | 0 | 0 |
| D11471 - Dezamizumab (USAN/INN) | 0 | 0 | 0 | 0 | 0 | 0 | 0 | 0 | 0 | 0 | 1 | 0 | 0 | 0 | 0 | 0 | 0 | 0 | 0 | 0 | 0 | 0 | 0 | 0 | 0 | 0 | 0 | 0 | 0 | 0 | 0 | 0 | 0 | 0 | 0 | 0 |
| D11473 - Nomacopan (USAN) | 0 | 0 | 0 | 0 | 0 | 0 | 0 | 0 | 1 | 0 | 0 | 0 | 0 | 0 | 0 | 0 | 0 | 0 | 0 | 0 | 0 | 0 | 0 | 0 | 0 | 0 | 0 | 0 | 0 | 0 | 0 | 0 | 0 | 0 | 0 | 0 |
| D11477 - Pozelimab (USAN) | 0 | 0 | 0 | 0 | 0 | 0 | 0 | 0 | 1 | 0 | 0 | 0 | 0 | 0 | 0 | 0 | 0 | 0 | 0 | 0 | 0 | 0 | 0 | 0 | 0 | 0 | 0 | 0 | 0 | 0 | 0 | 0 | 0 | 0 | 0 | 0 |
| D11568 - Edoxaban tosylate (USAN) | 0 | 0 | 0 | 1 | 0 | 0 | 0 | 0 | 0 | 0 | 0 | 0 | 0 | 0 | 0 | 0 | 0 | 0 | 0 | 0 | 0 | 0 | 0 | 0 | 0 | 0 | 0 | 0 | 0 | 0 | 0 | 0 | 0 | 0 | 0 | 0 |
| D11613 - Pegcetacoplan (JAN/USAN/INN); Empaveli (TN) | 0 | 0 | 0 | 0 | 0 | 0 | 0 | 1 | 0 | 0 | 0 | 0 | 0 | 0 | 0 | 0 | 0 | 0 | 0 | 0 | 0 | 0 | 0 | 0 | 0 | 0 | 0 | 0 | 0 | 0 | 0 | 0 | 0 | 0 | 0 | 0 |
| D11696 - Crovalimab (USAN/INN) | 0 | 0 | 0 | 0 | 0 | 0 | 0 | 0 | 1 | 0 | 0 | 0 | 0 | 0 | 0 | 0 | 0 | 0 | 0 | 0 | 0 | 0 | 0 | 0 | 0 | 0 | 0 | 0 | 0 | 0 | 0 | 0 | 0 | 0 | 0 | 0 |
| D11748 - Avacincaptad pegol sodium (USAN) | 0 | 0 | 0 | 0 | 0 | 0 | 0 | 0 | 1 | 0 | 0 | 0 | 0 | 0 | 0 | 0 | 0 | 0 | 0 | 0 | 0 | 0 | 0 | 0 | 0 | 0 | 0 | 0 | 0 | 0 | 0 | 0 | 0 | 0 | 0 | 0 |
| D11770 - Vonapanitase (USAN/INN) | 0 | 0 | 0 | 0 | 0 | 0 | 0 | 0 | 0 | 0 | 0 | 0 | 0 | 0 | 0 | 0 | 0 | 0 | 0 | 0 | 0 | 0 | 0 | 0 | 0 | 0 | 0 | 0 | 0 | 0 | 0 | 0 | 0 | 0 | 0 | 1 |
| D11797 - Apitegromab (USAN) | 1 | 0 | 0 | 0 | 0 | 0 | 0 | 0 | 0 | 0 | 0 | 0 | 0 | 0 | 0 | 0 | 0 | 0 | 0 | 0 | 0 | 0 | 0 | 0 | 0 | 0 | 0 | 0 | 0 | 0 | 0 | 0 | 0 | 0 | 0 | 0 |
| D11838 - Vilobelimab (USAN) | 0 | 0 | 0 | 0 | 0 | 0 | 0 | 0 | 1 | 0 | 0 | 0 | 0 | 0 | 0 | 0 | 0 | 0 | 0 | 0 | 0 | 0 | 0 | 0 | 0 | 0 | 0 | 0 | 0 | 0 | 0 | 0 | 0 | 0 | 0 | 0 |
| D11916 - Vutrisiran sodium (JAN); Amvuttra (TN) | 0 | 0 | 0 | 0 | 0 | 0 | 0 | 0 | 0 | 0 | 0 | 0 | 1 | 0 | 0 | 0 | 0 | 0 | 0 | 0 | 0 | 0 | 0 | 0 | 0 | 0 | 0 | 0 | 0 | 0 | 0 | 0 | 0 | 0 | 0 | 0 |
| D11917 - Vutrisiran (USAN/INN) | 0 | 0 | 0 | 0 | 0 | 0 | 0 | 0 | 0 | 0 | 0 | 0 | 1 | 0 | 0 | 0 | 0 | 0 | 0 | 0 | 0 | 0 | 0 | 0 | 0 | 0 | 0 | 0 | 0 | 0 | 0 | 0 | 0 | 0 | 0 | 0 |
| D11965 - alpha 1-Antitrypsin; Human alpha 1-Proteinase inibitor; freeze-dried concentrated; Lynspad (TN) | 0 | 0 | 0 | 0 | 0 | 0 | 1 | 0 | 0 | 0 | 0 | 0 | 0 | 0 | 0 | 0 | 0 | 0 | 0 | 0 | 0 | 0 | 0 | 0 | 0 | 0 | 0 | 0 | 0 | 0 | 0 | 0 | 0 | 0 | 0 | 0 |
| D11972 - Acoramidis (USAN) | 0 | 0 | 0 | 0 | 0 | 0 | 0 | 0 | 0 | 0 | 0 | 0 | 1 | 0 | 0 | 0 | 0 | 0 | 0 | 0 | 0 | 0 | 0 | 0 | 0 | 0 | 0 | 0 | 0 | 0 | 0 | 0 | 0 | 0 | 0 | 0 |
| D11973 - Acoramidis hydrochloride (JAN/USAN) | 0 | 0 | 0 | 0 | 0 | 0 | 0 | 0 | 0 | 0 | 0 | 0 | 1 | 0 | 0 | 0 | 0 | 0 | 0 | 0 | 0 | 0 | 0 | 0 | 0 | 0 | 0 | 0 | 0 | 0 | 0 | 0 | 0 | 0 | 0 | 0 |

| | | | | | | | | | | | | | | | | | | | | | | | | | | | | | | | | | | | | |
|---|---|---|---|---|---|---|---|---|---|---|---|---|---|---|---|---|---|---|---|---|---|---|---|---|---|---|---|---|---|---|---|---|---|---|---|---|
| D12003 - Simufilam (USAN) | 0 | 0 | 0 | 0 | 0 | 0 | 0 | 0 | 0 | 0 | 0 | 0 | 0 | 0 | 0 | 0 | 0 | 0 | 0 | 0 | 0 | 0 | 0 | 0 | 1 | 0 | 0 | 0 | 0 | 0 | 0 | 0 | 0 | 0 | 0 | 0 |
| D12004 - Simufilam hydrochloride (USAN) | 0 | 0 | 0 | 0 | 0 | 0 | 0 | 0 | 0 | 0 | 0 | 0 | 0 | 0 | 0 | 0 | 0 | 0 | 0 | 0 | 0 | 0 | 0 | 0 | 1 | 0 | 0 | 0 | 0 | 0 | 0 | 0 | 0 | 0 | 0 | 0 |
| D12149 - Plasminogen; Glu-plasminogen; Plasminogen; human-tvmh; Ryplazim (TN) | 0 | 0 | 0 | 0 | 1 | 0 | 0 | 0 | 0 | 0 | 0 | 0 | 0 | 0 | 0 | 0 | 0 | 0 | 0 | 0 | 0 | 0 | 0 | 0 | 0 | 0 | 0 | 0 | 0 | 0 | 0 | 0 | 0 | 0 | 0 | 0 |
| D12208 - Gefurulimab (USAN) | 0 | 0 | 0 | 0 | 0 | 0 | 0 | 0 | 1 | 0 | 0 | 0 | 0 | 0 | 0 | 0 | 0 | 0 | 0 | 0 | 0 | 0 | 0 | 0 | 0 | 0 | 0 | 0 | 0 | 0 | 0 | 0 | 0 | 0 | 0 | 0 |
| D12251 - Iptacopan (USAN/INN) | 0 | 0 | 0 | 0 | 0 | 1 | 0 | 0 | 0 | 0 | 0 | 0 | 0 | 0 | 0 | 0 | 0 | 0 | 0 | 0 | 0 | 0 | 0 | 0 | 0 | 0 | 0 | 0 | 0 | 0 | 0 | 0 | 0 | 0 | 0 | 0 |
| D12252 - Iptacopan hydrochloride (USAN); Iptacopan hydrochloride monohydrate | 0 | 0 | 0 | 0 | 0 | 1 | 0 | 0 | 0 | 0 | 0 | 0 | 0 | 0 | 0 | 0 | 0 | 0 | 0 | 0 | 0 | 0 | 0 | 0 | 0 | 0 | 0 | 0 | 0 | 0 | 0 | 0 | 0 | 0 | 0 | 0 |
| D12351 - Tinlarebant (USAN) | 0 | 0 | 0 | 0 | 0 | 0 | 0 | 0 | 0 | 0 | 0 | 1 | 0 | 0 | 0 | 0 | 0 | 0 | 0 | 0 | 0 | 0 | 0 | 0 | 0 | 0 | 0 | 0 | 0 | 0 | 0 | 0 | 0 | 0 | 0 | 0 |
| D12356 - Zilucoplan sodium (JAN) | 0 | 0 | 0 | 0 | 0 | 0 | 0 | 0 | 1 | 0 | 0 | 0 | 0 | 0 | 0 | 0 | 0 | 0 | 0 | 0 | 0 | 0 | 0 | 0 | 0 | 0 | 0 | 0 | 0 | 0 | 0 | 0 | 0 | 0 | 0 | 0 |
| D12357 - Zilucoplan (USAN/INN) | 0 | 0 | 0 | 0 | 0 | 0 | 0 | 0 | 1 | 0 | 0 | 0 | 0 | 0 | 0 | 0 | 0 | 0 | 0 | 0 | 0 | 0 | 0 | 0 | 0 | 0 | 0 | 0 | 0 | 0 | 0 | 0 | 0 | 0 | 0 | 0 |
| D12379 - Galegenimab (USAN) | 0 | 0 | 0 | 0 | 0 | 0 | 0 | 0 | 0 | 0 | 0 | 0 | 0 | 0 | 0 | 0 | 0 | 0 | 0 | 0 | 0 | 0 | 0 | 0 | 0 | 0 | 0 | 0 | 0 | 0 | 0 | 1 | 0 | 0 | 0 | 0 |
| D12402 - Zinpentraxin alfa (USAN) | 0 | 0 | 0 | 0 | 0 | 0 | 0 | 0 | 0 | 0 | 1 | 0 | 0 | 0 | 0 | 0 | 0 | 0 | 0 | 0 | 0 | 0 | 0 | 0 | 0 | 0 | 0 | 0 | 0 | 0 | 0 | 0 | 0 | 0 | 0 | 0 |
